# Dynamic convergence of neurodevelopmental disorder risk genes across neurodevelopment

**DOI:** 10.1101/2024.08.23.609190

**Authors:** Meilin Fernandez Garcia, Kayla Retallick-Townsley, April Pruitt, Elizabeth Davidson, Novin Balafkan, Jonathan Warrell, Tzu-Chieh Huang, Alfred Kibowen, Zhiyuan Chu, Yi Dai, Sarah E. Fitzpatrick, Ran Meng, Annabel Sen, Sophie Cohen, Olivia Livoti, Suha Khan, Charlotte Becker, Andre Luiz Teles e Silva, Jenny Liu, Grace Dossou, Jen Cheung, Susanna Liu, Sadaf Ghorbani, P.J. Michael Deans, Marisa DeCiucis, Prashant Emani, Huanyao Gao, Hongying Shen, Mark Gerstein, Zuoheng Wang, Laura M. Huckins, Ellen J. Hoffman, Kristen Brennand

**Author notes:** These authors contributed equally (Meilin Fernandez Garcia, Kayla Retallick-Townsley, April Pruitt, Elizabeth Davidson, Novin Balafkan).

## Abstract

Over three hundred and seventy-three risk genes, broadly enriched for roles in neuronal communication and gene expression regulation, underlie risk for autism spectrum disorder (ASD) and developmental delay (DD). Functional genomic studies of subsets of these genes consistently indicate a convergent role in neurogenesis, but how these diverse risk genes converge on a smaller number of biological pathways in mature neurons is unclear. To uncover shared downstream impacts between neurodevelopmental disorder (NDD) risk genes, here we apply a pooled CRISPR approach to contrast the transcriptomic impacts of targeting 29 NDD loss-of-function genes across human induced pluripotent stem cell (hiPSC)-derived neural progenitor cells, glutamatergic neurons, and GABAergic neurons. Points of convergence vary between the cell types of the brain and are greatest in mature glutamatergic neurons, where they broadly target not just synaptic and epigenetic, but unexpectedly, mitochondrial biology. The strongest convergent networks occur between NDD genes with common co-expression patterns in the post-mortem brain, biological annotations, and clinical associations, suggesting that convergence may one-day inform patient stratification and treatment. Towards this, ten out of eleven drugs tested that were predicted to reverse convergent signatures in human cells and/or arousal and sensory processing behaviors in zebrafish ameliorated at least one behavioral phenotype *in vivo*. Altogether, robust convergence in post-mitotic neurons represents a clinically actionable therapeutic window.

## INTRODUCTION

Autism spectrum disorder (ASD) and related developmental delay (DD) are highly heritable^1^. The aggregate impact of common variants of small effect reflects most genetic risk^2^, but in as many as a quarter of cases, potentially damaging rare inherited and *de novo* mutations in risk genes are detected^3^. There is significant overlap between those genes affecting ASD^4^ and those more broadly affecting developmental^5,6^ and psychiatric^7,8^ disorders. Altogether, neurodevelopmental disorder (NDD) risk genes are typically expressed during cortical development^9^, particularly the excitatory and inhibitory lineages^4^, and broadly split between two functional classes: neuronal communication (e.g., synaptic function) and gene expression regulation (e.g., chromatin regulators and transcription factors)^4,10–15^. Over half of NDD genes have roles in gene expression regulation^4^, sharing substantial overlap in genomic binding sites in the brain^16^, and with targets enriched for NDD risk genes^17–20^. Yet, evidence to support the parsimonious explanation that regulatory NDD genes preferentially target synaptic NDD genes, is lacking^4^. It remains unclear how disrupting NDD genes with distinct functions yields similar outcomes.

Many NDD genes seem to have broad roles outside their annotated function; for example, some chromatin regulators (e.g., *CHD8, CHD2,* and *POGZ*) localize to microtubules in the centrosome^21^, mitotic spindle^22^, and cilia^23,24^, suggesting the possibility that they function directly in neurogenesis and/or synaptic biology. Indeed, both regulatory and synaptic genes impact proliferation and patterning of progenitors (e.g., *ARID1B*^25,26^, *CHD8*^27,28^, *NRXN1*^29,30^, *SYNGAP1*^31^), excitatory transmission by glutamatergic neurons (e.g., *CHD8*^32,33^, *NRXN1*^34^, *SHANK3*^35^, *SYNGAP1*^36^), and inhibitory transmission by GABAergic neurons (e.g., *ARID1B*^37^, *CHD8*^32^, *NRXN1*^38^, *SHANK3*^39^). Do overlapping downstream impacts explain how heterogeneous gene mutations result in similar neuronal phenotypes and clinical outcomes^40^?

Many have proposed that diverse ASD genes are convergent^41–43^. Indeed, NDD genes are co-expressed in the brain^44–46^, suggesting that they are regulated together and involved in related biological processes, and result in highly interconnected protein-protein interactomes^47–50^, indicating functional relationships between NDD proteins. Even as the number of NDD genes grows, risk genes continue to converge on a finite number of biological pathways, developmental stages, brain regions and cell types^41^. Disentangling these complex etiologies remains an outstanding challenge.

Excitatory-inhibitory (E:I) imbalance is widely believed to underlie NDD^51–53^, whether arising from altered proportions of neuronal lineage cell types in the developing brain or synaptic deficits in glutamatergic or GABAergic neurons. Indeed, knockdown of subsets of NDD genes in human neural progenitor cells (NPCs)^22,54,55^, cerebral organoids^27,56,57^, and developing mouse^58^, tadpole^59^ and zebrafish^60^ brains reveal overlapping impacts on neurogenesis. Despite synaptic dysfunction being a hallmark of NDD, the extent to which downstream impacts of NDD genes also converge in mature neurons is largely unknown.

Given emerging evidence that epigenetic NDD genes have diverse and interconnected roles^21–24^, we tested the hypothesis that the nature of convergence is dynamic, influenced by developmental and cell-type contexts. We report a pooled CRISPR-knockout (KO) strategy targeting loss-of-function (LoF) mutations to 29 NDD genes, most with roles in chromatin biology (*ANK3, ARID1B, ASH1L, ASXL3, BCL11A, CHD2, CHD8, CREBBP, DPYSL2, FOXP2, KMT5B (SUV420H1), KDM5B, KDM6B, KMT2C, MBD5, MED13L, NRXN1, PHF12, PHF21A, POGZ, PPP2R5D, SCN2A, SETD5, SHANK3, SIN3A, SKI, SLC6A1, SMARCC2, WAC)* in induced NPCs, glutamatergic neurons, and GABAergic neurons *in vitro*. We describe convergent networks that are unique between cell types, and in neurons, enriched not just for synaptic biology, but also epigenetic regulation and, unexpectedly, mitochondrial function. Novel applications of machine learning allowed us to extend our analyses *in silico* across all known NDD genes, resolving how the degree of convergence between risk genes was influenced by clinical associations, biological function, and co-expression patterns in the post-mortem brain. Convergent analyses resolved the genes and cell types that underlie *in vivo* behavioral stratification and successfully predicted drugs capable of suppressing phenotypes in mutant zebrafish, suggesting that precision medicine-based approaches can successfully target shared downstream gene targets between multiple NDD genes. Novel points of convergence in post-mitotic neurons represent exciting new therapeutic targets occurring within a clinically actionable therapeutic window.

## RESULTS

### A systematic comparison of NDD gene effects across neuronal cell types

From 102 highly penetrant loss-of-function (LoF) gene mutations associated with NDD (previously described as 58 gene expression regulation, 24 neuronal communication, and 20 other)^4^, we used gene ontology and primary literature to identify 21 epigenetic modifiers specifically involved in chromatin organization, rearrangement, and modification (*ASH1L, ARID1B, ASXL3, BCL11A, CHD2, CHD8, CREBBP, PPP2R5D, KDM5B, KDM6B, KMT2C, KMT5B (SUV420H1), MBD5, MED13L, PHF12, PHF21A, SETD5, SIN3A, SKI, SMARCC2, WAC*), as well as two transcription factors with putative roles as chromatin regulators (*FOXP2, POGZ*). Three extensively studied synaptic genes (*NRXN1, SCN2A, SHANK3*) and three under-explored neuronal communication genes (*ANK3*, *DPYSL2*, *SLC6A1*) strongly associated with NDD were added (**SI Fig. 1A**). Many of these 29 genes differed in relative frequency of LoF gene mutations between ASD (n=16) and DD (n=4)^61^, schizophrenia^62^, and epilepsy^63,64^ (**Fig. 1A-B, SI Fig. 1B**), as well as general associations with GWAS for many neuropsychiatric disorders (MAGMA^65^) (**Fig. 1C; SI Fig. 1C**), indicating a pleotropic effect consistent with the shared genetic liability across neuropsychiatric disorders^66^. iNPCs, iGLUTs, and iGABAs (**SI Fig. 2A**), as well as their *in vivo* fetal counterparts (**SI Fig. 2B**), expressed all genes prioritized herein^67^.

**Figure 1.**
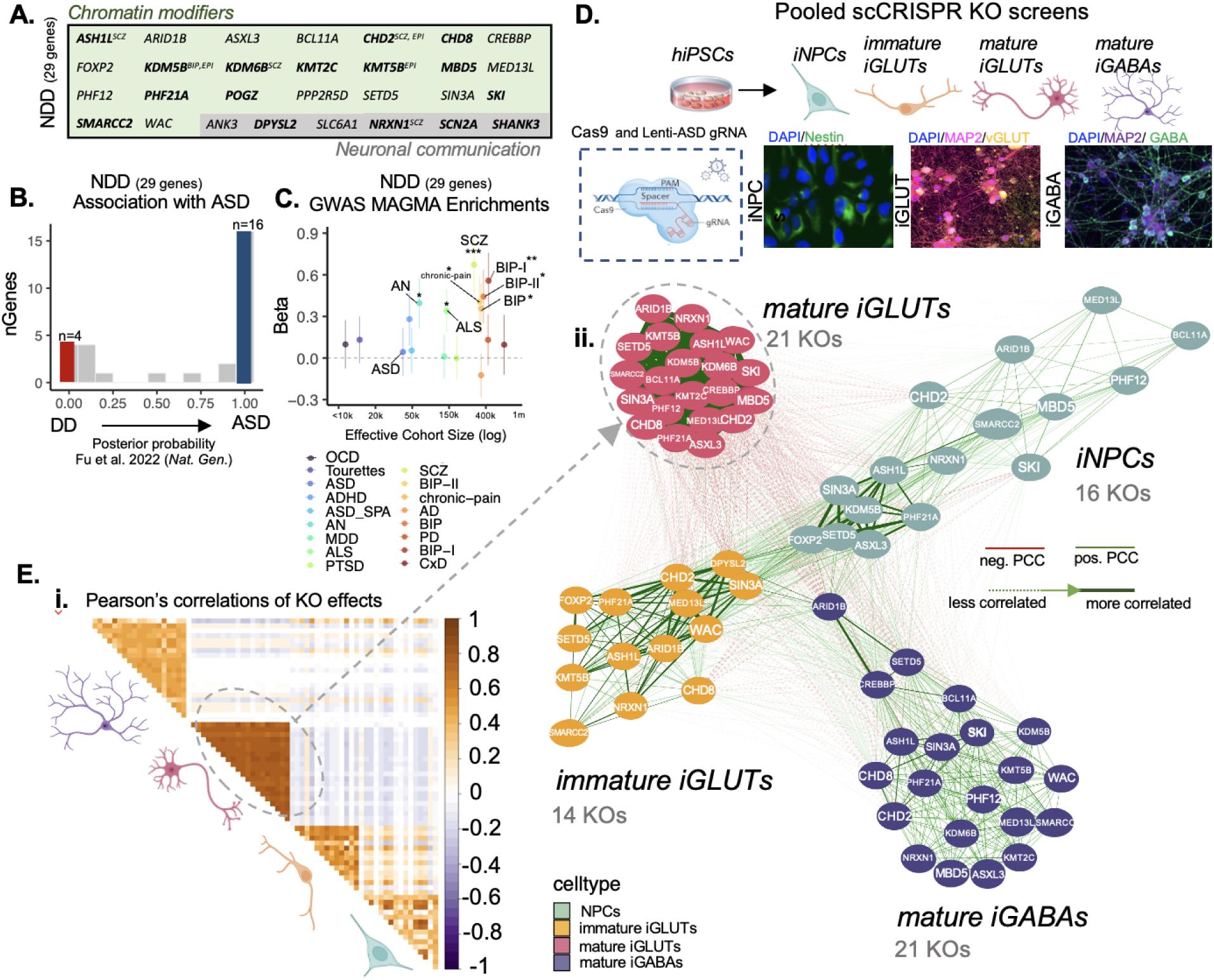
Knock-out (KO) effects of 21 NDD risk genes are most strongly correlated in mature neurons. **(A)** List of rare-variant target risk genes associated neurodevelopmental disorders (NDD) separated by chromatin modifiers and neuronal communication genes. Bold gene names indicate strong associations with ASD based on Fu et al. 2022. Gene targets of rare variants associated with schizophrenia (SCZ), epilepsy (EPI) and bipolar disorder (BIP) are annotated. **(B)** Strength of association with ASD, as estimated by distribution of posterior probability (p.p.) scores from Fu et al. 2022; 4 out of 29 NDD genes were more strongly associated with developmental delay (DD) (blue; p.p.<=0.1) while 16 out of 29 were more strongly associated with ASD (red; p.p.>=0.9). Further annotation of individual risk genes are shown in **SI Figures 1-2**. (**C)** MAGMA enrichments of targeted genes across GWAS for anorexia nervosa (AN), chronic pain, amyotrophic lateral sclerosis (ALS), SCZ, and BIP, BIP-I (bipolar subtype 1), and BIP-II (bipolar subtype 2). **^#^**nominal p-value<0.05, **\***FDR<0.05, **\*\***FDR<0.01, **\***FDR<0.001 **(D)** Schematic of hiPSC-derived cell-type specific scCRISPR-KO screen. Representative immunofluorescence for markers of NPCs (DAPI/Nestin), mature iGLUTs (DAPI/MAP2/vGLUT), and mature iGABAs (DAPI/MAP2/GABA). **(E)** Transcriptomic impact of NDD gene KO represented as the number of nominally significant (p<0.01) differentially expressed genes (DEGs). **(i)** Pearson’s correlation matrix of log2FC DEGs across all NDDs and cell-types. **(ii)** Cross cell-type correlation network diagram across NDD perturbations (number of NDD gene knockout (KO) perturbations resolved indicated in parentheses); the mature iGLUT cluster was most dense, and the iNPC most sparse.

Towards resolving whether regulatory genes confer continuous or distinct periods of susceptibility across neurodevelopment, we knocked out (KO) regulatory NDD genes in neural progenitor cells (SNaPs^68^, here termed iNPCs), immature and mature glutamatergic neurons (iGLUTs)^69^, and mature GABAergic neurons (iGABAs)^70^ (**Fig. 1D**). A pooled CRISPR approach (ECCITE-seq^71^) combined direct detection of sgRNAs and single-cell RNA sequencing to compare loss-of-function effects across 29 NDD genes. The CRISPR-KO library was generated from pre-validated gRNAs (three to four gRNAs per gene; **SI Table 1**). Sequencing of the gRNA library confirmed the presence of gRNAs targeting 24 genes (*ANK3, ARID1B, ASH1L, ASXL3, BCL11A, CHD2, CHD8, DPYSL2, FOXP2, KMT5B (SUV420H1), KDM5B, KDM6B, KMT2C, MBD5, MED13L, NRXN1, PHF12, PHF21A, SCN2A, SETD5, SIN3A, SKI, SMARCC2, WAC*), but three *(DPYSL2, FOXP2, SCN2A*) were present at lower frequency (**SI Fig. 3B-C**).

Control hiPSCs were induced to iNPCs, iGLUTs, and iGABAs (**SI Fig. 3A**), transduced first with lentiviral-Cas9v2 (Addgene #98291) and subsequently with the pooled lentiviral gRNA library three days before harvest, at day 7 (iNPC and immature iGLUT), day 21 (iGLUT), and day 36 (iGABA) (experimental workflow **SI Fig. 4A;** computational workflow **SI Fig. 4B;** experimental validation of CRISPR editing efficiency in **SI Fig. 5**). After filtering and QC (**SI Fig. 4C-E**), we resolved NDD transcriptomes for 118,436 single cells: 25,402 iNPC, 38,097 immature (d7) iGLUT, 28,388 mature (d21) iGLUT, and 26,549 mature (d36) iGABA. Because original gene-expression based clustering was driven by cellular heterogeneity, cell quality, and sequencing lane effects (**SI Fig. 6A**), independent of gRNA identity, we removed cells with high expression of subtype markers and adjusted for cellular heterogeneity (**SI Fig. 6B,C; SI Tables 2-3**). ‘Weighted-nearest neighbor’ (WNN) analysis assigned clusters based on both gRNA identity class and gene expression to ensure that cells assigned to a gRNA identity class demonstrated successful perturbation of the targeted NDD gene^72^. For those WNN clusters where most cells were assigned to a single KO target, the transcriptomic signatures were compared to non-targeting scramble control clusters. Altogether, 35,777 cells were used for downstream analyses: 12,107 iNPC, 3,171 immature iGLUT, 11,802 mature iGLUT, and 8,697 mature iGABA). An average of 474 cells were assigned to each individual sgRNA (757 iNPC, 227 immature iGLUT, 562 mature iGLUT, 414 mature iGABA), totaling 33,150 perturbed cells and 2,627 controls (882 iNPC, 90 immature iGLUT, 1,258 mature iGLUT, and 397 mature iGABA). The gene expression patterns of non-perturbed iNPCs and iNeurons (>30% of all pooled cells) were significantly correlated with fetal brain cells and cortical adult neurons.

Successful perturbations (scCRISPR-KO) were identified for 23 NDD genes (**SI Fig. 6,7**): 16 in iNPCs, 14 in immature iGLUT neurons, and 21 in mature iGLUT and iGABA neurons (**SI Fig. 6**). Nine NDD genes were perturbed in all four cell types (*ARID1B, ASH1L, CHD2, MED13L, NRXN1, PHF21A, SETD5, SIN3A, SMARCC2*; **SI Fig. 7A,B**). For most NDD genes, KO in mature iGLUTs yielded the largest number of differentially expressed genes (DEGs, pFDR<0.05) (**SI Fig. 7B**), an effect that was not driven by differences in the extent of perturbation of the NDD gene itself between cell types (**SI Fig. 7Ci**). The transcriptomic effects of individual NDD genes cluster by cell type: the strongest NDD gene correlations are in mature iGLUTs (i.e., all nominally significant (p<0.01) log2FC DEGs are most highly correlated with each other and least correlated with the other cell types, whether relative to all scramble control cells (**Fig. 1Ei,ii; SI Fig. 7Cii**) or random subsets of scramble control cells (**SI Fig. 8A,B**). DEGs across individual NDDs shared significant gene ontology enrichments (**SI Fig. 8C**), with mature iGLUTs frequently enriched for SCZ GWAS genes (12 of 21 NDD genes), whereas mature iGABAs for migraine GWAS genes (8 of 21) (**SI Fig. 9**).

Unsurprisingly, given the greater within cell-type correlations between NDD genes and the unique pathway enrichments across cell-types, very few DEGs shared significance and direction of effect for the same NDD gene perturbation across all four cell-types (FDR adjusted p_meta_<0.05, Cochran’s heterogeneity Q-test p_Het_ > 0.05; computational workflow, **SI Fig. 10A**); in fact, the only common DEG between cell types was frequently the targeted NDD gene itself. With a more relaxed statistical threshold (nominal p-value <0.05), modest shared effects of individual NDD genes could be resolved across cell types. These effects rarely resulted in perturbation of the other NDD genes themselves (**SI Fig. 10B**), showed very little overlap between NDD genes (**SI Fig. 10C**), and no significant enrichments with psychiatric GWAS after multiple testing correction (**SI Fig. 10D**).

### NDD gene knockouts resulted in cell-type-specific convergent genes and networks that were strongest in glutamatergic neurons

“Convergent genes” (**Fig. 2**) are those DEGs with significant and shared direction of effect across all NDD gene perturbations (FDR adjusted p_meta_<0.05, Cochran’s heterogeneity Q-test p_Het_ > 0.05)^73,74^ (computational workflow, **Fig. 2A**). Across the nine NDD genes perturbed in all four cell types (*ARID1B, ASH1L, CHD2, MED13L, NRXN1, PHF21A, SETD5, SIN3A, SMARCC2)*, convergence was highly cell-type specific (**Fig. 2; SI. Fig. 11A-C; SI Data 2**). Although the strength of convergence correlated across cell types (**Fig. 2C,ii**), it was greatest in mature iGLUTs (quantified as the ratio of convergent genes to the average number of DEGs across all 152 unique two-to-five gene combinations of these nine NDD genes) (**Fig. 2C,i**).The unique “top” convergent genes (**Table 1**) showed little overlap across all cell-types, with mature iGLUTs (11,473) having the largest absolute number of convergent genes (**Fig. 2D**). Convergent genes were enriched for schizophrenia GWAS loci (MAGMA^65^, FDR <0.05) (**Fig. 2Ei**), rare ASD and FMRP target genes (FDR <0.05) (**Fig. 2E,ii**), and pathways involved in neurodevelopment, mitochondrial function, and translational regulation **(SI Fig. 12).** When tested again across the 21 NDD genes perturbed in both iGLUTs and iGABAs, mature iGLUTs again showed the largest absolute number of convergent genes (iGLUTs, 10,557, **SI Fig. 13A**; iGABAs, 892, **SI Fig. 13B**). Intriguingly, although convergent genes were highly cell-type-specific, those NDD gene combinations that were highly convergent in one cell type were likely to be convergent in others; in neurons, top convergent sets most frequently included *ARID1B*, *SETD5* and *NRXN1* **(SI Fig. 11D).**

**Figure 2.**
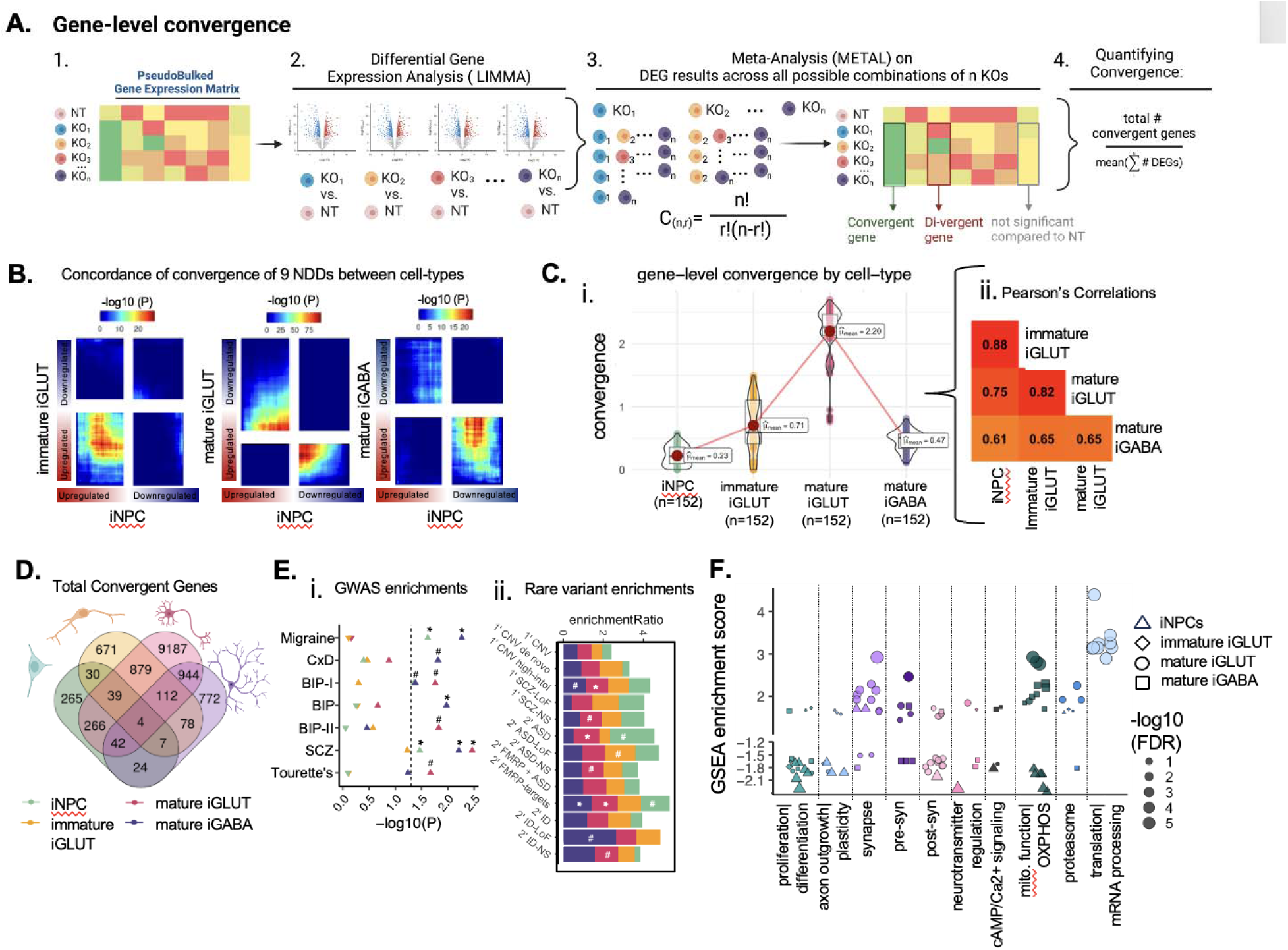
Gene-level convergence is greatest in mature glutamatergic neurons. In total, nine NDD genes showed evidence of knockout across all four cell types: *ARID1B, ASH1L, CHD2, MED13L, NRXN1, PHF21A, SETD5, SIN3A, SMARCC2*. For these nine, “convergent genes” are defined as those differentially expressed genes (DEGs) with significant and shared direction of effect across all NDD gene perturbations. **(A)** Schematic explaining cell-type specific convergence at the individual gene level via differential gene expression meta-analysis (FDR adjusted pmeta<0.05, Cochran’s heterogeneity Q-test pHet > 0.05). **(B)** Convergence across 9 NDD genes is unique to each cell type, using rank-rank hypergeometric (RRHO) test to explore correlation of convergent genes shared across 9 NDD perturbations (RRHO score = -log10*direction of effect) between cell-types. The top right quadrant represents down-regulated genes (meta-analysis z-score >0) for the y-axis and x-axis cell-type. The bottom left quadrant represents up-regulated convergent genes (meta-analysis z-score <0) for the y-axis and x-axis cell-type. Significance is represented by color, with red regions representing significantly convergent gene expression. **(C) (i)** The average strength of convergence, measured as the ratio of convergent genes to the average number of DEGs across all 152 unique combinations of 2-5 genes from the nine NDD genes, was highest in iGLUTs. **(ii)** The magnitude of convergence between the same NDDs tested in different cell types was highly correlated (Pearson’s correlation, P_holm_<2.2e-16); with the strongest relationship between immature and mature iGLUTs. **(D)** Venn diagram representing the absolute overlap (regardless of direction of dysregulation) of cell-type specific convergent genes shared across 9 NDDs. **(E) (i)** MAGMA enrichment –log10(p-value) of cell-type-specific (color of points) convergence and GWAS-risk associated genes with significance after multiple testing correction indicated as follows: ^#^unadjusted p-value=<0.05, *FDR<=0.05, **FDR<0.01, ***FDR<0.001. The direction of the triangles indicates a positive (upwards triangle) or negative (downwards triangle) enrichment beta. **(ii)** Over-representation analysis (ORA) enrichment ratios of cell-type-specific (color of bars) convergence and rare variant target genes. Significance after multiple testing correction indicated as follows: ^#^unadjusted p-value=<0.05, *FDR<=0.05, **FDR<0.01, ***FDR<0.001. **(F)** Gene set enrichment analysis (GSEA) identified downstream pathways involved in neural proliferation, neurite outgrowth, synaptic vesicle transport, and mitochondrial function as cell-type specific targets of convergent genes across 9 NDDs. Results were filtered for pathways with nominal p-values <0.05. Normalized GSEA enrichment scores represent the direction of enrichment based on the meta-analyzed Z-score for each convergent gene. Cell-type is represented by shape and the size of each point represents the –log10(FDR).

**Table 1.**
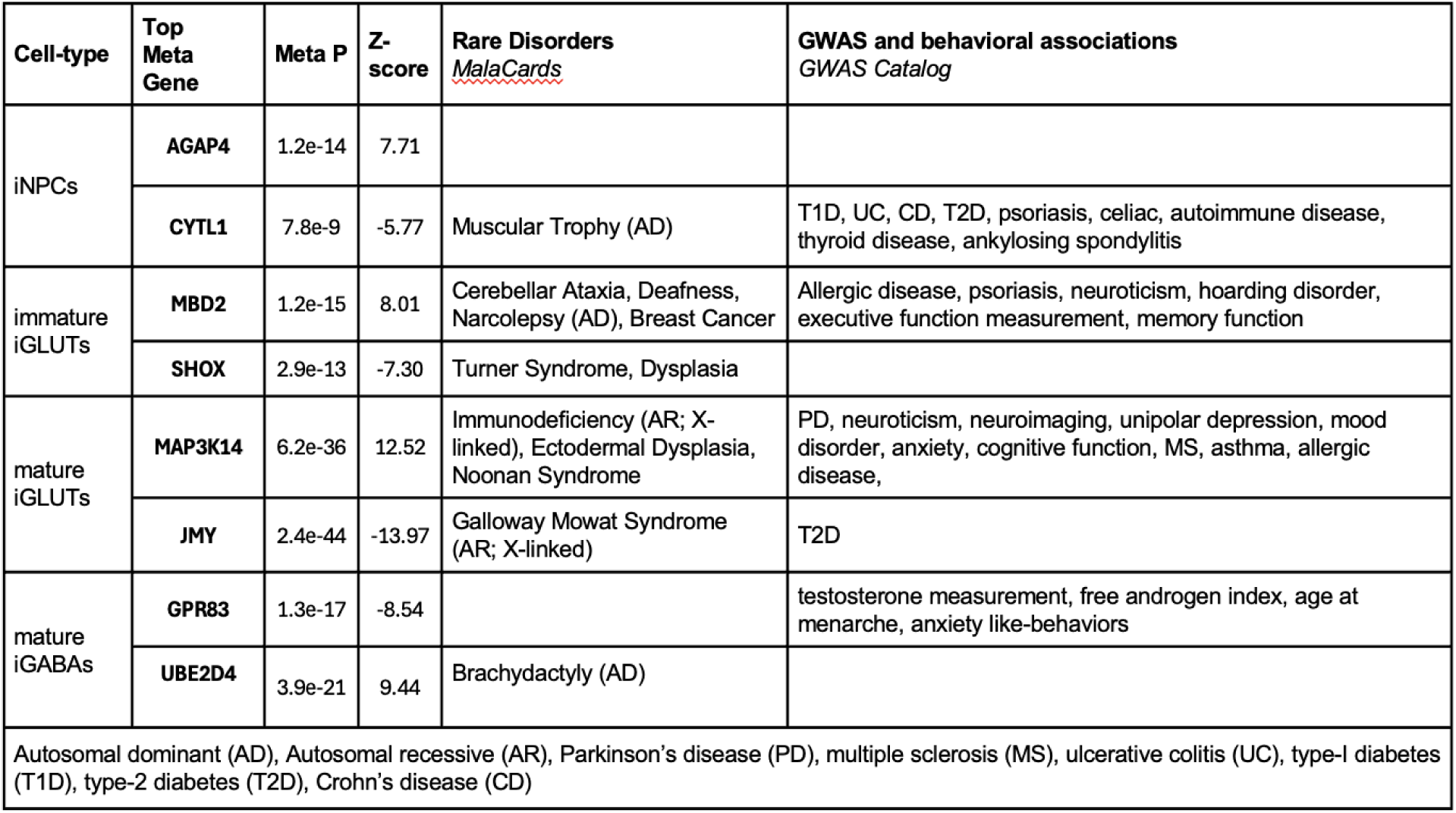
Disorder and behavioral associations of top convergent up and down-regulated genes by cell-type from MalaCards, OMIM, and GWAS catalogue.

Given that the biological impact of convergence is likely to be impacted by the strength of shared gene regulatory relationships and functions, we re-examined convergence within the framework of co-expression networks (Bayesian bi-clustering). “Convergent networks” (**Fig. 3**) are co-expressed genes that share similar expression patterns across NDD gene perturbations^73,74^ (computational workflow, **Fig. 3A**). The network connectivity score (“network convergence”) informs the strength and composition across cell types (i.e., networks with more interconnectedness and containing genes with greater functional similarity have increased convergence). Convergent networks generated from the 9 NDD genes perturbed in all four cell types (**Fig. 3B**) or across the 21 NDDs genes in both iGLUTs and iGABAs (**SI Fig. 13C**) revealed the greatest convergent network strength in iGLUTs. Network-level convergence was weakly correlated between cell types **(Fig. 3C**); the number of convergent unique network nodes was greatest in iGLUTs, distinct across cell types (**Fig. 3D**; **Tables 2-4; SI Data 2**), and significantly enriched for rare variants linked to schizophrenia and ASD (**Fig. 3E**; **Tables 2-4).** Convergent networks in iNPCs highlighted pathways associated with neurogenesis (e.g., cell cycle, cell division, EPO signaling) (**Fig. 3F**), while in mature iGLUTs they were enriched for synaptic function (transmembrane transport and receptor signaling, secretory vesicles, SNARE complex) (**Fig. 3G**).

**Figure 3.**
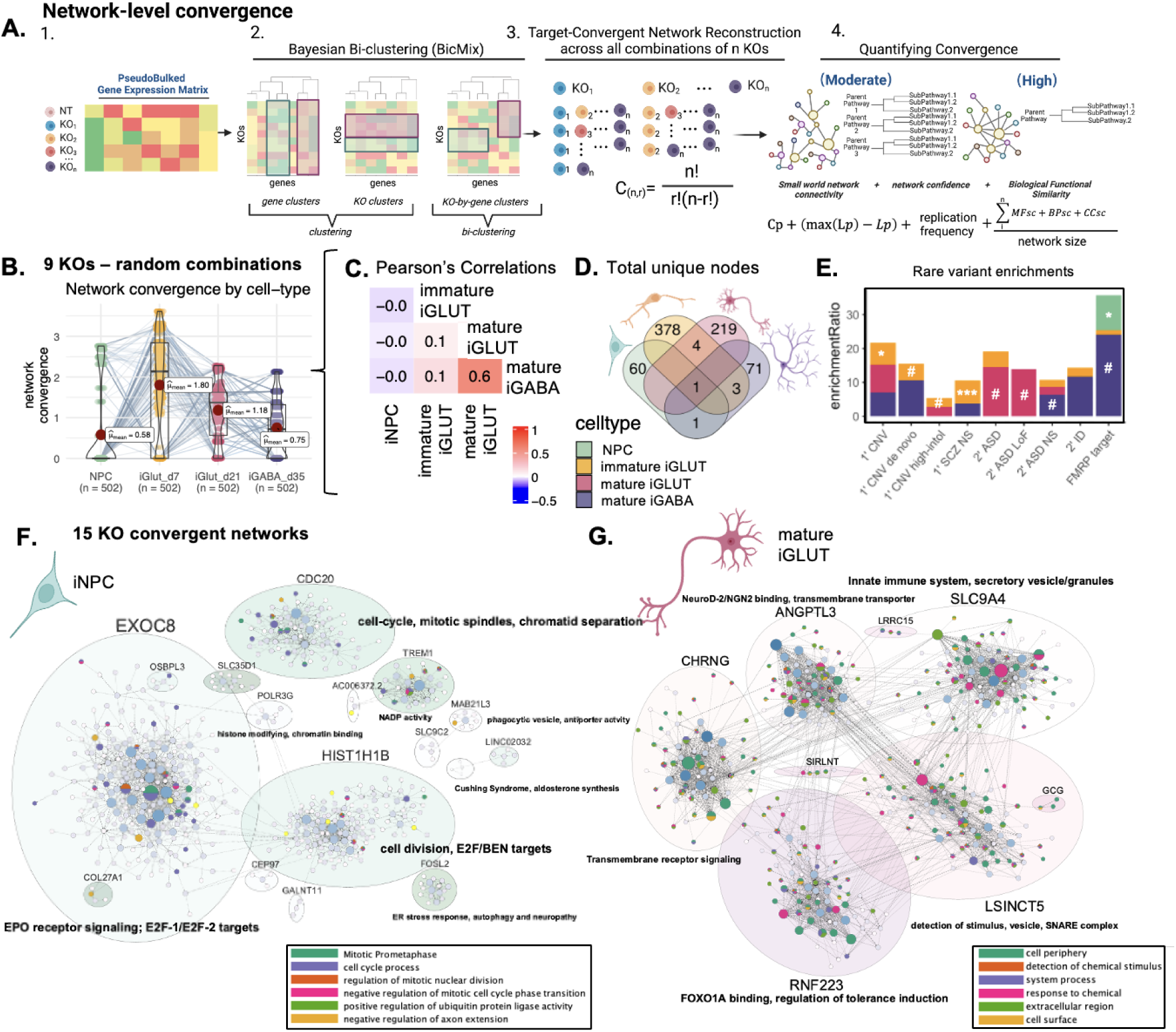
Network-level convergence resolves cell-type-specific and developmental-specific node genes. “Convergent networks” are co-expressed genes that share similar expression patterns across NDD gene perturbations, here resolved for the nine NDD knockouts resolved across all four cell types: *ARID1B, ASH1L, CHD2, MED13L, NRXN1, PHF21A, SETD5, SIN3A, SMARCC2*. **(A)** Schematic explaining cell-type specific convergence at the network level using Bayesian bi-clustering and unsupervised network reconstruction. **(B)** Strength of network convergence across all random combinations of 9 NDD KO perturbations by cell-type. **(i)** The mean strength of network convergence is significantly different by cell-type, with the highest convergence present in immature iGLUTs. The same KO combinations tested in one cell type may not resolve convergence in another cell type. Each point represents a resolved network, and its calculated convergence strength. Dots that represent the same combinations of KO perturbations, but tested in each cell type, are connected by a line. **(C)** Convergent network strength was most correlated between mature iGLUTs and iGABAs (Pearson’s Correlation Coefficient (PCC) = 0.6, P_Holm_ <2.2e-16). Convergent network strength in iNPCs was not correlated with network strength in neurons. **(D)** Venn diagrams of the total number of unique node genes within convergent networks for each cell-type. The lack of overlapping node genes between cell types **(D),** as well as the weak correlations of convergence strength between immature and mature cell-types **(C),** suggest greater cell-type specificity in the magnitude of network-level convergence compared to gene-level convergence. **(E)** Enrichment ratios from over-representation analysis (ORA) of cell-type specific (color of bars) convergent node genes for rare variant targets. (^#^unadjusted p-value=<0.05, *FDR<=0.05, **FDR<0.01, ***FDR<0.001). **(F, G)** Representative cell-type specific network plots for convergence across 15 genes (*ARID1B, ASH1L, ASXL3, BCL11A, KDM5B, CHD2, MBD5, MED13L, NRXN1, PHF12, PHF21A, SETD5, SIN3A, SKI, SMARRC2)* from **(F)** iNPCs and **(G)** mature iGLUTs. Network genes were filtered for protein-coding genes, clustered, and annotated based on the primary node gene for each cluster. Gene set enrichment analysis of the networks identified unique functions by cell type. Convergent networks in iNPCs were enriched for pathways associated with neurogenesis (e.g., cell cycle, cell division, EPO signaling), while in mature iGLUTs for pathways associated with synaptic function (transmembrane transport and receptor signaling, secretory vesicles, SNARE complex).

**Table 2.** Disorder and behavioral associations of top nodes by cell-type from MalaCards, OMIM, and GWAS catalogue.

| Cell-type | Top protein-coding nodes across all networks | Rare disorders *MalaCards* | GWAS and behavioral associations *GWAS Catalog* |
| --- | --- | --- | --- |
| iNPCs | KIAA2012 |  | ADHD/conduct disorder (rs1521882), educational attainment (rs12623702, rs4675248, rs2160317, rs2177083, rs34189321,rs58100125), migraine/T2D (rs6748072), amygdala volume (rs72936662) |
| immature iGLUTs | MYH15 | Deafness (AR); de novo SCZ CNV | Social interaction measurement (rs13082569), cognitive function (rs3860537), unipolar depression (rs1531188), MDD (rs113689582), insomnia (rs62266174, rs6768511, rs6786515, rs6795280), BIP (rs1531188), ANX (rs4855559), educational attainment (rs115910830, rs2290601, rs3860537, rs60785803) |
| mature iGLUTs | GALNTL5 | ASD, Spastic paraplegia 48 (AR) |  |
| mature iGABAs | EREG | Epilepsy, Immune Deficiency Disease | ADHD (rs1350666), wellbeing measurement (rs112444088), learning & memory (pathway) |
| Autosomal recessive (AR), Anxiety disorder (ANX), Bipolar Disorder (BIP), Type-2 diabetes (T2D) |  |  |  |

**Table 3.**
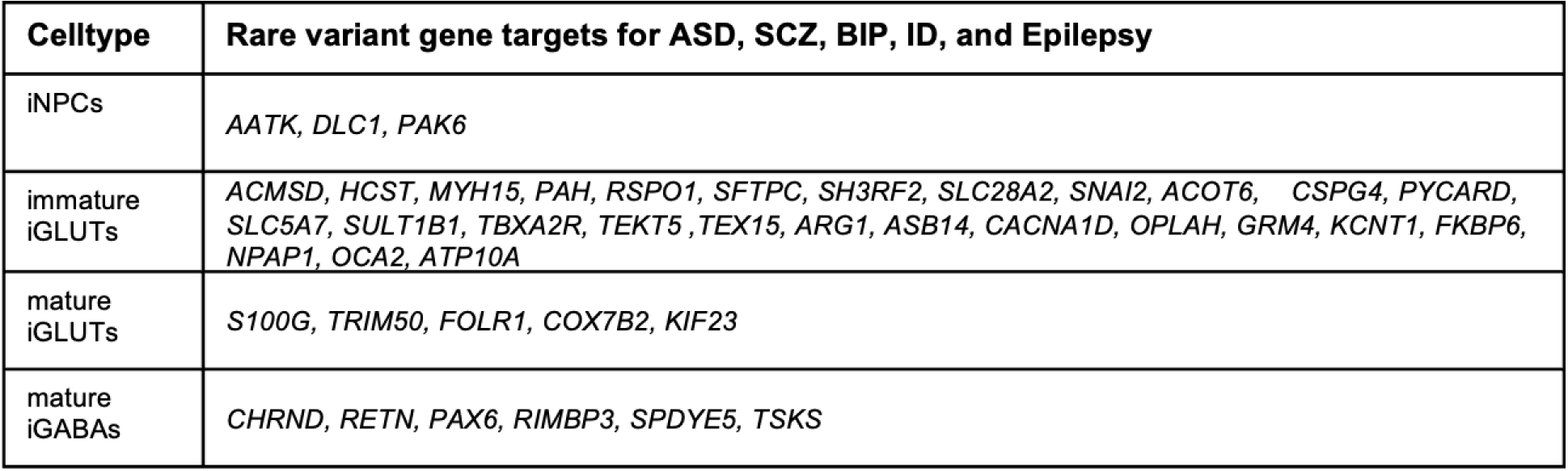
Convergent nodes that overlap with CNV and rare variant target genes for each cell-type.

**Table 4.**
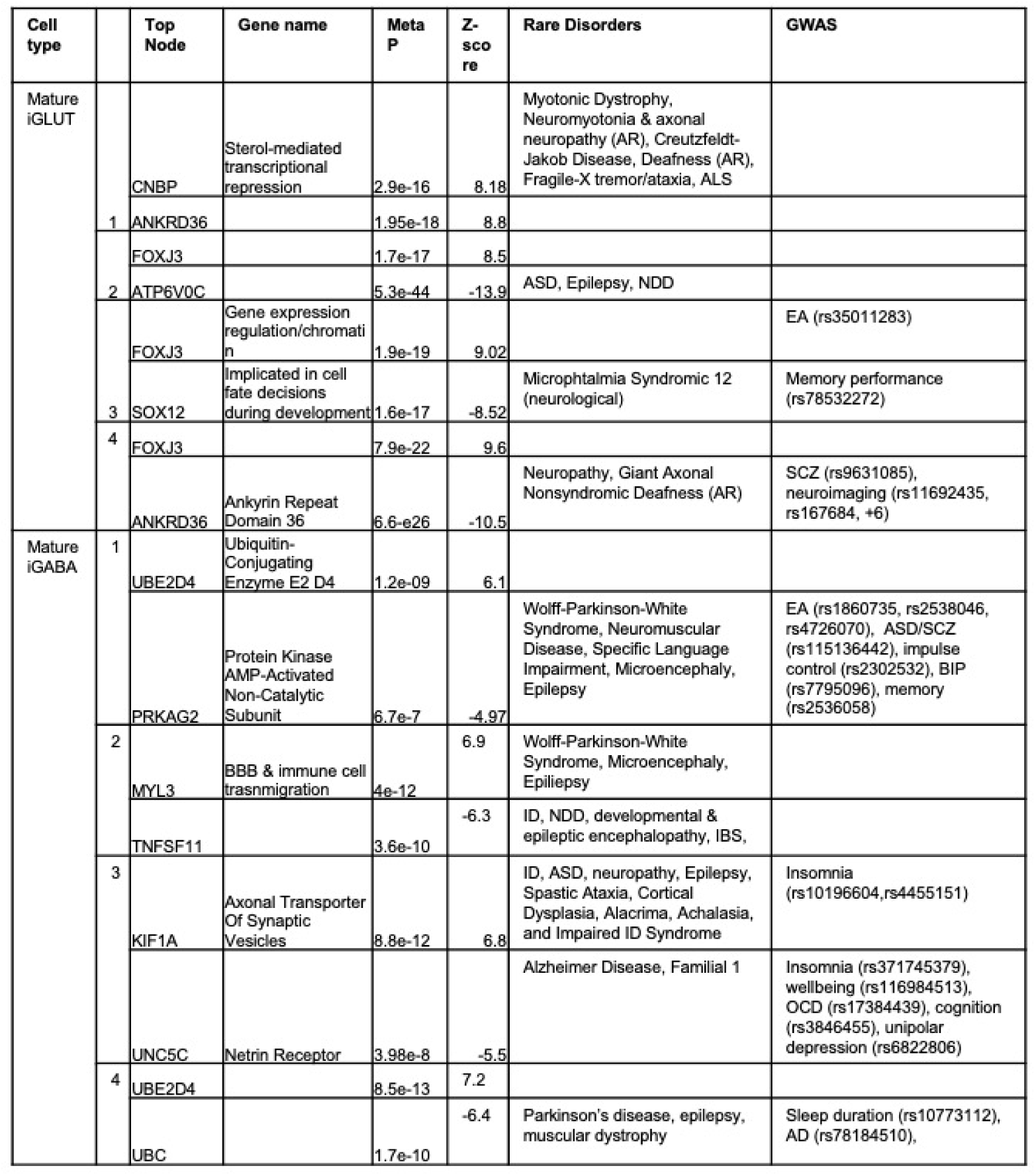
Disorder and behavioral associations of top nodes by cell-type and behavioral set from MalaCards, OMIM, and GWAS catalogue.

### Convergent networks are strongest between NDD genes with shared co-expression patterns in the post-mortem brain, biological annotations (synaptic or epigenetic), and clinical outcomes (ASD or DD)

To resolve the extent to which functional similarity and co-expression patterns between NDD genes predicted convergence, we trained a prediction model (random forest linear regression)^75^ using 70% of our data, evaluated it using 30% of our data, and validated in an external dataset^73^ (computational workflow, **Fig. 4A**; model predictor variables, **Fig. 6B**; more information **SI Fig. 14,15**). Cell type, brain co-expression (dorsolateral prefrontal cortex, DLPFC), and functional similarity (i.e., gene ontology) correlate with convergence (**Fig. 4C**) and well-predicted gene level convergence (97% variance explained; mean of squared residuals (RMSE)=0.021) (**Fig. 4Di**) and moderately predicted network-level convergence (53% variance explained; RMSE=0.73) (**Fig. 4Dii**). Our trained model accurately predicted gene-level (Pearson’s R=0.998, P<0.001, RMSE=0.15) (**Fig. 4Ei; SI Fig. 15C**) and network-level convergence in our testing set (R=0.72, P<2.2e-16, RMSE=0.85) (**Fig. 4Eii; SI Fig. 15D),** and performed moderately well in predicting network-level convergence (R=0.26, P<0.001, RMSE=0.68) (**Fig. 4Fii; SI Fig. 15Eii)** and to a lesser extent gene-level convergence (R=0.14, P<0.001 RMSE=1.75) (**Fig. 4Fi; SI Fig. 15Ei)** in the external dataset.

**Figure 4.**
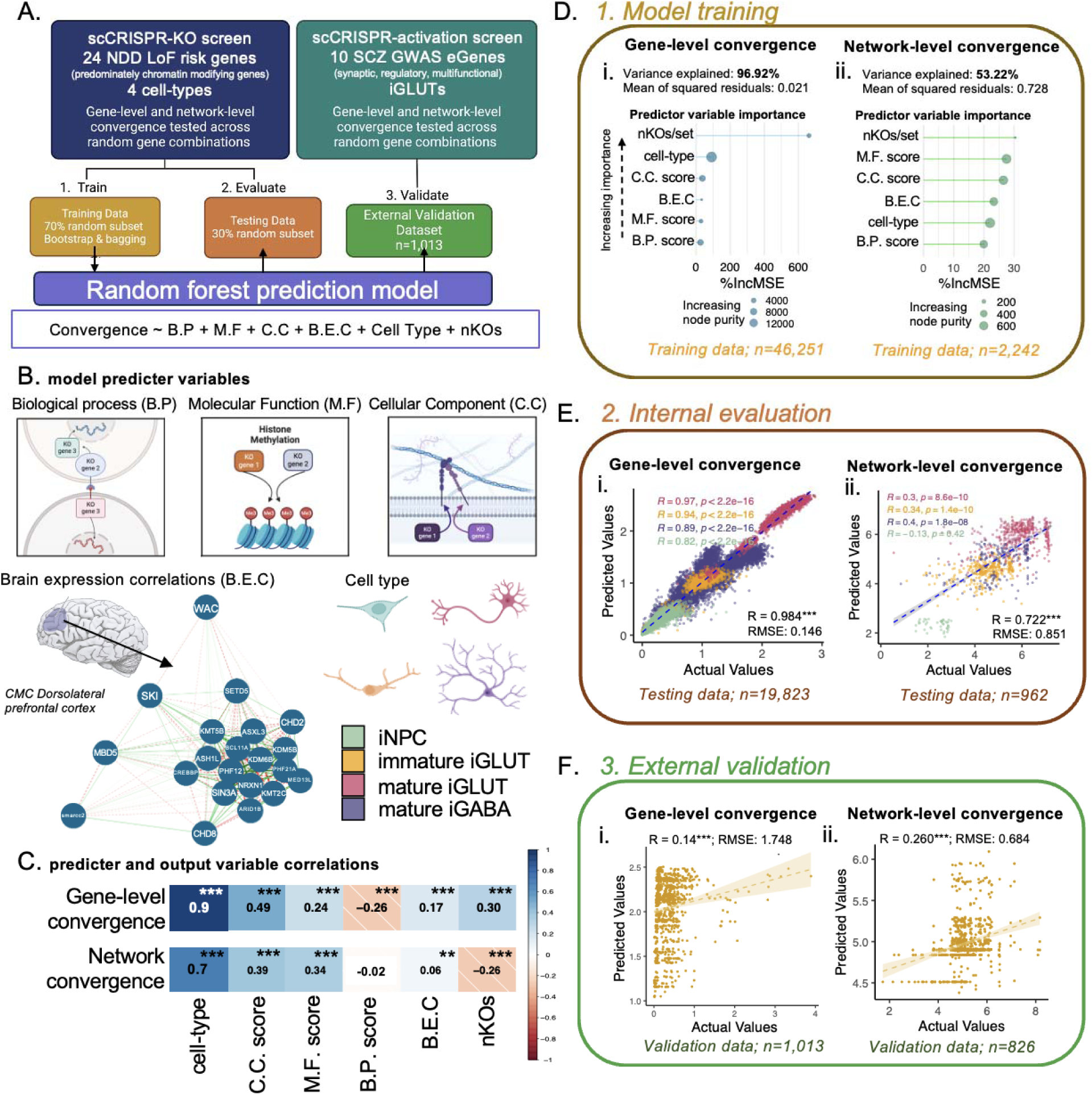
Functional similarity and brain co-expression between NDD genes predict gene-level and network-level convergence, with unique influences by cell-type. **(A)** Schematic for training random forest models for gene and network-level convergence with external validation in a SCZ CRISPRa screen. **(B)** Predictor variables included in the model include scores of functional similarity, dorsolateral prefrontal cortex (DLPFC) brain co-expression, cell-type, and the number of KOs. (**B.P score** = semantic similarity of GO: Biological Process membership between KO genes; **C.C. score** = semantic similarity of GO: Cellular Component membership between KO genes; **M.F score** = semantic similarity of GO: Molecular functions membership between KO genes; **B.E.C** = dorsolateral prefrontal cortex expression correlations between KO genes; **nKOs** = number of KO genes tested for convergence). **(C)** Pearson’s correlations of predictor variables and gene-level and network-level convergence (PBonferroni<=0.01**, PBonferroni<=0.01***). **(D)** Functional similarity, brain co-expression, cell-type, and the number of KOs assayed strongly predicted gene-level convergence (97% variance explained by the model; mean of squared residuals=0.02) and moderately predicted network-level convergence (53% variance explained; mean of squared residuals=0.73). **(i-ii)** Importance of each of the predictor variables was assessed by two metrics: the percent mean increase in squared residuals (%IncMSE) and the increase in node purity. In the model – number of KO genes in a set is the most important predictor of convergence based on %Inc MSE, but not node purity. However, the impact of nKOs on gene-level convergence is much stronger – likely an artifact of the method used for measuring convergence. For network level convergence, each variable has a IncMSE between 20-30%. **(E)** Internal evaluation of the model using 30% of the original data resulted in high concordance between convergence predicted by the model and the measured convergence. Predicted gene-level **(i)** [gene-level convergence: n=19,823; Pearson’s R=0.984; p<2.2e-16; root mean squared error (RMSE) =0.15] and network-level **(ii)** convergence [network-level convergence: n=962; rho=0.722; p<2.2e-16; RMSE=0.85)] by the model strongly correlated with the measured convergence in the testing sets. Correlation of predicted vs. accrual convergence values are color-coded by cell-type with corresponding color-coded correlations and p-values listed in the upper right corners of the scatterplots. **(F)** External validation in an independent scCRISPRa screen of SCZ target genes predicted showed moderate, but significant, correlation between convergence predicted by the model and the measured convergence. **(i)** gene-level (n=1013, R=0.14, p=1.1e-05, RMSE=1.748) and **(ii)** network-level convergence (n=826, R=0.26, p=2.9e-14, RMSE=0.68).

To query whether convergence reflected clinical associations to ASD or DD, we again quantified convergence as the ratio of convergent genes to the average number of DEGs (see **Fig. 2E**), here across all (2-5 gene) combinations of all NDD genes perturbed in each cell type (e.g., 27,824 unique combinations of 21 NDD genes in iGLUTs and iGABAs; **SI Fig. 16A**). Convergence, both gene-level (**SI Fig. 16A,C**) and network-level (**SI Fig. 16B,D**), was greater between genes with stronger associations to ASD compared to DD^61^, particularly in mature neurons (**SI Fig. 16E-F**). Yet this analysis was limited by the relatively small number of predominantly ASD (n=16) and DD (n=4) included in our dataset (**Fig. 1B**).

To extend our comparisons of convergence across larger sets of NDD genes, particularly those clinically defined as predominantly ASD or DD genes^61^, or those with biologically annotated synaptic or epigenetic roles^4^, we asked if it was possible to train a machine learning model to predict cell-type-specific impacts of CRISPR knockout of all 102 NDD genes^4^. An integrative Linear Network of Cell Type Phenotypes (**LNCTP**) model, previously trained on >2.8 million nuclei from the prefrontal cortex across 388 individuals, accurately imputes single-cell expression following simulated perturbations^76^. By retraining the LNCTP model using our scCRISPR-KO data (**Fig. 5A**), we resolved convergent genes within three *in silico* post-mortem brain network models (bulk prefrontal cortex (PFC) tissue, excitatory neurons only, and inhibitory neurons only), noting that the LNCTP model better replicates experimental iGLUT data (**Fig. 5B**).

**Figure 5.**
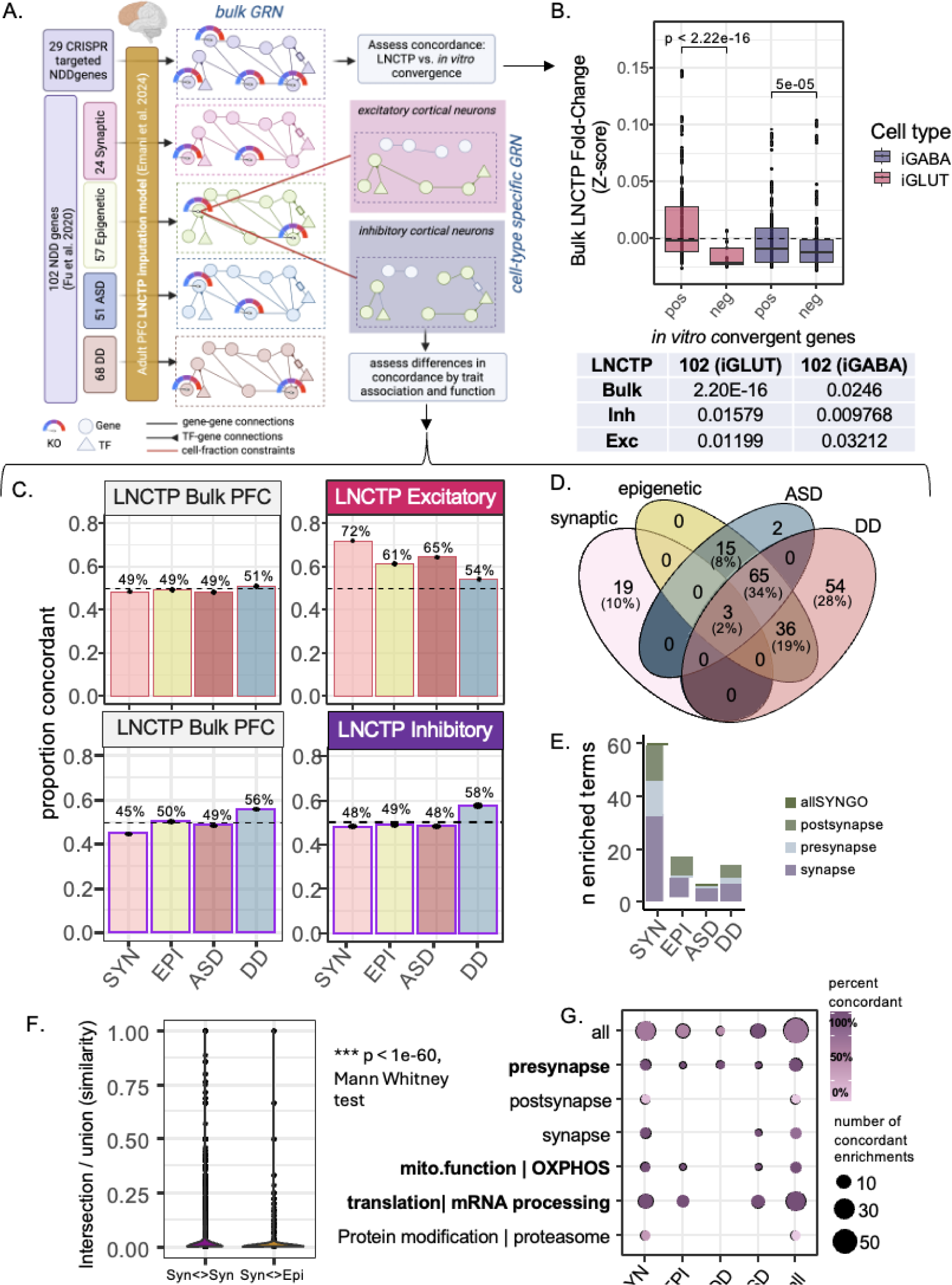
LNCTP predicts effects of convergent genes *in silico*. (**A)** LNCTP imputation and perturbation model: an energy-based network model is trained to impute bulk and cell-type specific expression data in the prefrontal cortex over a population of post-mortem individuals from PsychENCODE using a panel of 1325 genes and embedded cell-type specific Gene Regulatory Networks (GRNs) (*LNCTP in silico model*); a chosen gene is then perturbed by fixing its expression, and the effects on other genes are predicted by the model; *in silico* category-specific convergent genes are then identified by analyzing the fold-changes across subjects (*LNCTP Simulating Perturbations*). **(B)** Predicted *in silico* log fold-changes for the *in vitro* positive and negative convergent genes across the 29 CRISPR perturbations, in Bulk, Excitatory and Inhibitory neuron networks (*LNCTP Simulating Perturbations*, 2-tailed t-test p-values shown). **(C)** Proportion of genes showing same direction fold-changes in *in silico* and *in vitro* perturbations across classes of perturbation and cell-type (left), and the intersection of convergent in silico genes across classes of perturbation (*LNCTP in silico convergent genes*, synaptic-epigenetic genes reduced and ASD-DD genes enriched, p<1e-3, 2-tailed hypergeometric test). **(D)** Venn diagram of *in silico* convergent genes across all categories by clinical (ASD vs DD) or functional (synaptic vs epigenetic) annotation. **(E)** Number of terms enriched for convergent genes across all categories for 102 in silico perturbations. **(F)** Semantic distance of pairs of enriched terms within or between sets determined by synaptic and epigenetic convergent gene rankings (*LNCTP semantic distance test*, 2-tailed Mann Whitney test) (G) Percent of concordant genes in each perturbation and ontology category within the leading-edge enriched genes (*LNCTP in silico convergent genes*).

Expanded LNCTP *in silico* comparisons across all 102 NDD genes (**Fig. 2; SI. Fig. 17**) predicted greater convergence in excitatory neurons compared to inhibitory neurons, consistent with our *in vitro* findings (**Fig. 2C, 3D**), even more so for synaptic NDD genes (n=24) relative to regulatory genes (n=58) (**Fig. 5C**). Predominantly ASD genes (n=50) had greater predicted convergence in excitatory neurons (**Fig. 5C**), whereas predominantly DD genes (n=40) in inhibitory neurons (**Fig. 5C**). Overall, across functional or clinical categories, despite limited overlap in specific convergent genes (**Fig. 5D**) and terms (**Fig. 5E, F**), there was overall enrichment for synaptic, epigenetic, and mitochondrial biology (**Fig. 5G**), consistent with *in vitro* scCRISPR-KO (**Fig. 2F**).

### Convergent genes and networks in glutamatergic neurons targeted synaptic, epigenetic, and mitochondrial biology

Convergent genes and networks revealed cell-type-specific disease (**Fig. 2E**) and functional enrichments (**Fig. 2F, 5G,6A-B**), many consistent with established NDD etiology in neurogenesis^22,27,54–60^ and synaptic biology^47–50^. For example, iNPCs were significantly enriched for pathways involved in proliferation and differentiation, whereas mature iGLUTs showed unique enrichments in neuronal communication (e.g., pre-synaptic function) and regulation of gene expression (e.g., mRNA processing and protein translation). Unexpectedly, both mature iGLUT and iGABA neurons were enriched for mitochondrial biology (e.g., oxidative phosphorylation: mature iGLUTs: NEs=2.8, p<2.2e-16, FDR<0.001; mature iGABAs: NES=1.67, p=0.023, FDR<0.05).

Functional validation of five NDD genes (*KMT5B, NRXN1, CHD8, ASH1L, ARID1B*) in inducible Cas9 (iCas9)^77^ NPCs (CD184⁺/CD133⁻ NPCs) in arrayed format revealed effects on proliferation (Ki67; **Fig. 6C; SI Fig. 18A**), neurogenesis (NPCs: CD184+/CD44-/CD24+, neurons: CD184-/CD44-/CD24+; **SI Fig. 18B**), and gliogenesis (astrocytes: CD184+/CD44+; **SI Fig. 18C**) that varied between genes. Likewise, a pooled CRISPR analysis in iCas9 cortical organoids confirmed effects on neurogenesis, again with variable effects between NDD genes (**SI Fig. 19**).

**Figure 6.**
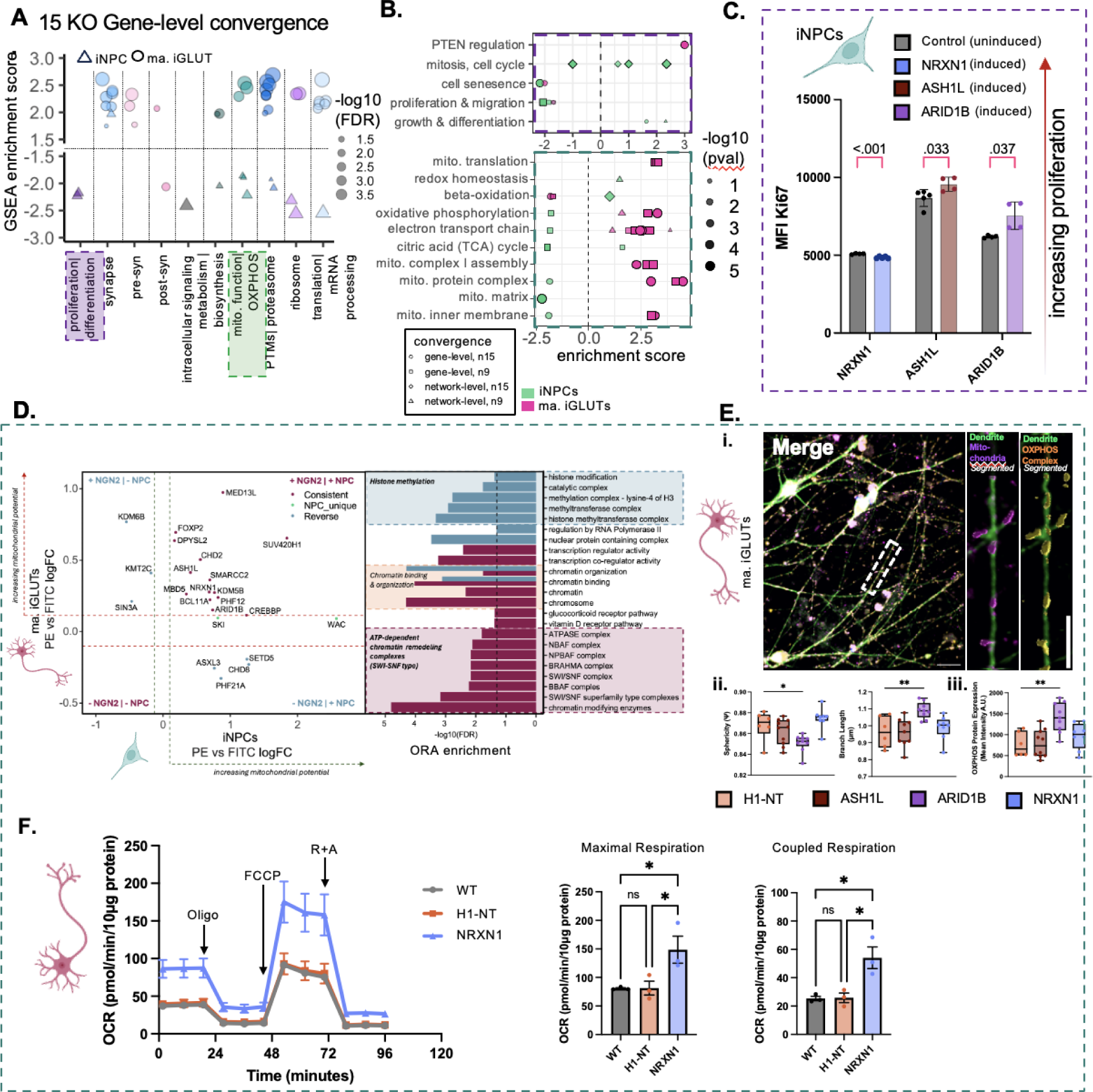
NDD knock-outs converge on mitochondrial function. **(A)** Gene set enrichment analysis (GSEA) identified downstream pathways involved in neurogenesis, neurite outgrowth, synaptic biology, and mitochondrial function as cell-type specific targets of convergent genes across 15 NDD KOs **(*ARID1B, ASH1L, ASXL3, BCL11A, KDM5B, CHD2, MBD5, MED13L, NRXN1, PHF12, PHF21A, SETD5, SIN3A, SKI, SMARRC2)*** in iNPCs and mature iGLUTs. Results were filtered for pathways with nominal p-values <0.05. Normalized GSEA enrichment scores represent the direction of enrichment based on the meta-analyzed Z-score for each convergent gene. Cell-type is represented by shape and the size of each point represents the –log10(FDR). **(B)** Summary of network and gene-level pathway enrichments (from Fig. 2**-3**) for shared effects of nine and fifteen NDD KOs in iNPCs and mature iGLUTs. **(C)** Proliferation assessment of NPCs using Ki-67 median fluorescence intensity (MFI) measured with flow cytometry, wildtype (purple: no iCas9 induction) versus knockout (green: doxycycline to induce iCas9). 4-6 replicates per condition, unpaired t-test with Welch correction; p-values corrected for multiple comparisons using FDR. **(D)** Scatter plot of gRNA log₂ fold-change (high-(PE-high) and low-(FITC-high) Δψm-sensitive dye JC-1 membrane-potential fractions) in NPCs (x-axis) and mature iGLUT neurons (y-axis), with points colored by enrichment category (shared NPC and iGLUT in red; distinct between NPC and iGLUT in blue). Right: Bar chart of –log₁₀(FDR) for over-represented gene sets in the tene gene KOs enriched in both lineages. **(E)** (i) High resolution, high-throughput microscopy of mitochondrial morphology (scale bar 10 μm): an isolated dendrite labelled with a dendritic marker (MAP2), mitochondrial marker (TOMM20) and marker of the OXPHOS complex (Total OXPHOS) (scale bar 5 μm). **(ii)** Effect of ARID1B-KO on mitochondrial sphericity and branch length independent of changes in mitochondrial volume and surface area **(SI Fig. 20-21). (iii)** Effect of ARID1B-KO on average fluorescence intensity of OXPHOS proteins. Each datapoint indicates one well of a 96-well, representing hundreds of μm^2^ of neuronal area and tens of thousands of individual mitochondria (*adjusted p<0.05, ** adjusted p<0.01). **(F)** Effect of NRXN1-KO on maximal respiration and coupled respiration in iGLUTs. Oligo: oligomycin; FCCP: carbonyl cyanide 4-(trifluoromethoxy) phenylhydrazone; R+A: rotenone and antimycin A. Data are presented as mean ± SEM. Statistical analysis was performed using one-way ANOVA. *p<0.05. Each datapoint represents one well of a 24-well Seahorse assay plate. The experiment was independently replicated twice.

To assess how loss of NDD-associated genes affects mitochondrial function, we performed a pooled CRISPR knockout screen using a nearly identical library (same backbone, guide density, and control set) in the H1-iCas9 line. Transduced cells were differentiated into NPCs and iGLUTs by day 21, stained with the Δψm-sensitive dye JC-1, and sorted by fluorescence-activated cell sorting (FACS) into high-(PE-high) and low-(FITC-high) membrane-potential fractions, following amplicon sequencing to quantify gRNA representation in each fraction (**Fig. 6D**). Of the fifteen KOs, ten resulted in elevated mitochondrial membrane potential (MPP) in both NPCs and iGLUTs, the remaining five caused cell-type-specific impacts on mitochondrial membrane potential. Pathway enrichment of the ten NDD genes that increased mitochondrial membrane revealed a convergence on chromatin remodeling complexes, microRNAs, and transcription factors.

For three NDD KOs (*ASH1L, ARID1B, NRXN1)*, we validated mitochondrial effects in arrayed format, using a platform with the ability to resolve dose-dependent changes in mitochondrial fragmentation following pharmacological insults **(SI Fig. 20**). By high content imaging, we analyzed and quantified 1 × 10^4^ mitochondria per genotype, with morphological measurements taken for mitochondrial (TOMM20-positive) volume, surface area, and sphericity (roundness) as well as total OXPHOS complex, within neuronal dendrites (MAP2-positive) of mature (d21) iGLUTs. Among the three NDD KOs, *ARID1B* resulted in increased mitochondrial networking (indicated by decreased mitochondrial sphericity and increased branch length; one-way ANOVA, Šidák’s adjusted p=0.0213 and p=0.0081 respectively) concomitant with increased levels of OXPHOS proteins (one-way ANOVA, Šidák’s, adjusted p=0.0024) (**Fig. 6E; SI Fig. 21A**), overall consistent with increased mitochondrial efficiency. Second, we tested oxygen consumption using Seahorse Cell Mito Stress test. NRXN1 KO resulted in increased coupled and maximal respiration in iGLUTs (one-way ANOVA, p<0.05; **Fig. 6F**); increased mitochondrial reliance, in the absence of fused mitochondria with elevated OXPHOS protein levels point to a possible metabolic overload due to reduced mitochondrial efficiency (**Fig. 6E**). In contrast, ARID1B and ASH1L KOs did not show significant changes in these Seahorse parameters (**SI Fig. 21B–C**). Taken together, both ARID1B and NRXN1 KO neurons show evidence of increased mitochondrial activity, ARID1B KO through enhanced fusion and elevated expression of OXPHOS complexes, whereas NRXN1 KO by increasing OXPHOS activity to meet ATP demands. As observed for neurogenesis in iNPCs, single gene knockouts iGLUTs confirmed convergent effects on mitochondrial biology, finding distinct but related phenotypes between NDD genes.

### Pharmacological targeting of convergent genes reversed behavioral phenotypes in mutant zebrafish

By design, *in vitro* models substantially limit the complexity of the observed impact of NDD genes, lacking higher circuit-level effects. Towards applying molecular convergence *in vitro* to explore the mechanisms of phenotypic convergence *in vivo,* the convergence of sets of NDD genes were next explored on the basis of shared behavioral effects in zebrafish mutants (**Fig. 7; SI Tables 4-5**). A comprehensive *in vivo* high-throughput, automated behavioral analysis in larval zebrafish^60^ revealed clear stratification of NDD genes based on basic arousal and sensory processing behaviors in the developing vertebrate brain (**Fig. 7A; SI Fig. 22**). Given that zebrafish brain gene expression was significantly correlated with *in vitro* human-derived mature neurons (**Fig. 7B; SI Fig. 23**), we asked whether behavioral stratification of NDD mutants in larval zebrafish can be attributed to molecular convergence. For fifteen NDD genes for which we have matched behavioral and molecular analyses, zebrafish stable mutant lines and CRISPR F0 mutants were clustered based on 24 sleep-wake and visual-startle parameters, yielding four distinct clusters of genes: set 1 (*nrxn1a, mbd5, kdm5bab*), set2 (*phf12ab, skiab, chd2, smarcc2*), set 3 (*kdm6bab, kmt5b, kmt2cab*), and set 4 (*wacab, arid1b, phf21aab, chd8, ash1l*) (**Fig. 7A; SI Data 3)**. Gene-level convergence between NDD genes in these sets was distinct, largely non-overlapping between cell- typs, and stronger in mature iGLUTs than mature iGABAs (**Fig. 7C**). Across behavioral sets, rare ASD, SCZ, and ID LoF genes were enriched primarily in iGLUTs, with all sets converging on FMRP targets, highly intolerant CNVs, and ASD variants (**Fig. 7D**). Phenotypes related to developmental delay, behavior, and motor function showed unique enrichments by set, predominately in the iGLUTs, whereas all sets were enriched for seizure, hypertonia, and abnormal skeletal muscle morphology (**Fig. 7E**). Candidate drugs predicted to reverse convergent genes (i.e., drugs with anticorrelating transcriptomic signatures) in iGLUTs and iGABAs were prioritized from the 776 cMAP^78^ drugs with matched clinical and experimental zebrafish data. Top enriched drugs included antidepressants, antipsychotics, and statins (**SI Data 2; SI Fig. 24A**). Whereas some drugs were broadly predicted to reverse convergent signatures in all four NDD gene sets (e.g., the antipsychotic perphenazine), others uniquely targeted specific sets (e.g., naltrexone in set 2 iGLUTs, sirolimus in set 3 iGLUTs, and valsartan in set 3 iGABAs). Sets 3 and 4 showed the greatest number of cMAP enrichments. By considering existing pharmacological effects of the top drugs on zebrafish behavior,^60^ some of the predicted drug reversers were shown to oppose effects on NDD related phenotypes in zebrafish (**SI Fig. 24B**). Yet, the direction of effect predicted based on transcriptomic convergence in human neurons did not always align with anti-correlating behavioral effects in zebrafish (e.g., moxifloxacin, perphenazine).

**Figure 7.**
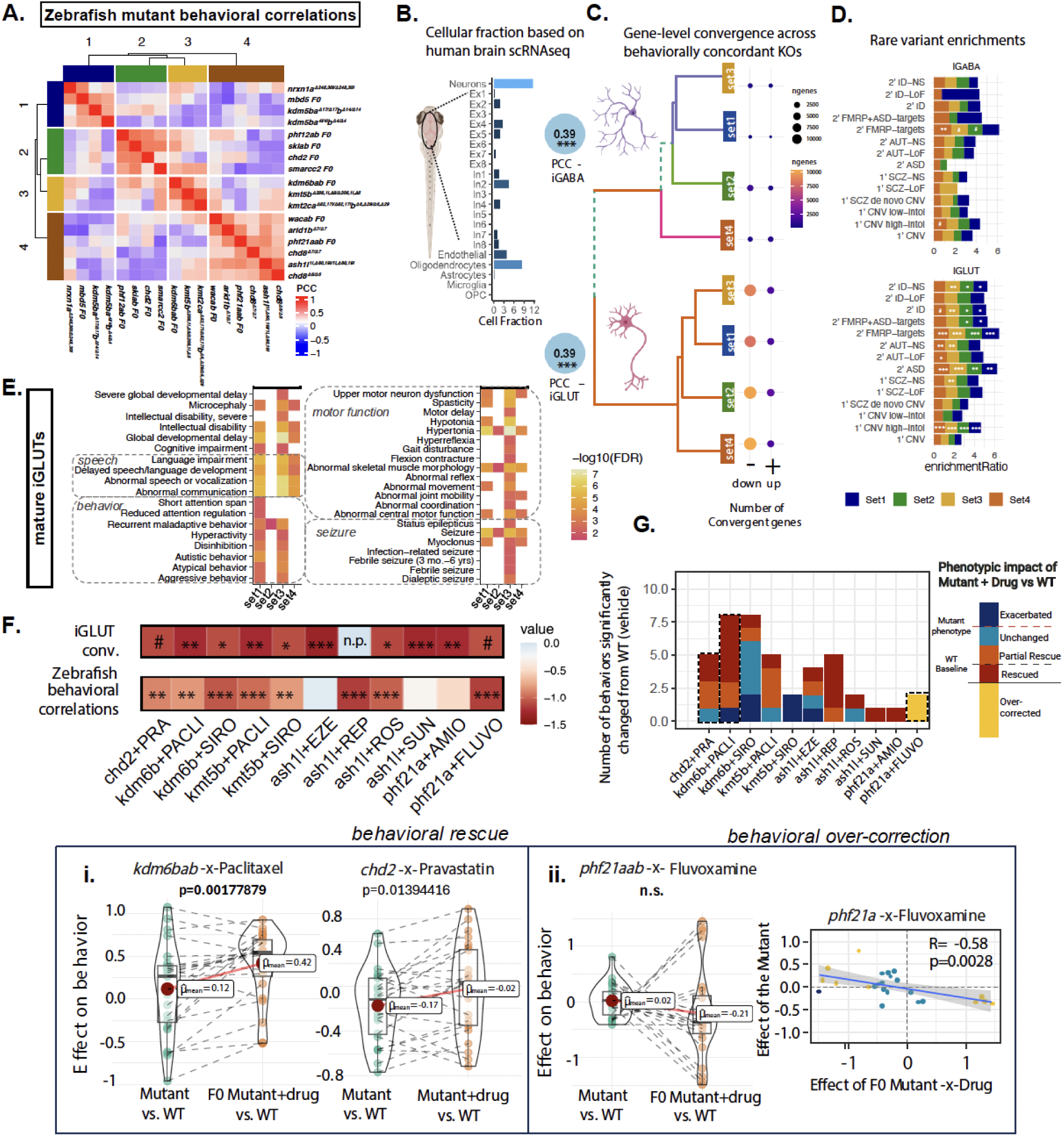
NDD gene mutants with shared behavioral phenotypes in zebrafish resolve unique and cell-type-specific gene-level convergent signatures and are rescued by predicted medications. **(A)** NDD risk genes uniquely cluster based on sleep-wake/visual-startle behavioral responses in zebrafish mutants. set 1: *nrxn1a, mbd5, kdm5bab;* set 2: *phf12ab, skiab, chd2, smarcc2*; set 3: *kdm6bab, kmt5b, kmt2cab*; set 4: *wacab, arid1b, phf21aab, chd8, ash1l*. **(B)** Gene expression in human mature iGLUTs and iGABAs correlate with expression in the zebrafish brain. Cellular deconvolution of wild-type larval zebrafish brain expression based on adult human single-cell brain reference identifying neurons as the largest proportion of cells in the fish brain. Gene expression in wild-type zebrafish brain significantly positively correlates with gene expression of mature iGLUTs (rho=0.39, Holm’s adj.P<0.001) and iGABAs (rho=0.39, Holm’s adj.P <0.001). **(C)** For each of the four behaviorally defined sets, gene-level convergence (DEGs with significant and shared direction of effect across all NDD genes within each of the four sets (FDR adjusted p_meta_<0.05, Cochran’s heterogeneity Q-test p_Het_ > 0.05)) is largely non-overlapping between mature iGLUTs and iGABAs, with unique enrichments for common psychiatric risk gene targets. Number of convergent genes that are up (+) or down (-) regulated for each NDD set are indicated. **(D)** In both iGABAs and iGLUTs, all four behavioral sets were enriched for FMRP targets. Gene targets of neurodevelopmental rare variants were only significantly enriched for convergent signatures in mature iGLUTs; behavioral set 4 uniquely significantly enriched for secondary targets of ASD loss-of-function variant and set 3 uniquely enriched for primary targets of SCZ non-synonymous variants. **(E)** In iGLUTs, NDD related behaviors were only enriched in sets 1 and 3, with enrichments for language, speech, and intellectual delays in sets 1,3 and 4. All sets were enriched for seizure and hypertonia. **(F)** Potential “rescue” drugs for these 4 phenotypic groups were selected from enrichment scores using cMAP and filtered for drugs included in a screen of 376 compounds for behavioral effects in zebrafish. Top candidates that were significantly negatively enriched for iGLUT convergence from cMAP and negatively correlated with mutant behavioral features were tested in mutant lines representative of sets 2-4. n.p. indicates that the drug repaglinide was not present in the cMAP dataset. Mutant-x-Drug combinations were as follows: *chd2^Δ7/Δ7^*-x-pravastatin; *kdm6bab* F0-x-paclitaxel; *kdm6bab* F0-x-sirolimus; *kmt5b*^Δ208,1i, Δ5/Δ208,1i, Δ5^-x-paclitaxel; *kmt5b*^Δ208,1i, Δ5/Δ208,1i, Δ5^-x-sirolimus; *ash1l*^1i, Δ60,19i/ 1i, Δ60,19i^-x-ezetimibe; *ash1l*^1i, Δ60,19i/ 1i, Δ60,19i^-x-repaglinide; *ash1l*^1i, Δ60,19i/1i, Δ60,19i^-x-rosuvastatin; *ash1l*^1i, Δ60,19i/1i, Δ60,19i^-x-sunitinib; *phf21aab* F0-x-amiodarone; *phf21aab* F0-x-fluvoxamine. **(G)** For behaviors that were significantly different between mutant+DMSO and WT+DMSO (p<0.05), we characterized the effect of the mutant-x-Drug on behavior as either (a) *exacerbated* [sig. effect mutant+Drug-v-WT > sig. effect mutant-v-WT], (b) *unchanged* [sig. effect mutant+drug-v-WT = sig. effect mutant-v-WT], (c) *partial rescue* [effect mutant+Drug-v-WT < effect mutant-v-WT], (d) *rescued* [sig. effect mutant-v-WT, no sig. effect mutant+Drug-v-WT], (e) *over-corrected* [mutant+Drug-v-WT opposite direction of sig. effect mutant-v-WT]. All drugs reversed at least one dysregulated behavior except for sirolimus in *kmt5b*. **(i)** Comparison of the magnitude of effect (beta) on behavior between the mutant+DMSO compared to mutant+Drug groups shows rescue of select behavioral features in *kdm6b* and *chd2* mutants by paclitaxel (Shapiro Wilk’s Normality p=, Student T statistic=-3.533, p=0.0017788, df=23) and pravastatin (Student T statistic=-3.533, p=0.0017788, df=23), respectively. **(ii)** the *phf21a* mutant phenotype was strongly opposed by fluvoxamine (Pearson’s correlation=-0.58, p=0.0028).

The top negatively enriched drugs for iGLUT convergence from cMAP and anti-correlating drugs predicted from a pharmaco-behavioral screen of 376 drugs in larval zebrafish were empirically tested in representative mutants from sets 2-4, which showed the strongest cMAP enrichments (**Fig. 7F**). We determined whether the phenotypic impact of mutant-x-drug combinations led to partial rescue, rescue, over-correction, or exacerbation of the mutant phenotype across significant arousal and startle behavioral parameters (**Fig. 7G**). Ten out of eleven drugs rescued at least one dysregulated behavioral parameter (**Fig. 7G, SI Fig. 24C-E**). Paclitaxel robustly rescued behavioral parameters in *kdm6bab* F0 mutants and pravastatin partially and completely rescued select parameters in *chd2^Δ7/Δ7^* mutants (**Fig. 7Gi**), including nighttime sleep bouts in *kdm6bab* F0 mutants and responses to lights-ON stimuli in *chd2^Δ7/Δ7^* mutants (**SI Fig. 24Fi-ii**). Interestingly, we also observed over-correction of the *phf21aab* F0 mutant phenotype by fluvoxamine (**Fig. 7Gii**), such as increased sleep bouts that were significantly decreased following fluvoxamine treatment (**SI Fig. 24Fiii**). Taken together, *in vivo* behavioral profiling of NDD genes in zebrafish overlaps with *in vitro*-defined convergent networks and identifies pharmacological suppressors of specific behavioral phenotypes.

## DISCUSSION

Towards empirically resolving the common pathways converged upon by NDD risk gene effects, 29 NDD genes were targeted through a pooled CRISPR-KO strategy. The molecular points of convergence across NDD risk genes varied between the cell types of the brain, being greatest in mature glutamatergic neurons, where they were enriched not just for pathways with well-established links to ASD etiology (e.g., gene regulation, synaptic biology), but also mitochondrial function^79^. While downstream effects of epigenetic NDD genes unexpectedly targeted mitochondrial genes, in fact, five percent of NDD cases meet diagnostic criteria for classic mitochondrial disorders^80^. Mitochondrial DNA mutations^81,82^, haplotypes^83^ and heteroplasmy^81,84^ have all been associated with NDD. Not only do mitochondrial mutations cause synaptic and behavioral phenotypes^85^, but multiple lines of human and animal evidence link NDDs to mitochondrial deficits and oxidative stress^10,86–91^, with neuronal and/or behavioral phenotypes reversed by antioxidant treatment^87,89–91^. Conversely, knockout of NDD genes in NPCs primarily alter neurogenesis^54,57,59^ and developmental dynamics^27,92^. Put simply, perturbations of the same NDD genes resulted in different convergent networks across cell types. This observation connects the pleiotropic nature of many NDD genes and pathophysiological evidence linking multiple cell types and distinct cellular functions to NDD.

What explains phenotypic convergence between NDD genes with distinct annotated functions? The strength of convergence was most highly correlated to common clinical associations, biological annotations, and co-expression patterns in the post-mortem brain. Critically, these factors are inter-dependent. NDD risk genes most strongly implicated in DD are enriched for expression in progenitor cells and immature neurons, and those in ASD in mature neurons^61^. Indeed, cellular identities and biological pathways are captured by patterns of gene co-expression^93,94^. Transcriptomic and epigenomic analyses of post-mortem brain from NDD cases likewise indicate convergent molecular signatures^95^ and subtypes of NDD^96^. Thus, we posit that shared clinical and phenotypic effects of distinct NDD genes in fact reflect the patterns of co-expression in the developing brain.

Personalized medicine seeks to tailor treatments to individual patients^97^; for example, cancer^98^ and monogenic disease^99^ patients with specific genetic mutations receive targeted treatments. Previous efforts to classify genes that predict NDD clinical features or treatment response applied gene ontology^4,61^ or differential neurodevelopmental KO effects *in vitro*^54^ or *in vivo*^59^. Here, we proposed to stratify risk genes based on convergent molecular impacts in human neurons. Our overarching hypothesis, in doing so, was that by resolving shared downstream gene targets between multiple NDD genes, we might inform a precision medicine-based approach that did not necessarily need to target risk genes one-at-a-time. Although convergent networks did not predict behavioral stratification of zebrafish mutants, they did inform drug prediction, with ten out of eleven drugs tested found to ameliorate at least one mutant behavioral phenotype *in vivo*. This ability to reverse, rather than prevent, a behavioral phenotype, indicates that targeting convergent networks in post-mitotic neurons may represent a clinically-actionable neurodevelopmental window that persists through symptom onset. The extent to which convergent downstream targets, whether associated with risk or resilience, can be manipulated to prevent or ameliorate NDD signatures and phenotypes warrants future investigation.

Although rare LoF NDD gene mutations tend to confer large effects in the individuals who carry them, the small effects of common variants account for much of the genetic risk for NDD at the population level^2,100^. The differences in expressivity and incomplete penetrance of high effect-size rare variants is frequently attributed to diversity across polygenic backgrounds^101^; *in vitro*, NDD gene effects are indeed influenced by the individual genomic context^27^. In psychiatry, common genetic variants are more associated with cross-disorder behavioral dimensions^102^ and rare variants with co-occurring intellectual disability^103^. Common risk variants interact with rare mutations to determine individual-level liability in ASD^104–106^, schizophrenia^107,108^, epilepsy^109^, Huntington’s disease^110^ and more^111^. Our results, highlighting that convergence downstream of NDD gene effects are enriched for cross-disorder GWAS variants and rare LoF genes, inform pleiotropy of genetic risk for psychiatric disorders. Moving forward, we argue that it is critical that empirical functional genomic studies systematically consider the impact of common and rare variants together, including screening the impact of LoF genes in hiPSC lines derived from donors with high and low polygenic risk scores^112^. Intriguingly, even susceptibility to environmental risk factors for NDD (e.g., valproic acid^113^) seems to be mediated by genetic background^114^. Deeper phenotypic characterization of NDD effects across donors will be critical in determining how complex genetic (or environmental) interactions shape cellular phenotypes, circuit function, and human behavior in the clinic.

In the post-mortem brain, NDD gene signatures are not just associated with downregulation of co-expression modules involving synaptic signalling^115^, but also upregulation of microglial and astrocyte gene modules^88,96,115–120^. The extent to which increased neuroimmune activity in NDD is a response to cellular or environmental sources of inflammation, or indicative of a role for glia cells in risk is unclear; evidence supports both possibilities. Consistent with a model of maternal immune activation during neurodevelopment^121^, glucocorticoids and inflammatory cytokines perturb the expression of psychiatric risk genes^122,123^, altering the regulatory activity of psychiatric risk loci^124^, and interfering with neuronal maturation in brain organoids^125^. Yet, *in vivo* analysis of NDD genes in zebrafish revealed global increases in microglia^60^ and *in vitro* screening in human microglia uncovered roles in endocytosis and uptake of synaptic material^126^. Indeed, given the reciprocal relationships between neuronal activity and glial function, epigenetic state, and gene expression^127–130^, it seems probable that both cell-autonomous and non-cell-autonomous effects underlie and/or exacerbate NDD gene effects.

In summary, we demonstrate that convergent effects of NDD risk genes vary between cell types. Our analyses suggest that clinical convergence between regulatory and synaptic genes in the etiology of NDD is driven more so by co-expression patterns of risk genes then direct regulation of epigenetic genes on synaptic targets. If the convergence of multifold risk genes on a smaller number of shared molecular pathways indeed explains how genetically heterogeneous mutations result in similar clinical features, then genetic stratification of cases will inform novel therapeutic targets. We predict that such individualized points of therapeutic intervention may be most effective when targeting mature glutamatergic neurons, which not only harbor the strongest convergent effects but also represent a therapeutic window that is actionable after diagnosis.

## MATERIALS AND METHODS

### Generation of neural cells

Informed consent was obtained at the National Institute of Mental Health, under the review of the Internal Review Board of the NIMH. hiPSC work was reviewed by the Internal Review Board of the Icahn School of Medicine at Mount Sinai as well as by the Embryonic Stem Cell Research Oversight Committee at the Icahn School of Medicine at Mount Sinai and Yale University. Fibroblasts were genotyped by IlluminaOmni 2.5 bead chip genotyping^131,132^, PsychChip^133^, and exome sequencing^133^; hiPSCs^133^ were validated by G-banded karyotyping (Wicell Cytogenetics) and genome stability monitored by Infinium Global Screening Array v3.0 (lllumina). SNP genotype was inferred from all RNAseq data using the Sequenom SURESelect Clinical Research Exome (CRE) and Sure Select V5 SNP lists to confirm that neuron identity matched donor. Control hiPSCs were cultured in StemFlex media (Gibco, #A3349401) supplemented with Antibiotic-Antimycotic (Gibco, #15240062) on Geltrex-coated plates (Gibco, #A1413302). Cells were passaged at 80-90% confluence with 5mM EDTA (Life Technologies #15575-020) for 3 min at room temperature (RT). EDTA was aspirated and cells dissociated in fresh StemFlex media. Media was replaced every 48-72 hours for 4-7 days until the next passage.

Transient transcription factor overexpression from stable clonal hiPSCs was used to induce control hiPSCs to iNPCs (here SNaPs)^68^, iGLUTs^69^, and iGABAs^70^. iNPCs are rapidly generated by 48-hour induction with *NGN2*^68,134^. iGLUTs are induced via transient overexpression of *NGN2*, and are >95% glutamatergic neurons, robustly express excitatory genes, and show spontaneous excitatory synaptic activity by three-to-four weeks *in vitro*^29,34,35,67,69,135–141^. iGABA neurons are induced via transient overexpression of *ASCL1* and *DLX2,* and are >95% GABAergic neurons, robustly express inhibitory genes, and show spontaneous inhibitory synaptic activity by five-to-six weeks^38,70,137,142,143^. iNPCs, iGLUTs, and iGABAs express most NDD genes, including all genes prioritized herein^67^.

We transduced hiPSCs from two control donors (553-3, karyotypic XY; 3182-3, karyotypic XX) with lentiviral *pUBIQ-rtTA* (Addgene #20342) and *tetO-NGN2-eGFP-NeoR* (Addgene #99378) for iNPCs and iGLUTs, or *pUBIQ-rtTA* (Addgene #20342), *tetO-ASCL1-PuroR* (Addgene #97329), and *tetO-DLX2-HygroR* (Addgene #97330) for iGABAs. Following transduction by spinfection at 1000g for 1 hour at 37°C, hiPSCs were subjected to 48-hour antibiotic selection (1mg/mL neomycin G418 (Thermo #10131027), 0.5µg/mL puromycin (Thermo #A1113803), and/or 250µg/mL hygromycin (Thermo, #10687010) and then clonalized by expansion from single colonies. Ultimately, clonal and inducible iNPC/iGLUT 3182-3-clone5 (XX) and iGABA 553-3-clone34 (XY) hiPSCs were validated lentiviral genome integration by PCR, doxycycline induced transcription factor expression by qPCR, and robust and consistent neuronal induction confirmed by RNA-seq and immunocytochemistry for relevant cell type markers. Analyses throughout reflect data from iGLUT 3182-3-clone5 (iNPC, d7 iGLUT and d21 iGLUT) and iGABA 553-3-clone34 (d36 iGABA).

#### iNPCs

At DIV0, 3182-3-clone5 hiPSCs were dissociated and plated at 1.5 × 10^6^ cells per well onto Geltrex-coated 6-well plates (1:250 dilution coating) in SNaP Induction Media (DIV0): DMEM/F12 with Glutamax (ThermoFisher, 11320082), Glucose (0.3% v/v), N2 Supplement (1:100, ThermoFisher, 17502048), Doxycycline (2 μg/mL; Sigma-Aldrich, D9891), LDN-193189 (200 nM; Stemgent, 04-0074), SB431542 (10 μM; Tocris, 1614), and XAV939 (2 μM; Stemgent, 04-00046) supplemented with 25 ng/mL Chroma I ROCK2 Inhibitor. After 24 hours, DIV2, cells were fed with Selection Media: DMEM/F12 with Glutamax, Glucose (0.3% v/v), N2 Supplement (1:100), Doxycycline (2 μg/mL), Geneticin (0.5 mg/mL; Thermofisher, 10131035), LDN-193189 (100 nM), SB431542 (5 μM), and XAV939 (1 μM). After 48 hours post induction (DIV2), SNaPs were dissociated with Accutase for 10 minutes at 37°C, quenched in DMEM, pelleted at 800g for 5 minutes, and replated at 1.5×10^6^ cells per well onto Geltrex-coated 6-well plates in SNaP Selection Media supplemented with Geneticin (0.5 mg/mL). After 16-18 hr (DIV3), medium was switched to SNaP maintenance Medium: DMEM/F12 with Glutamax, Penn/Strep (1:100), MEM-NEAA (1:100; Life Technologies, 10370088), B27 minus Vitamin A (1:50; Life Technologies, 12587010), N2 Supplement (1:100; Life Technologies, 17502048), recombinant human EGF (10 ng/mL; R&D Systems, 236-EG-200), recombinant human basic FGF (10 ng/mL; Life Technologies, 13256029), Geneticin (0.5 mg/mL), and Chroman I (25 ng/mL). Cells were fed every 48 hours with SNaP maintenance medium lacking Chroman I and Geneticin. Cells were dissociated and seeded weekly at a density of 1.25-1.5×10^6^ cells per well onto Geltrex-coated 6-well plates until NPC morphology was observed and persistent. Cells were expanded and cryofrozen.

#### DIV7 iGLUTs

3182-3-clone5 iNPCs were thawed and seeded at 1× 10^6^ cells per well onto Geltrex-coated 12-well plates. *NGN2* expression was induced with Doxycycline (2 μg/mL) for 24 hrs (DIV0) with antibiotic selection for 48 hrs (DIV1-3) in SNaP maintenance medium. At DIV 4 SNaPs were dissociated with Accutase, switched into Neuronal Medium: Brainphys (Stemcell, 05790), Glutamax (1:100), Sodium Pyruvate (1 mM), Anti-Anti (1:100), N2 (1:100), B27 without vitamin A (1:50), BDNF (20 ng/mL; R&D, 248-BD-025), GDNF (20 ng/mL; R&D, 212), dibutyryl cAMP (500 μg/mL; Sigma, D0627), L-ascorbic acid (200 μM; Sigma, A4403), Natural Mouse Laminin (1.2 μg/m; Thermofisher, 23017015) and seeded in Geltrex-coated (1:120 dilution coating) 12-well plates. Medium was changed every 24 hrs until DIV7 harvest.

#### D21 iGLUTs

hiPSCs were harvested in Accutase (Innovative Cell Technologies, AT-104) for 5 minutes 37°C, dissociated into a single-cell suspension, quenched in DMEM, pelleted via centrifugation for five minutes at 1000 rcf and resuspended in StemFlex containing 25 ng/mL Chroma I ROCK2 Inhibitor and 2.0 μg/mL doxycycline (DIV0), seeded 1 × 10^6^ cells per well onto Geltrex-coated 6-well plates (1:250 dilution coating), and incubated overnight at 37°C. The next day, DIV1, hiPSCs were subjected to 48-hour antibiotic selection by medium replacement with Induction Media: DMEM/F12 (Thermofisher, 10565018), Glutamax (1:100; Thermofisher, 10565018), N-2 (1:100; Thermofisher, 17502048), B27 without vitamin A (1:50; Thermofisher, 12587010), Antibiotic-Antimycotic (1:100) with 1.0μg/mL doxycycline and 0.5mg/ml Geneticin. At DIV3, cells were treated with 4.0μM cytosineβ-D-arabinofuranoside hydrochloride (Ara-C) and 1.0μg/mL doxycycline to arrest proliferation and eliminate non-neuronal cells in the culture. At DIV4 immature neurons were dissociated with Accutase and 5 units/mL DNAse I at 37°C for 7-10 min, quenched in DMEM, centrifuged for five minutes at 1,500 rpm and resuspended in 25 ng/mL Chroma I ROCK2 Inhibitor, 1.0 μg/mL doxycycline and 4.0μM Ara-C and switched to Neuron Medium: Brainphys (Stemcell, 05790), Glutamax (1:100), Sodium Pyruvate (1 mM), Anti-Anti (1:100), N2 (1:100), B27 without vitamin A (1:50), BDNF (20 ng/mL; R&D, 248-BD-025), GDNF (20 ng/mL; R&D, 212), dibutyryl cAMP (500 μg/mL; Sigma, D0627), L-ascorbic acid (200 μM; Sigma, A4403), Natural Mouse Laminin (1.2 μg/mL; Thermofisher, 23017015) and seeded 7 × 10^5^ cells per well onto Geltrex-coated (1:60 dilution coating) 12-well plates and incubated overnight at 37°C. The next day, DIV 6, Chroman I was removed from culture and Ara-C lowered to 2.0 μM with a full Neuronal medium change. At DIV 7 a full Neuronal Medium change was performed to remove doxycycline and Ara-C from culture, to allow for antibiotic resistant genes silencing. From DIV7 onwards, half neuronal medium changes were performed every 72 – 96 hrs until mature DIV 21 for harvest.

#### DIV36 iGABAs

hiPSCs were harvested in Accutase (Innovative Cell Technologies, AT-104) for 5 minutes 37°C, dissociated into a single-cell suspension, quenched in DMEM, pelleted via centrifugation for five minutes at 1000 rcf and resuspended in StemFlex containing 25 ng/mL Chroma I ROCK2 Inhibitor and 2.0 μg/mL doxycycline (DIV0), seeded 1.5-2× 10^6^ cells per well onto Geltrex-coated 6-well plates (1:250 dilution coating), and incubated overnight at 37°C. The next day, DIV1, hiPSCs were subjected to 48-hour antibiotic selection by medium replacement with Induction Media: DMEM/F12 (Thermofisher, 10565018), Glutamax (1:100; Thermofisher, 10565018), N-2 (1:100; Thermofisher, 17502048), B27 without vitamin A (1:50; Thermofisher, 12587010), Antibiotic-Antimycotic (1:100) with 1.0μg/mL doxycycline, 1.0 μg/mL puromycin (Sigma, P7255) and 250 μg/mL hygromycin (Sigma, 10687010). At DIV3, cells were treated with 4.0μM cytosineβ-D-arabinofuranoside hydrochloride (Ara-C) and 1.0μg/mL doxycycline to arrest proliferation and eliminate non-neuronal cells in the culture. At DIV5 immature neurons were dissociated with Accutase and 5 units/mL DNAse I at 37°C for 7-10 min, quenched in DMEM, centrifuged for five minutes at 1,500 rpm and resuspended in 25 ng/mL Chroma I ROCK2 Inhibitor, 1.0 μg/mL doxycycline and 4.0μM Ara-C and switched to Neuron Medium: Brainphys (Stemcell, 05790), Glutamax (1:100), Sodium Pyruvate (1 mM), Anti-Anti (1:100), N2 (1:100), B27 without vitamin A (1:50), BDNF (20 ng/mL; R&D, 248-BD-025), GDNF (20 ng/mL; R&D, 212), dibutyryl cAMP (500 μg/mL; Sigma, D0627), L-ascorbic acid (200 μM; Sigma, A4403), Natural Mouse Laminin (1.2 μg/mL; Thermofisher, 23017015) and seeded 7 × 10^5^ cells per well onto Geltrex-coated (1:60 dilution coating) 12-well plates and incubated overnight at 37°C. The next day, DIV 6, Chroman I was removed from culture and Ara-C lowered to 2.0 μM with a full Neuronal medium change. At DIV 7 a full Neuronal Medium change was performed to remove doxycycline and Ara-C from culture, to allow for antibiotic resistant genes silencing. From DIV7 onwards, half neuronal medium changes were performed every 72-96 hrs until mature DIV 36 for harvest.

### CRISPR knockout gRNA library design (Thermofisher) and validation

From the 102 highly penetrant loss-of-function (LoF) gene mutations associated with ASD (58 gene expression regulation, 24 neuronal communication genes, 9 cytoskeletal genes, and 11 multifunction genes)^4^, gene ontology and primary literature research identified 26 epigenetic modifiers specifically involved in chromatin organization, rearrangement, and modification. ASD gene expression (RNA-seq RPKM in iGLUTs) was plotted against significance of ASD association (TADA FDR Values), to ensure selection of genes with the highest expression and highest clinical association. Gene expression was confirmed across development in the brain (BrainSpan^144^), and in bulk and scRNA-seq. 21 epigenetic modifiers (*ASH1L, ASXL3, ARID1B, CHD2, CHD8, CREBBP, KDM5B, KDM6B, KMT2C, KMT5B, MBD5, MED13L, PHF12, PHF21A, POGZ, PPP2R5D, SETD5, SIN3A, SKI, SMARCC2, WAC,*) as well as two transcription factors with putative roles as chromatin regulators (*FOXP2, BCL11A*) were selected. Gene regulatory transcription factors, general transcription factors, and DNA replication genes were excluded. Three extensively studied synaptic genes (*NRXN1, SCN2A, SHANK3*) with roles in ASD were included as positive controls and three under-explored genes for ASD role in neuronal communication genes (*ANK3, DPYSL2, SLC6A1*) were also included in the library.

Individual DNA from glycerol stocks of Invitrogen™ LentiArray™ Human CRISPR Library gRNAs-PuroR (ThermoFisher, A31949) (3-4 individual gRNAs per gene, see **SI Table 1**) were prepared using GeneJET Plasmid Miniprep Kit (K0503) and pooled at an equimolar ratio and a 5-fold ratio of scramble control gRNA plasmid. Library quality was confirmed by restriction enzyme digest (10x Cutsart NEB), agarose gel purification using QIAquick Gel Extraction Kit (#28706) to check library purity, followed by Mi-seq for gRNA count distribution. Based on the abundance of gRNAs from Mis-seq, 4 NDD gene targets were highly unlikely to be resolved in the final experiments – *POGZ, PP2R5D, SHANK3, SLC6A1* – and 3 with low abundance and less likely to be resolved (*SCNA2, FOXP2, DYPSL2*).

Lentiviral Cas9v2-HygroR (Addgene, 98291) and pooled LentiArray-gRNA-PuroR CRISPR-KO library were packaged as high-titer lentiviruses (Boston Children’s Hospital Viral Core) and experimentally titrated in each cell type. Highest viable MOI was used for Cas9v2 and MOI < 0.5 for lentivirus gRNAs pool library.

#### CRISPR and gRNA delivery

Lentiviral Cas9v2-HygroR (Addgene #98291) transduction of iNPCs, day 4 (iGLUTs), or day 5 (iGABAs) occurred via spinfection (one hour at 1,000 g) and followed by 72 hr hygromycin (250 μg /mL) (except for iGABAs, which express inducible hygromycin resistance at this stage). Pooled Invitrogen™ LentiArray™ Human gRNA-PuroR CRISPR-KO Library gRNAs (ThermoFisher #A31949) (MOI 0.3-0.5) were transduced via spinfection three days prior to harvest (e.g., d4 for D7 iGLUTs, d18 for D21 iGLUTs, d33 for d36 iGABA), with fresh medium containing puromycin (1 μg/mL) added 16-24 hours post transduction of gRNAs. For mature iGLUTs and iGABAs, as doxycycline was removed from medium at DIV7, and by DIV18 neurons had lost transcription factor linked antibiotic resistance, at 24 hours post-transduction (DIV19 or DIV34) puromycin (1 μg/mL) and hygromycin (250 μg /mL) were added to media for 48-hr antibiotic selection prior to harvest.

### Dissociation of different neural cell types to single cells for scRNAseq assays

Cells were dissociated 72 hrs post gRNA library delivery for single cell sequencing, as iNPCs, DIV7 and DIV21 iGLUTs, or DIV36 iGABAs as follows: iNPCs and DIV7 iGLUTs were dissociated in accutase for 5min @37°C, washed with DMEM/10%FBS, centrifuged at 1,000xg for 5 min, gently resuspended, and counted.

DIV21 iGLUTs and DIV36 iGABAs were dissociated with papain. Papain was pre-warmed (39°C) for 30 minutes in HBSS (ThermoFisher, 14025076), HEPES (10 mM, pH 7.5) EDTA (0.5 mM), Papain (0.84 mg/mL; Worthington-Biochem, LS003127). The cells were washed with PBS-EDTA (0.5 mM) and 300 uL of papain solution and 5 units of DNAse I was added per well of 12-well plate and incubated at 37°C for 10-15 minutes, 125 rpm. Dissociation was quenched with DMEM-10%FBS. Detached neurons were broken by gentle manual pipetting, pelleted at 600 g for 5 minutes, resuspended in DMEM-10%FBS, filtered through a cell strainer and counted and submitted for 10X sequencing.

Cells were loaded into 10X in four lanes per cell type, targeting 20,000 cells per lane for a total of ∼80,000 targeted cells per cell type. scRNA-seq was performed at Yale Genomics Core with the 10X single cell 5’ v2 HT with CRISPR barcode kit.

### Bulk RNAseq and CRISPR-editing efficiency evaluation

The H1 hESC line with iCas9 (NIHhESC-10-0043), generously provided by the Huangfu Lab, was used to assess the editing efficiency of the gRNAs^77,145^ and conduct the mitochondrial pooled and arrayed experiments. NPCs were generated using the dual SMAD inhibition approach per the STEMdiff SMADi Neural Induction Kit protocol (STEMCell Technologies, #08581). To validate gene KO, NPCs were transduced with LV particles carrying four gRNAs per target gene. After 48 h of selection with 1 µg/mL puromycin, Cas9 expression was induced by adding dox at 2 µg/mL for 72 h. Following induction, cells were collected for bulk RNA-seq. Total RNA was extracted using TRIzol™ reagent (Invitrogen). PolyA RNA-seq library preparation and sequencing were conducted at the Yale Center for Genomic Analysis (YCGA). Raw fastq files were quality-checked by FastQC, then mapped to human genome reference hg38 (STAR^146^). GRNA targeted-loci for each sample were extracted (SAMtools^147^). Variation/small insertion/deletion at site of interest and mutation efficiency at corresponding loci was called (CrispRVariants R package^148^), after excluding possible germline variants from Cas9-non-induced samples.

### Proliferation and neurogenesis analysis

For proliferation analysis using Ki-67, NPCs were seeded into 24-well plates and either treated with doxycycline (induced) to activate Cas9 or left untreated (uninduced). The cells were cultured for 7 days, representing approximately three NPC generations. On day 7, cells were collected, and ∼1 × 10^6^ cells were stained with Ki-67-FITC (#130-117-803, Miltenyi Biotec) using the Foxp3/Transcription Factor Staining Buffer Set (#00-5523, Invitrogen), following the manufacturer’s protocol.

To evaluate the effects of gene KOs on neurogenesis and gliogenesis, transduced NPC-iCas9 lines were spontaneously differentiated into human cortical neurons and glial cells. Briefly, 1 × 10^6^ cells were seeded in GelTrex-coated (1:5) 6-well plates and cultured in complete neuronal media containing BrainPhys™ Neuronal Medium, Glutamax (100X), Sodium Pyruvate (100 mM), B-27 (-RA) supplement (50X), N2 (100X), Anti-Anti (100X), Natural Mouse Laminin (1 mg/ml), dbcAMP (500 mg/ml), L-Ascorbic Acid (200 µM), BDNF (20 µg/ml), and GDNF (20 µg/ml). Media was refreshed every three days. On day 25, cells were collected and stained for FACS analysis using surface markers previously described^149^ to differentiate NPCs (CD184+/CD44-/CD24+), neurons (CD184-/CD44-/CD24+), and glia (CD184+/CD44+). CD271, a marker for mesenchymal stem cells, was excluded from the original panel as NPCs were pre-purified via FACS using CD133+/CD184+/CD271-markers before differentiation. A minimum of 50,000 cells per gate were acquired using a BD LSRFortessa™ Cell Analyzer at the Yale Flow Cytometry Core. Flow cytometry data were analyzed using FlowJo™ v10.10 Software (BD Life Sciences).

All statistical analyses for flow cytometry assessment were conducted using GraphPad Prism version 9.5.1 (528) for macOS (GraphPad Software, San Diego, CA). Each well was treated as an independent replicate. Differences between knockout (induced) and control (uninduced) groups were assessed by comparing the mean fluorescence intensity (MFI) of the target fluorophore using an unpaired t-test with Welch correction to account for individual group variance. Multiple comparisons were corrected using the False Discovery Rate (FDR) method with a two-stage step-up procedure (Benjamini, Krieger, and Yekutieli) at an FDR threshold of 5%.

### FACS Analysis of Mitochondrial Membrane Potential and CRISPR Screen Read-out via Amplicon Sequencing

For our mitochondrial assays we used a nearly identical library (same backbone, guide density, and control set) screened exclusively in the H1 inducible Cas9 (H1-iCas9) hPSC line. Mitochondrial inner membrane potential (Δψm) was measured in H1-iCas9, following differentiation to NPCs or iGlut on day 21. Cells were harvested, counted, and aliquoted at 1 × 10^6 cells per sample. JC-1 dye (MitoProbe™ JC-1 Assay Kit; Invitrogen #M34152) was dissolved in DMSO at a stock concentration of 200 µM and added to each sample to achieve a final concentration of 2 µM, then incubated for 30 min at 37 °C in 5% CO₂. A 50 µM CCCP control was included to induce complete mitochondrial depolarization. After staining, cells were washed once in their respective culture medium, resuspended in FACS buffer (Invitrogen eBioscience Staining Buffer #00422226), and analyzed immediately on a Thermo Fisher “Bigfoot” spectral cell sorter using 488 nm excitation with 525/50 nm (FITC) and 585/40 nm (PE) emission filters. Debris and doublets were excluded by forward/side scatter gating, and CCCP-treated samples were used to define FITC and PE gates. Approximately 1 × 10^6 events per sample were recorded. Cells were then pelleted (300 × g, 5 min) and genomic DNA extracted using the Qiagen DNeasy Blood & Tissue Kit (#69504).

UMI-tagged amplicon libraries were generated in three PCR steps. In PCR-1, genomic DNA was amplified with Platinum™ II Hot-Start PCR Master Mix (Invitrogen, #14000012) and UMI-containing primers (Forward: 5′-ACACTCTTTCCCTACACGACGCTCTTCCGATCTACGTGACGTAGAAAGTAATAATTTCTTGGGT-3′; Reverse: 5′-GTGACTGGAGTTCAGACGTGTGCTCTTCCGATCTN(25)NNNNNNNNNACTCGGTGCCACTTTTTCAA-3′) under the following conditions: 94 °C for 2 min; 4–6 cycles of 98 °C for 5 s, 60 °C for 15 s, 60 °C for 30 s. The resulting ∼180 bp products were purified and concentrated using the Zymo DNA Clean & Concentrator-5 kit (#D4013) and eluted in 10 µL nuclease-free water (Thermo Fisher #AM9938). In PCR-2, purified product was amplified with adaptor primers (Forward: 5′-ACACTCTTTCCCTACACGACGCTCTTCCGATCT-3′; Reverse: 5′-GTGACTGGAGTTCAGACGTGTGCTCTTCCGATCT-3′) for 22 cycles under identical cycling conditions in a ∼20 µL reaction. A seven-cycle indexing PCR (PCR-3) was performed by the sequencing facility Yale Center for Genome Analysis (YCGA) prior to sequencing. Final libraries were sequenced on an Illumina NovaSeq platform (paired-end 150 bp, 5 million reads per sample).

Flanking sequences on both sides of each gRNA were trimmed using BBDuk, and reads were then mapped to gRNA reference sequences and counted using MAGeCK^150^. Raw counts for each gRNA were normalized to counts of scrambled gRNA. Abundance of each target gene was then calculated by summing of all gRNAs targeting that gene. Log2-transformed fold changes of gRNA-targets abundance were compared between PE-high samples and FITC-high samples.

### Immunostaining

*Cells* were fixed with fixative solution (4 % sucrose and 4 % paraformaldehyde prepared in Dulbecco’s Phosphate Buffered Saline (DPBS)) for 10 min at room temperature (RT). Following this, cells were washed twice with DPBS and incubated in blocking solution (2% normal donkey serum prepared in DPBS) supplemented with 0.1% Triton for two hours at RT. After this, cells were incubated overnight at 4 °C in the primary antibody solution prepared in blocking solution. Cells were washed three times with DPBS, incubated at RT in secondary antibody prepared in blocking solution, then washed three times with DPBS. In the second wash, cells were incubated in DBPS supplemented DAPI (Sigma D9542,1 μg/mL) for 2 min at RT.

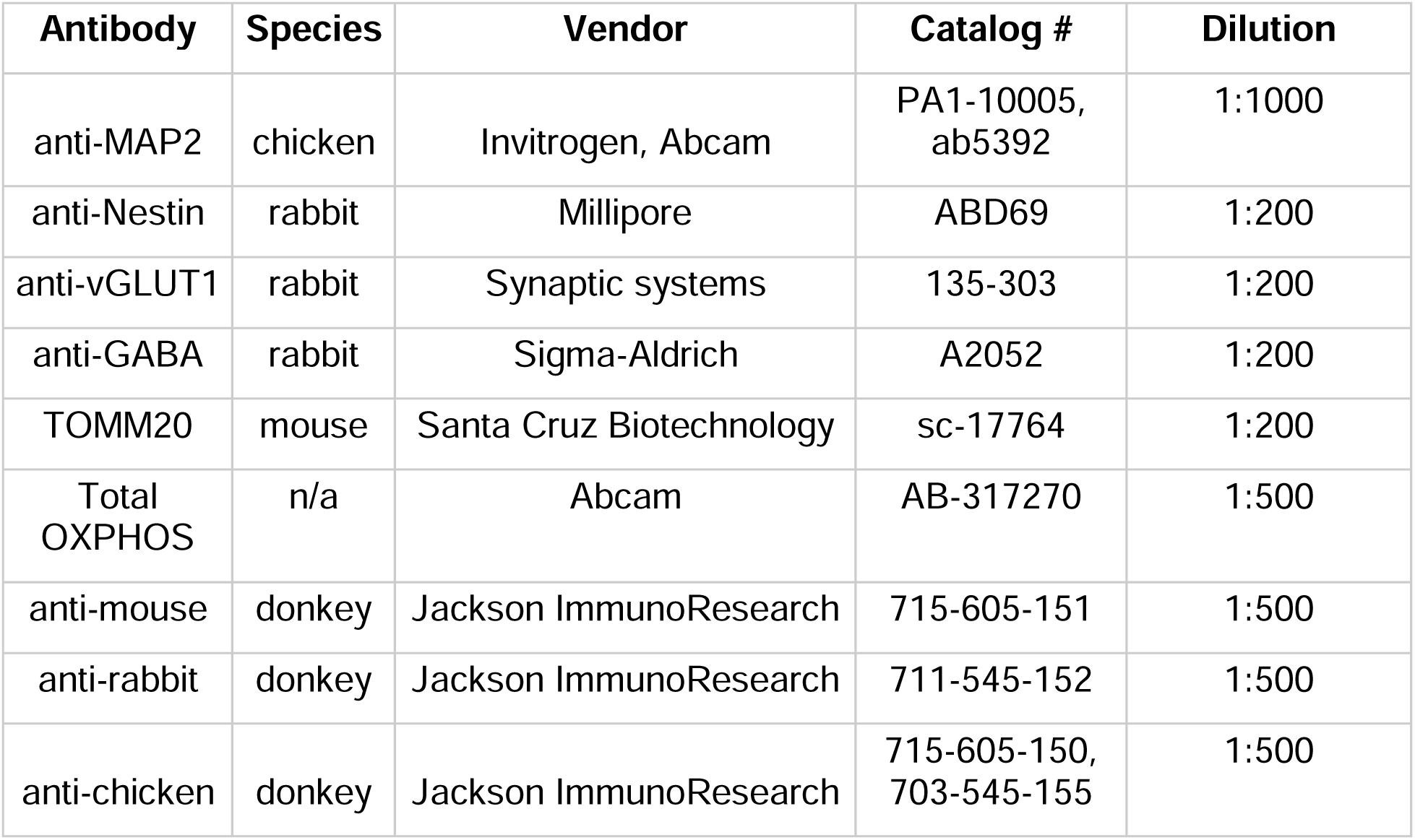

Fixed cultures were acquired using a DragonFly Confocal Dual Spinning Disk confocal, at 60x magnification and 1.4 numerical aperture. All images were acquired with a fixed laser intensity and exposure time across experimental conditions. Four images were acquired per well, and 4-10 wells were acquired per experimental condition. Each well represents a biological replicate and statistical datapoint. Therefore, each replicate represents hundreds of μm^2^ of neuronal area and tens of thousands of individual mitochondria.

Mitochondria morphology features were determined using the Surface module of Imaris 10.2. Likewise, OXPHOS complex features were determined using the surface module of Imaris 10.2. The Volume, Area and Sphericity features of the Surface modules were selected for analysis. Mitochondria networking features were determined using published, open-source methods^179^. A one-way ANOVA with a Šidák’s multiple comparisons test was performed on data on GraphPad Prism 10.

To validate robustness and sensitivity of the microscopy assay, we treated D14 iGluts overnight with carbonyl cyanide 4-(trifluoromethoxy) phenylhydrazone/FCCP (Sigma-Aldrich, SML2959) at 5 μM, 10 μΜ and 50 μΜ doses. Following this, we conducted the immunostaining, mitochondrial structural analysis and statistical analyses outlined above.

### Seahorse XF Mito Stress Test

Day 5 iGLUTs were plated at 1.65 × 10 cells/well in XF24 microplates (Agilent, 100777-004) and cultured to day 21. One hour prior to measurement, growth medium was removed, leaving 50 µL per well, and replaced with 1 mL of pre-warmed Seahorse XF DMEM (Agilent, 103575-100) supplemented with 25 mM glucose (Agilent, 103577-100) and 0.23 mM pyruvate (Agilent, 103578-100). Plates were equilibrated for 1 h at 37 °C in a non-CO₂ incubator. Immediately before the assay, the medium was replaced with 500 µL of fresh assay buffer. Oxygen-consumption rate (OCR) was recorded on a Seahorse XFe24 Analyzer (Agilent) using the standard Mito Stress Test. The program consisted of three sequential injections—1.5 µM oligomycin (Sigma-Aldrich, 75351), 1.5 µM carbonyl cyanide 4-(trifluoromethoxy) phenylhydrazone/FCCP (Sigma-Aldrich, C2920), and a mix of 0.5 µM rotenone (Sigma-Aldrich, R8875) + 0.5 µM antimycin A (Sigma-Aldrich, A8674)—separated by four measurement phases (baseline plus post-injection 1–3). Each phase comprised three cycles of 3 min mixing, 2 min waiting, and 3 min measurement. After the assay, cells were lysed using M-PER™ Mammalian Protein Extraction Reagent (ThermoFisher, 78501) supplemented with cOmplete™ Mini Protease Inhibitor Cocktail (Sigma-Aldrich, 11836153001) and PhosSTOP™ (Sigma-Aldrich, 4906845001), according to the manufacturer’s instructions. Total protein concentrations were determined using the Pierce™ Dilution-Free™ Rapid Gold BCA Protein Assay (ThermoFisher, A55860), and OCR values were normalized to total protein content.

### CRISPR organoid assays

H1-hESC-iCas9 cells were transduced with a pooled gRNA library containing four gRNAs per target gene, with 20% of the library comprising non-targeting gRNAs. Following selection with 1 µg/mL puromycin, the established cell line was used to generate cortical organoids following a well-established protocol^151^ with slight modifications. In brief, embryoid bodies (EBs) were generated using AggreWell plates (Stemcell Technologies) according to the manufacturer’s instructions. Once formed, EBs were transferred to ultralow-attachment 10 cm plates (Corning) for further culture. Patterning was initiated using StemFlex base media (A3349401, Gibco) supplemented with 100 nM LDN193189 (x) and 10 µM SB431542 (x). The media was refreshed daily. Organoids were cultured on an orbital shaker at 53 rpm for the duration of the protocol. On Day 6, the patterning media was replaced with growth media; Neurobasal A medium (10888022, Gibco), 1× GlutaMAX (35050061, Gibco), and 1× B27 (12587010, Gibco), supplemented with 20 ng/mL FGF (PeproTech) and 20 ng/mL EGF (PeproTech). On Day 14, Cas9 expression was induced by treating the organoids with 2 µg/mL doxycycline (Sigma-Aldrich) for 72 hours. From Day 25, FGF and EGF were replaced with 20 ng/mL BDNF (PeproTech) and 20 ng/mL NT-3 (PeproTech). Media changes were performed every other day. Starting from Day 42, organoids were maintained in growth media without additional supplements. Media was refreshed 2–3 times per week.

The organoids were maintained in culture for ∼80 days, at which point five organoids from three biological replicates were collected for DNA extraction using the DNeasy Blood & Tissue Kit (#69504, Qiagen). Extracted DNA was subjected to PCR amplicon sequencing with unique molecular identifiers (UMIs) using a three-step PCR protocol. In the first step (PCR-1), UMI-containing primers (5’-ACACTCTTTCCCTACACGACGCTCTTCCGATCTACGTGACGTAGAAAGTAATAATTTCTTGGGT-3’) and (5’-GTGACTGGAGTTCAGACGTGTGCTCTTCCGATCTN(25252525)NNNNNNNNNACTC GGTGCCACTTTTTCAA-3’) were used for 4 cycles. PCR-2 utilized adaptor primers (5’-ACACTCTTTCCCTACACGACGCTCTTCCGATCT-3’) and (5’-GTGACTGGAGTTCAGACGTGTGCTCTTCCGATCT-3’) for 22 cycles. PCR-3, performed by the sequencing facility, added sample-specific indexing in 7 additional cycles. The prepared libraries were sequenced on a NovaSeq platform with paired-end 150 bp reads, generating 10 million reads per sample at the Yale Center for Genomic Analysis (YCGA).

Fragments amplified by PCR were sequenced on NovaSeq 6000 sequencer pair end at 150bp with ∼10 million reads per sample. Flanking sequence on both side of gRNAs were trimmed using BBDuk, and reads were then mapped to gRNA reference sequences and counted using MAGeCK package^150^. Raw counts for each gRNA were normalized to counts of scrambled gRNA. Abundance of each gRNA-target genes were then calculated by sum of all gRNAs targeting that gene after excluding gRNAs with low KO-efficiency (<5%). Average Log2-transformed fold change of gRNA-targets abundance were compared between doxycycline-induced versus uninduced samples on day 77 samples.

### Analysis of single-cell CRISPRko screens in NPCs, DIV 7, DIV 21 iGLUTs and DIV 36 iGABAs

mRNA sequencing reads were mapped to the GRCh38 reference genome using the *Cellranger* Software. To generate count matrices for GDO (gRNA) libraries, the kallisto indexing and tag extraction (kite) workflow were used. Count matrices were used as input into the R/*Seurat* package^152^ to perform downstream analyses, including QC, normalization, cell clustering, GDO demultiplexing, and covariate regression^71,153^.

Normalization and downstream analysis of RNA data were performed using the Seurat R package (v.5.1.0), which enables the integrated processing of multimodal single-cell datasets. CRISPR-screen experiments in each cell-type were processed independently. Within each cell-type, ∼100-80,000 cells were sequenced across 4 lanes. gRNA and RNA UMI feature counts were filtered removing the top and bottom decile of cells based on distribution of counts in each cell-type. The percentage of all the counts belonging to the mitochondrial, ribosomal, and hemoglobin genes calculated using Seurat::PercentageFeatureSet were filtered with cell-type specific thresholds, given the relatively high proportion of mitochondrial genes expressed in neurons. Mitochondrial, ribosomal, and hemoglobin genes as well as MALAT1 were removed (^RP[SL][[:digit:]]|^RPLP[[:digit:]]|^RPSA|^HB[AEGQ][[:digit:]]|^HB[ABDMQ]|^MT-|^MALAT1$). Lowly expressed genes, those that had at fewer than 2 read counts in 90% of samples were also removed. Hashtag and guide-tag raw counts were normalized using centered log ratio transformation, where counts were divided by the geometric mean of the corresponding tag across cells and log-transformed. gRNA demultiplexing was performed using the Seurat::MULTIseqDemux function for each lane individually and then counts were merged across lanes (**SI Fig. 3B**). In NPCs, 94,363 cells were retained after filtering and removal of negatively assigned cells with 62,7% classified as doublets and 37.3% classified as singlets. In DIV7 and DIV21 iGLUTs, 57,685 and 31,473 cell were retained with 34% and 9.8% doublets and 66% and 90.2% singlets respectively. In DIV35 iGABAs, 64,462 cells were retained with 48.3% doublets and 51.7% singlets. For all downstream analysis only cells with “singlet” gRNA classification were used (26,549-38,097 cells per experiment) (**SI Fig. 4C-E**). Number of singlet cells by gRNA per cell-type shown in **SI Fig. 6AB**.

### Cell-type specific population heterogeneity correction

Gene-expression based clustering was largely driven by cellular heterogeneity, cell quality, and sequencing lane effects. gRNA identity was not correlated with these covariates (**SI. Fig. 7**), so we adjusted for transcriptomic variability arising from cellular heterogeneity by applying maturity and cellular subtype scores across both perturbed and non-perturbed cells. First, variation related to cell-cycle phase of individual cells was accounted for by assigning cell cycle scores using Seurat::CellCycleScoring which uses a list of cell cycle markers^154^ to segregate by markers of G2/M phase and markers of S phase. Second, to address variance due to cellular heterogeneity within a single experiment, we adapted the method applied by Seurat::CellCycleScoring to calculate a “Maturity. Score” and “Subtype.Score” for each cell based on cellular subtype (more variable in mature GABAergic neurons) and developmental time-point specific markers (mora variable in NPCs and immature iGLUTs) (**SI Table 2-3**). Cells with outlier maturity scores and subtype scores were removed from downstream analyses. RNA UMI count data were then normalized, log-transformed and the percent mitochondrial, hemoglobulin, and ribosomal genes (markers of cell quality), lane, cell cycle scores (Phase), and maturity scores regressed out using Seurat::SCTransform. The scaled residuals of this model represent a ‘corrected’ expression matrix, that was used for all downstream analyses.

Although demultiplexing assigned the correct guide identity to each cell, to remove “false positives” whereby gRNAs were assigned but gene expression was unperturbed, the transcriptomes of gRNA clusters were evaluated relative to scramble gRNAs, ensuring that cells assigned to a guide-tag identity class demonstrated successful perturbation of the targeted NDD gene. To remove subsequent “false negatives”, whereby a successful CRISPR-KO may not result in significant down-regulation of the targeted gene^71^ yet still achieve an overall transcriptomic profile distinct from scramble populations, we performed ‘weighted-nearest neighbor’ (WNN) analysis to assign clusters based on both guide-tag identity class and gene expression^72^. To identify successfully perturbed cells, the transcriptomes of gRNA clusters were compared to Scramble-gRNA control clusters by differential gene expression analysis (Wilcoxon Rank Sum) comparing each cluster to all other clusters. Non-targeting WNN clusters and KO gRNA WNN clusters were filtered by setting a quantile base average expression threshold of target genes based on the distribution of target gene average expression across all other clusters. Clusters were the collapsed by gRNA identity; gRNAs with less than 75 cells were removed from analysis. These cells were then used for downstream differential gene-expression analyses^155^. For each cell-type individually, single-cell gene expression matrices were PseudoBulked using scuttle::aggregateAcrossCells function across lanes (4 pseudo-bulk samples per perturbation), lowly-expressed genes were removed (leaving 18-22,000 genes) followed by edgeR/limma differential gene expression analysis. Concordance between Wilcox-rank sum differential gene expression analysis using single-cell data and limma:voom using PseudoBulked data was assessed for each gene.

Altogether, Wilcoxon Rank Sum was applied to measure NDD gene knockdown from single-cell DEG analysis. Given the concordance between the DEG results using single-cell Wilcox and pseudo-bulk limma:voom (**SI Fig. 6C**), all main figure and all SI figures thereafter applied pseudobulked data analyzed with limma.

To validate whether the high correlation within cell type was due to exactly the same scramble control cells, we re-performed DEGs using random selection of subset of scramble cells for each cell type (**SI Fig. 8**). Briefly, for each gene, 50% (if number of pseudobulked sample cells > 50) or 80% (if number of pseudobulked sample cells < 50) of scramble cells were randomly selected using sample function from R. DEGs were then performed as described above using limma/dreamlet package between KOs and subset of scrambles different among genes. The process was repeated three times to avoid random selection bias and median of each gene logFC was used as the final logFC. Average overlap of random scramble cells across different genes is approximately 50%.

### Meta-analysis of gene expression across perturbations^73^

Across NDD KOs, DEGs were meta-analyzed (METAL^156^), and “convergent” genes were defined as those with significant and shared direction of effect across all NDD gene perturbations and with non-significant heterogeneity (FDR adjusted p_meta_<0.05, Cochran’s heterogeneity Q-test p_Het_ > 0.05). To test convergence between NDD-KOs, meta-analyses were performed across all possible combinations of 2-5 KO perturbations with and without sub-setting for those shared across cell types (>40,000 combinations across cell-types) (**SI Data 1**).

### Bayesian Bi-clustering to identify Convergent Networks^73^

Across NDD KOs, convergent networks were generated by Bayesian bi-clustering^157^ and undirected gene co-expression network reconstruction from the NDD KOs. Not constrained by statistical cut-offs, and able to capture the effect of more lowly expressed genes, convergent networks may be a more sensitive measure of convergence. Networks were built based on bi-clustering (BicMix)^158^ using log2CPM expression data from all the replicates across each of the NDD gene sets and Scramble gRNA jointly. We performed 40 runs of BicMix on these data and the output from iteration 400 of the variational Expectation-Maximization algorithm was used. Target Specific Network reconstruction^159^ was performed to identify convergent networks across all possible combinations of the 9 NDD gene KO perturbations shared across cell-types (n=502 combinations/cell-type) and randomly sampled combinations of 2-21 KO perturbations without sub-setting for those shared across cell types (n=1400-2300 combinations).

### Influence of Functional Similarity on Convergence Degree

To test the influence of functional similarity and brain co-expression between KOs on convergence and compare the degree of convergence between the same KOs in different cell-types we established two methods for defining and measuring convergence. First, gene-level convergence using meta-analysis as described above, with the strength of convergence for each set defined as ratio of convergent genes to the average number of DEGs.

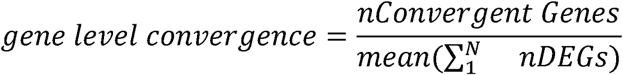

Second, network-level convergence based on undirected network reconstruction from Bayesian bi-clustering as described above. Bi-clustering identifies co-expressed genes shared across the downstream transcriptomic impacts of any given set of KO perturbations, thus, the resolved networks are the transcriptomic similarities between distinct perturbations (convergence). We calculated the “degree of convergence” for each network based on previously described metric^73^. Briefly, convergence scores are based on (1) network connectivity as defined by the sum of the clustering coefficient (Cp) and the difference in average length path (Lp) from the maximum average length path resolved across all possible sets [(max)Lp-Lp] and (2) similarity of network genes based on biological pathway membership scored by taking the sum of the mean semantic similarity scores ^160^ between all genes in the network and (3) minimum percent duplication rate across 40 runs. Duplication thresholds are network-dependent and a metric of confidence in the connections.

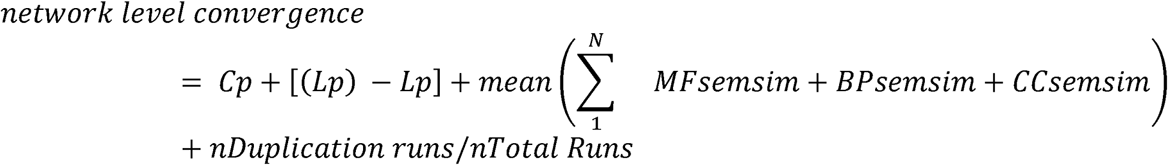

Functionally similarity scores across the NDD KO genes represented in each set was calculated using (1) Gene Ontology Semantic Similarity Scores: the average semantic similarity score based on Gene Ontology pathway membership within Biological Pathway (BP), Cellular Component (CC), and Molecular Function (MF) between NDD genes in a set^160^ and (2) brain expression correlation (BEC) score: based on the strength of the correlation in NDD gene expression in the CMC (n=991 after QC) post-mortem dorsa-lateral pre-frontal cortex (DLPFC) gene expression data,.

We performed Pearson’s correlation analysis (Holm’s adjusted P) on similarity scores and the degree of network convergence to determine the influence of the similarity of the initial KO genes on downstream convergence. We compared the average strength of convergence across cell-types using a parametric Welch’s F-test and pairwise Games-Howell test.

### Enrichment analysis of convergence for risk loci using MAGMA

We intersected cross cell-type perturbation specific and cross perturbation cell-type-specific gene-level convergence with genetic risk of psychiatric and neurological disorders/traits [attention-deficit/hyperactivity disorder (ADHD)^161^, anorexia nervosa (AN)^162^, autism spectrum disorder (ASD)^2^, alcohol dependence (AUD)^163^, bipolar disorder (BIP)^164^, cannabis use disorder (CUD)^165^, major depressive disorder (MDD)^166^, obsessive-compulsive disorder (OCD)^167^, post-traumatic stress disorder (PTSD)^168^, and schizophrenia (SCZ)^169^, Cross Disorder (CxD)^170^, Alzheimer disease (AD)^171^, Parkinson disease (PD)^172^, amyotrophic lateral sclerosis (ALS)^173^, Tourette’s^174^, migraine^175^, chronic pain^176^, and neurotic personality traits^177^ GWAS summary statistics] using multi-marker analysis of genomic annotation (MAGMA)^65^. SNPs were mapped to genes based on the corresponding build files for each GWAS summary dataset using the default method, snp-wise = mean (a test of the mean SNP association). A competitive gene set analysis was then used to test enrichment in genetic risk for a disorder across gene sets with an FDR<0.05.

To test if observed effects were due to the differential size of the gene sets for each GWAS or owing to the fact that DEGs are more likely to include neural genes, which are more likely to be associated with brain disorder, GWAS sets were filtered for genes expressed in each cell-type prior to enrichment testing and enrichment tests were performed after randomly down-sampling GWAS Gene Sets to 100, 250, 500, 750, and 1000 genes (**SI Fig. 9**), performed ten times within each set size (i.e., 50 tests for each GWAS).”

### Over-representation analysis, functional enrichment annotation, and biological theme comparison of convergence

To identify pathway enrichments unique to individual KOs, convergent genes, and convergent networks based on zebrafish behavioral subgroups (see zebrafish methods below), we performed biological theme comparison and GSEA using ClusterProfiler^178^. Using FUMAGWAS: GENE2FUNC, the 102 ASD genes were functionally annotated and overrepresentation gene-set analysis for each convergent gene set was performed^179^. Using WebGestalt (WEB-based Gene SeT AnaLysis Toolkit)^180^, over-representation analysis (ORA) was performed on all convergent gene sets against publicly available genset lists GeneOntology, KEGG, DisGenNet, Human Phenotype Ontology, and a curated gene list of rare-variant targets associated with ASD,SCZ, and ID^67^.

### Random forest prediction model of convergence strength

To determine how well functional similarity between KOs can predict gene-level and network-level convergence we trained a random forest model^75^ (randomForest package in R) for each type of convergence, evaluated the model in an independent internal dataset, and validated the model in an external CRISPRa activation screen^73^. Data from randomly tested gene combinations (2-5 KO sets at the gene level and 2-10 KO sets at the network level) tested across cell-types were randomly down-sampled into a training set (70%) and testing set (30%) – all with comparable proportions of data by cell-type. The random forest model was trained with bootstrap aggregation using C.C, M.F, B.P semantic similarity scores, brain expression correlation, number of genes, and cell-type as predictors. The Random Forest linear regression model was evaluated in the testing data by comparing actual values to predicted values, estimating the root mean squared error and performing Pearson’s correlations. Predictor models were validated using an external dataset of 10 CRISPR-activation perturbations of SCZ common variant target genes with multifunctional annotations broadly grouped as signaling/cell communication (*CALN1, NAGA, FES, CLCN3, PLCL1*) and epigenetic/regulatory (*SF3B1, TMEM219, UBE2Q2L, ZNF804A, ZNF823*)^73^, and assessed based the root mean squared error and Pearson’s correlation between actual and predicted convergence strength.

### LNCTP in silico model

To investigate the perturbation of ASD genes in silico, we adapt the Linear Network of Cell-Type Phenotypes (LNCTP) model^76^ to predict the effects of changes in gene expression in the prefrontal cortex, across neuronal and non-neuronal cell-types. The LNCTP is defined as an energy model representing the joint distribution of a collection of phenotypes of interest conditioned on the genotype. Since we are interested primarily in the effects of gene expression perturbations on the expression of other genes, we use only the imputation segment of the LNCTP model (excluding the prediction of higher-order phenotypes and cell-cell interactions).

The probabilistic model for the imputation-based LNCTP may be expressed as:

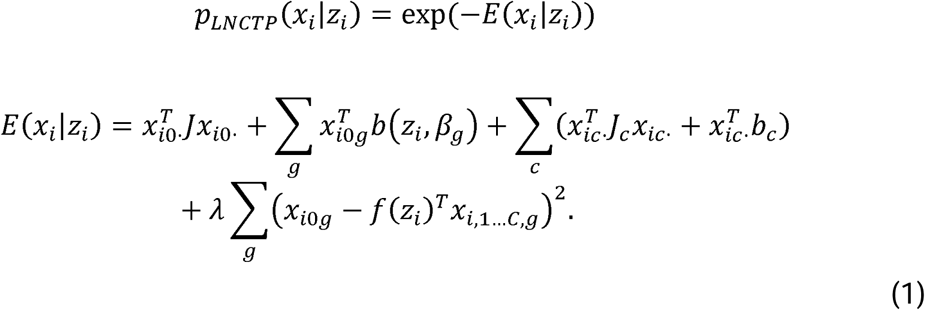

Here, *z_i_* represents the genotype of individual *i*, and *x_i_* represents bulk and cell-type specific gene expression from individual *i*. We further index the gene expression by *C* cell-types (which are here: Excitatory Neurons, Inhibitory Neurons, Oligodendrocytes, Astrocytes, Oligodendrocyte Precursor Cells, Endothelial Cells and Microglia), which will be denoted *x*_1_, *x*_2_,…*x*_c_, and we will use *x*_0_ to denote the bulk expression. The variables *f*_1_…*c* represent the estimated cell-fractions in the bulk observations (predicted from the genotype, *z*). The parameters of the model are θ = {*β*_1…*G*,_ *J*_1…*G*_} and *λ* acts as a hyperparameter. The parameters *β*_1…*G*_ and *J*_1…*G*_ reflect the gene specific expression biases and pairwise interactions respectively, whose non-zero elements are determined by the sparsity structure arising from eQTLs and Gene Regulatory Network (GRN) linkages respectively; the non-zero elements of *J_c_* occur only between genes connected in the GRN of cell-type *c*.

Further details on the training of the model in Eq. (1) can be found in^76^; here, we outline the specific differences in the training for the purposes of our analysis. As in^76^, we use genetics and expression data from post-mortem PFC samples from the PsychENCODE consortium. However, we group together samples from all higher-order phenotypes during training (control (CTR), schizophrenia (SCZ), bipolar disorder (BPD) and autism spectrum disorder (ASD)), and split the data into three partitions of size 760, 100 and 100 for training, validation and testing respectively (each including samples from all higher-order phenotypes). Further, we include all 29 CRISPR targeted genes, 102 NDD genes^61^, Transcription Factors^76^ and neuropsychiatric TWAS-selected genes^76^, and the top 100 up and down regulated CRISPR convergent genes in iGLUT and iGABA cells (400 genes in total), in the model, generating 1325 genes in total. The eQTL and GRN linkages from PsychENCODE are then restricted to this subset of genes.

### LNCTP Simulating Perturbations

To perform perturbations in this model corresponding to the 29 CRISPR targeted genes, we use the following perturbation-conditioned version of the LNCTP model:

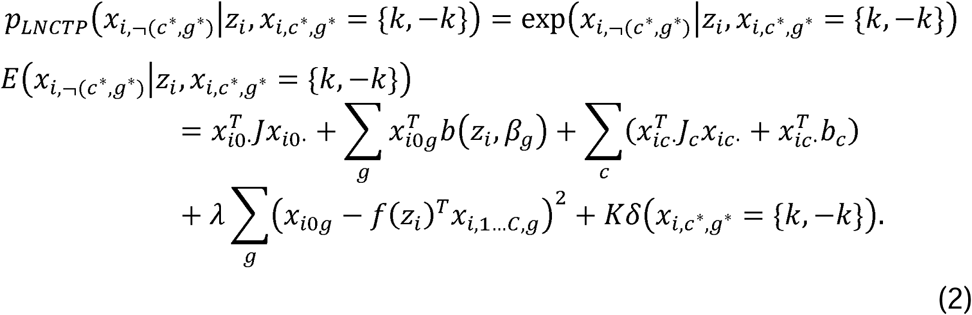

where (*c*\*,*g*\*) denotes the perturbed gene and cell type, whose expression is set to *k* or -*k*, *δ*(*a*) turn in the bulk network, using k=2, and applying a negative perturbation to mimic the effect of the CRISPR perturbation. We note that, since the model is trained on Z-scored log-normalized expression counts, this corresponds to introducing a large negative fold-change to the selected gene. The *in silico* predicted log fold-changes per individual across all genes (per cell-type) are then calculated by comparing the expected values before and after perturbation:

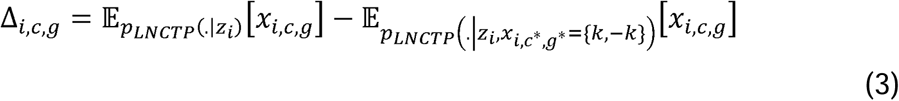

and the final predicted log fold-changes are calculated by taking the expectation across individuals. We use the sampling approach in^76^ to evaluate the expectations in Eq. (3).

To perform perturbations across all 102 NDD genes, for efficiency we learn a reduced model by remove the dependency on *z_i_* in Eq. (1). We sample cell-type specific expression values for each individual from the full model, and then fit the reduced model by refitting the model parameters to maximize the likelihood of the full data vectors (consisting of the original bulk and sampled cell-specific expression vectors for each individual). Perturbations are performed in the reduced model as in Eq. (2) and fold-changes are calculated as in Eq. (3), while removing the dependency on *z_i_* and the *i* subscripts respectively.

### LNCTP in silico convergent genes

To identify *in silico* convergent genes for a set of perturbations, *S* = 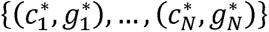, we calculate Δ*_c,g_* using Eq. (3) for each perturbation, writing 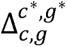 for the log fold-change to (*c*,*g*) generated by applying perturbation (*c*\*, *g*\*), and 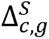 for the set of log fold-changes by applying all perturbations in *S*. Then, the set of *in silico* convergent genes for *S* is found by selecting those for which 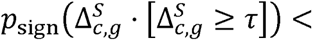, where *p_sign_*(.) is the p-value from a 2-tailed one-sample sign-test. The threshold *τ* is introduced to reduce noise from perturbations which are estimated to generate small log fold-changes, and throughout we set *τ =* 0.3.

For the comparison of *in silico* convergent genes derived from different perturbation sets *S*, we apply two-sided hypergeometric tests to the gene sets defined as above (using all 1325 genes in our model as the background set). For Gene Set Enrichment Analysis of convergent genes derived from *S*, we apply clusterprofiler^178^ to the full set of genes in our model, ranked by 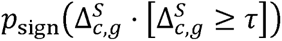 as defined above.

### LNCTP semantic distance test

To test the semantic distance between enriched terms for two sets of perturbations *S*_1_ and *S*_2_, we generate the set of enriched terms *T*_1_, *T*_2_ by applying GSEA to each set as described above (using Benjamini Höchberg correction and an FDR threshold of 0.2 to select enriched terms *T*_1_ and *T*_2_). We then calculate the similarity between terms *t*_1_ and *t*_2_ by evaluating *s*(*t*_1_,*t*_2_) = |*G*(*t*_1_) ∩ *G*(*t*_2_) |/| *G*(*t*_1_) ∪ *G*(*t*_2_)|, where *G*(*t*) denotes the set of genes occurring in the leading edge of term *t*. We test for a significant semantic distance between *S*_1_ and *S*_2_ by evaluating *s*(*t*_1_,*t*_2_) between all pairs *t*_1_ ∈ *S*_1_, *t*_2_ ∈ *S*_2_, versus all pairs t1 ∈ *S*_1_, *t*_2_ ∈ *S*_2_ and *t*_1_ ∈ *S*_2_, *t*_2_ ∈ *S*_2_, and applying a one-sided rank-sum test for the for a smaller similarity in the former pairs versus the latter.

### Transcriptional correlations between hiPSC-derived neural cells, fetal and adult brain cell types, and the zebrafish brain

We compared wild-type (WT) zebrafish brain expression to gene expression in our hiPSC-derived models and to sign-cell expression data for the fetal and adult PFC (PsychENCODE^181,182^: http://resource.psychencode.org/Datasets/Derived/SC_Decomp/DER-20_Single_cell_expression_processed_TPM.tsv). We first filtered zebrafish gene names and converted them to the appropriate *Homo sapiens* orthologs using the R package *orthogene* (v3.2.1^183^); genes without matched orthologs were dropped from both species. Pseudo-bulk expression data from scramble control cells were used as the baseline expression across NPCs, D7 iGLUTs, D21 iGLUTs, and D36 iGABAs. Pearson’s correlation coefficients between *in vitro* cells, fetal and adult postmortem brain cells, and zebrafish brain were calculated and a Bonferroni correction applied.

### Zebrafish

All procedures involving zebrafish were conducted in accordance with Institutional Animal Care and Use Committee (IACUC; Protocol #2024-20054) regulatory standards at Yale University. Zebrafish larvae were raised at 28°C on a 14:10 hour light:dark cycle. Larvae were grown in 150 mm Petri dishes in blue water (0.3g/L Instant Ocean, 1 mg/L methylene blue, pH 7.0) at a density of 60-80 larvae per dish. Behavioral assays were conducted in zebrafish larvae at 5-7 dpf. At these developmental stages, sex is not yet determined.

### Zebrafish mutant generation

We performed automated, high-throughput, quantitative behavioral profiling of larval zebrafish to measure arousal and sensorimotor processing as a readout of circuit-level deficits resulting from gene perturbation.^60^ We quantified 24 parameters across sleep-wake activity and visual-startle responses in 18 stable homozygous mutant or F0 mosaic crispant lines for 15 NDD genes (**SI Tables 4-5**). Stable zebrafish lines were generated by our lab (*arid1b^Δ7/Δ7^*, *chd2^Δ7/Δ7^, chd8^Δ7/Δ7^, chd8^Δ5/Δ5^, kdm5ba^Δ17/Δ17^b^Δ14/Δ14^*, *kdm5ba^Δ4i/Δ4i^b^Δ4/Δ4^*)^60^ or provided as a generous gift from the Thyme lab (*ash1l^1i,Δ60,19i/1i,Δ60,19i^, kmt5b^ΔD208,1i,ΔD5/ΔD208,1i,ΔD5^, kmt2ca^Δ82,17i/Δ82,17i^b^Δ6,Δ29/Δ6,Δ29^, nrxn1a^Δ218/Δ218^*)^184,185^. F0 crispants for the following genes were generated according to ref. ^186^: *chd2, kdm6bab, mbd5, phf12ab, phf21aab, skiab, smarcc2, wacab*. Briefly, we designed two CRISPR crRNAs per allele, prioritizing early exons for targeting. CRISPR RNPs were assembled individually and then combined prior to injection at the one-cell stage. The number of scrambled guides injected into the control group was matched to the number of CRISPR guides used for the experimental group. Injected embryos were raised to 5 dpf at which point the behavioral assays (described below) were conducted. We identified unique behavioral fingerprints for each NDD gene mutant, revealing convergent and divergent phenotypes across mutants (**SI Fig. 22B**). To classify convergent behavioral subgroups that may share circuit-level functions, we performed correlation analyses with hierarchical clustering across mutants. We identified four distinct subgroups of NDD genes with highly correlated behavioral features (**Fig. 7A**).

### Behavioral assays

Larvae were placed into individual wells of a 96 well plate (7701-1651; Whatman, Clifton, NJ) containing 650 μL of standard embryo water (0.3 g/L Instant Ocean, 1 mg/L methylene blue, pH 7.0) per well within a Zebrabox (Viewpoint LifeSciences; Viewpoint Life Sciences, Montreal, Quebec, Canada). Locomotion was quantified with automated video-tracking system (Zebrabox and ZebraLab software). The visual-startle assay was conducted at 5 days post fertilization (dpf) as described^60^. To assess larval responses to lights-off stimuli (VSR-OFF), larvae were acclimated to white light for 1 hour, and baseline activity was tracked for 30 minutes followed by five 1-second dark flashes with intermittent white light for 29 seconds. To evaluate larval responsivity to lights-on stimuli (VSR-ON), the assay was reversed, where larvae were acclimated to darkness for 1 hour, and baseline activity was tracked for 30 minutes followed by five 1-second white light flashes with intermittent darkness for 29 seconds. For VSR-OFF and VSR-ON, six behavioral parameters were quantified using custom MATLAB code^60^ (available on github at https://github.com/ehoffmanlab/Weinschutz-Mendes-et-al-2023-behavior; DOI:10.5281/zenodo.7644898): (i) average intensity of all startle responses; (ii) average post-stimulus activity; (iii) average activity after first stimulus; (iv) stimulus versus post-stimulus activity; (v) intensity of responses to the first stimulus; (vi) intensity of responses to the final stimulus. The sleep-wake paradigm was conducted between 5-7 dpf, following the VSR-OFF and VSR-ON assays. During a 14h:10h white light:darkness cycle, larvae activity and sleep patterns were tracked within the Zebrabox and analyzed with custom MATLAB code^60^ (available on github at (https://github.com/JRihel/Sleep-Analysis/tree/Sleep-Analysis-Code; DOI: 10.5281/zenodo.7644073). Six behavioral parameters were quantified for daytime and nighttime: (i) total activity; (ii) total sleep; (iii) waking activity; (iv) rest bouts; (v) sleep length; (vi) sleep latency. Across VSR-OFF, VSR-ON, and sleep-wake assays, we analyzed 24 parameters.

### Behavioral analysis

Linear mixed models (LMM) were used to compare phenotypes of each behavioral parameter between homozygous mutant versus wild-type or crispant versus scramble-injected fish for each gene of interest. Variations of behavioral phenotypes across experiments were accounted for by including the date of the experiment as a random effect in LMM. Hierarchical clustering analysis was performed to cluster mutants and behavioral parameters based on signed -log10-transformed p-values from LMM, where sign indicates direction of the difference in behavioral phenotype when comparing stable mutant to wild-type or crispant to scrambled-injected. Pearson correlation analysis was used to assess correlations between mutants based on the difference in the 24 parameters. Difference was evaluated using signed -log10-transformed p-values.

### Drug prioritization based on zebrafish pharmaco-behavioral profiles

NDD gene-associated mutant and crispant behavioral phenotypes were compared to a dataset of 376 U.S. FDA-approved drugs that were screened for their behavioral effects in larval zebrafish using the visual-startle and sleep-wake assays described above. These drugs have a significant effect on at least two behavioral parameters (LMM, p<0.05/3, corrected for three behavioral assays). Pearson’s correlation analysis was used to identify drugs that significantly correlate (correlation >0.5, p<0.05, t-statistic) or anti-correlate (correlation <-0.5, p<0.05, t-statistic) with mutant behavioral signatures (**SI Data 2-3**).

### Drug prioritization based on perturbation signature reversal in LiNCs Neuronal Cell Lines

To identify drugs that could reverse cell-type specific convergence across different KOs, we used the Query tool from The Broad Institute’s Connectivity Map (Cmap) Server^78^. Briefly, the tool computes weighted enrichment scores (WTCS) between the query set and each signature in the Cmap LiNCs gene expression data (dose, time, drug, cell-line), normalizes the WTCS by dividing by the signed mean within each perturbation (NCS), and computes FDR as fraction of “null signatures” (DMSO) where the absolute NCS exceeds reference signature. We prioritized drugs that were negatively enriched for convergent signatures specifically in neuronal cells (either neurons (NEU) or neural progenitor cells (NPCs) with NCS <= −1.00, FDR<=0.05) and filtered for drugs that had clinical data in humans and paired behavioral phenotyping in zebrafish (**SI Data 2**).

### Targeted drug rescue of behavioral phenotypes in zebrafish

For mutant-x-drug experiments, larval activity was monitored from 5-7 dpf using the behavioral assays described above. Individual wild-type zebrafish larvae were added to each well of a 96-well plate containing 650 μl of standard embryo water. A 5 mM stock solution of each compound dissolved in DMSO or DMSO alone (control) was pipetted directly into each well after which the visual-startle and sleep-wake assays were performed. Drugs were tested at a final concentration of 10 μM (0.1% DMSO final concentration) in 12-24 background-matched homozygous or wild-type larvae or 24 crispant or scrambled control-injected larvae with genotyping conducted after each experiment to confirm genotypes for stable mutant lines and confirm on-target mutations in crispants.

For behaviors that were nominally significantly different between mutant+DMSO and WT+DMSO (p<0.06), we characterized the effect of the mutant-x-Drug on behavior as: i) “exacerbated” [significant effect mutant+Drug-v-WT > significant effect mutant-v-WT] if mutant behavior p<=0.06 and mutant-x-drug behavior p.value <= mutant behavior p.value with increased absolute beta values (i.e., stronger p-value with appreciable difference in the magnitude of effect but not direction); ii) “unchanged” [significant effect mutant+drug-v-WT = significant effect mutant-v-WT]; iii) “partial rescue” [significant effect mutant+Drug-v-WT < significant effect mutant-v-WT], if mutant behavior p<=0.06 and mutant-x-drug behavior p>0.06 or if mutant behavior p.value <= mutant-x-drug behavior p.value with reduced effects on the absolute beta value; iv) “rescued” [sig. effect mutant-v-WT, no sig. effect mutant+Drug-v-WT], mutant behavior p<=0.06 and mutant-x-drug behavior p>0.06; v) “over-corrected” [mutant+Drug-v-WT opposite direction of sig. effect mutant-v-WT]. mutant behavior p<=0.06 and mutant-x-drug behavior p<=0.06, with opposing directions of effect. Note “drug specific/side-effects” indicate significant mutant-by-drug effects.

## STATEMENT OF ETHICS

Yale University Institutional Review Board waived ethical approval for this work. Ethical approval was not required because the hiPSC lines, lacking association with any identifying information and widely accessible from a public repository, are thus not considered to be human subject research. Post-mortem brain data are similarly lacking identifiable information and are not considered human subject research.

All procedures involving zebrafish were conducted in accordance with Institutional Animal Care and Use Committee (IACUC; Protocol #2024-20054) regulatory standards at Yale University.

## CONFLICT OF INTEREST STATEMENT

The authors declare no conflict of interest.

## FUNDING SOURCES

This work was supported by F31MH130122 (K.R.T), HHMI Gilliams Fellowship (A.P.), Autism Science Foundation (A.P.), T32MH014276 (M.F.G.), T32GM136651 (E.D., S.F.) R01MH123155 (K.J.B.), RM1MH132648 (K.J.B. and E.J.H.), R01MH121074 (K.J.B.), R01MH116002 (E.J.H.) R21MH133245 (E.J.H.), and R01ES033630 (L.H., K.J.B.), R01MH124839 (LMH), R01MH118278 (L.MH.), Simons Foundation (#1012863KB, #573508EH and #345993EH), Spector Fund, (E.J.H. and Swebilius Foundation (E.J.H.); Kavli Foundation (E.J.H.); BD2: Breakthrough Discoveries for thriving with Bipolar Disorder (#DG230102 H.S., M.D., T.C.H., K.J.B.), the European Union’s Horizon 2020 research and innovation programme under the Marie Skłodowska-Curie grant (#101065629 N.B.); and Interdepartmental Neuroscience Program at Yale (A.P).

## AUTHOR CONTRIBUTIONS

MFG designed and executed the NDD gene ECCITEseq in iNPCs, iGLUTs, and iGABAs, with technical assistance from SC, OL, and JC and support from PJMD. KRT conducted all bioinformatic analyses and generated all figures, with technical assistance from AS. Arrayed KO lines and neuronal cultures were generated, validated, phenotyped, and analyzed by NB. NB, T-CH, CB, ALTS, and JL conducted mitochondrial phenotypic analyses. SBT, AP, ED, YD, SEF, and SK, generated, phenotyped, and analyzed zebrafish mutants with technical assistance from GD and bioinformatic supervision from ZW. JW, RM, ZC, and MG conducted LNCTP analyses. Funding and mentorship provided by LH, EJH, and KJB. Manuscript was written by KJB with extensive feedback from LH, EJH, and KRT as well as contributions for all authors.

Special thanks to Michael Talkowski and Douglas Ruderfer for countless discussions on convergence and to Summer Thyme and the Thyme lab for sharing zebrafish mutant lines.

## INCLUSION AND DIVERSITY

One or more of the authors of this paper self-identifies as an under-represented ethnic minority in their field of research or within their geographical location. One or more of the authors of this paper self-identifies as living with a disability. One or more of the authors of this paper self-identifies as a gender minority in their field of research. One or more of the authors of this paper self-identifies as a member of the LGBTQIA+ community. One or more of the authors of this paper received support from a program designed to increase minority representation in their field of research.

## DATA AVAILABILITY

All source donor hiPSCs have been deposited at the Rutgers University Cell and DNA Repository (study 160; http://www.nimhstemcells.org/).

sc-RNA sequencing data reported in this paper will be uploaded to Gene expression omnibus (GEO) prior to publication. Previously published SCZ-CRISPRa screen datasets that were used for external validation of random forest models are available on the GEO (GSE200774) and on Synapse (syn27819129).

## CODE AVAILABILITY

The full analysis pipeline (including code and processed data objects) used for analysis of single-cell CRISPR-KO data, evaluation and characterization of gene-level and network level convergence, and predictive modeling using random forest will be publicly available through Synpase prior to publication.

Custom MATLAB software developed by the Hoffman Lab to analyze visual-startle response parameters is available on github at https://github.com/ehoffmanlab/Weinschutz-Mendes-et-al-2023-behavior; https://doi.org/10.5281/zenodo.7644898. Custom MATLAB sofyware developed by Jason Rihel to analyze sleep-wake assays is available on github at https://github.com/JRihel/Sleep-Analysis/tree/Sleep-Analysis-Code; https://doi.org/10.5281/zenodo.7644073.

## Notes

### Competing Interest Statement

The authors have declared no competing interest.

### Summary of Updates

Figure 6 was an old version that did not match figure legend.

## REFERENCES

1 Sandin, S. et al. The Heritability of Autism Spectrum Disorder. JAMA 318, 1182–1184 (2017). 10.1001/jama.2017.12141

2 Grove, J. et al. Identification of common genetic risk variants for autism spectrum disorder. Nat Genet 51, 431–444 (2019). 10.1038/s41588-019-0344-8

3 Mahjani, B. et al. Prevalence and phenotypic impact of rare potentially damaging variants in autism spectrum disorder. Mol Autism 12, 65 (2021). 10.1186/s13229-021-00465-3

4 Satterstrom, F. K. et al. Large-Scale Exome Sequencing Study Implicates Both Developmental and Functional Changes in the Neurobiology of Autism. Cell 180, 568–584 e523 (2020). 10.1016/j.cell.2019.12.036

5 Kaplanis, J. et al. Evidence for 28 genetic disorders discovered by combining healthcare and research data. Nature 586, 757–762 (2020). 10.1038/s41586-020-2832-5

6 Coe, B. P. et al. Neurodevelopmental disease genes implicated by de novo mutation and copy number variation morbidity. Nat Genet 51, 106–116 (2019). 10.1038/s41588-018-0288-4

7 Singh, T. et al. Rare coding variants in ten genes confer substantial risk for schizophrenia. Nature 604, 509–516 (2022). 10.1038/s41586-022-04556-w

8 Palmer, D. S. et al. Exome sequencing in bipolar disorder identifies AKAP11 as a risk gene shared with schizophrenia. Nat Genet 54, 541–547 (2022). 10.1038/s41588-022-01034-x

9 Willsey, A. J. et al. Coexpression networks implicate human midfetal deep cortical projection neurons in the pathogenesis of autism. Cell 155, 997–1007 (2013). 10.1016/j.cell.2013.10.020

10 De Rubeis, S. et al. Synaptic, transcriptional and chromatin genes disrupted in autism. Nature 515, 209–215 (2014). 10.1038/nature13772

11 O’Roak, B. J. et al. Recurrent de novo mutations implicate novel genes underlying simplex autism risk. Nat Commun 5, 5595 (2014). 10.1038/ncomms6595

12 Talkowski, M. E. et al. Sequencing chromosomal abnormalities reveals neurodevelopmental loci that confer risk across diagnostic boundaries. Cell 149, 525–537 (2012). 10.1016/j.cell.2012.03.028

13 Neale, B. M. et al. Patterns and rates of exonic de novo mutations in autism spectrum disorders. Nature 485, 242–245 (2012). nature11011 [pii] 10.1038/nature11011

14 O’Roak, B. J. et al. Multiplex targeted sequencing identifies recurrently mutated genes in autism spectrum disorders. Science 338, 1619–1622 (2012). 10.1126/science.1227764

15 Sanders, S. J. et al. De novo mutations revealed by whole-exome sequencing are strongly associated with autism. Nature 485, 237–241 (2012). nature10945 [pii] 10.1038/nature10945

16 Fazel Darbandi, S., et al. Five autism-associated transcriptional regulators target shared loci proximal to brain-expressed genes. Cell reports 43, 114329 (2024). 10.1016/j.celrep.2024.114329

17 Markenscoff-Papadimitriou, E. et al. Autism risk gene POGZ promotes chromatin accessibility and expression of clustered synaptic genes. Cell reports 37, 110089 (2021). 10.1016/j.celrep.2021.110089

18 Notwell, J. H. et al. TBR1 regulates autism risk genes in the developing neocortex. Genome Res 26, 1013–1022 (2016). 10.1101/gr.203612.115

19 Sugathan, A. et al. CHD8 regulates neurodevelopmental pathways associated with autism spectrum disorder in neural progenitors. Proc Natl Acad Sci U S A 111, E4468–4477 (2014). 10.1073/pnas.1405266111

20 Cotney, J. et al. The autism-associated chromatin modifier CHD8 regulates other autism risk genes during human neurodevelopment. Nat Commun 6, 6404 (2015). 10.1038/ncomms7404

21 O’Neill, A. C. et al. Spatial centrosome proteome of human neural cells uncovers disease-relevant heterogeneity. Science 376, eabf9088 (2022). 10.1126/science.abf9088

22 Sun, N. et al. Autism genes converge on microtubule biology and RNA-binding proteins during excitatory neurogenesis. bioRxiv, 2023.2012.2022.573108 (2024). 10.1101/2023.12.22.573108

23 Kostyanovskaya, E. et al. Convergence of autism proteins at the cilium. bioRxiv (2025). 10.1101/2024.12.05.626924

24 Teerikorpi, N. et al. Ciliary biology intersects autism and congenital heart disease. bioRxiv (2024). 10.1101/2024.07.30.602578

25 Li, C. et al. Single-cell brain organoid screening identifies developmental defects in autism. bioRxiv, 2022.2009.2015.508118 (2022). 10.1101/2022.09.15.508118

26 Martins-Costa, C. et al. ARID1B controls transcriptional programs of axon projection in an organoid model of the human corpus callosum. Cell stem cell 31, 866–885 e814 (2024). 10.1016/j.stem.2024.04.014

27 Paulsen, B. et al. Autism genes converge on asynchronous development of shared neuron classes. Nature 602, 268–273 (2022). 10.1038/s41586-021-04358-6

28 Villa, C. E. et al. CHD8 haploinsufficiency links autism to transient alterations in excitatory and inhibitory trajectories. Cell reports 39, 110615 (2022). 10.1016/j.celrep.2022.110615

29 Flaherty, E. et al. Neuronal impact of patient-specific aberrant NRXN1alpha splicing. Nat Genet 51, 1679–1690 (2019). 10.1038/s41588-019-0539-z

30 Sebastian, R. et al. Schizophrenia-associated NRXN1 deletions induce developmental-timing- and cell-type-specific vulnerabilities in human brain organoids. Nature Communications 14, 3770 (2023). 10.1038/s41467-023-39420-6

31 Birtele, M. et al. Non-synaptic function of the autism spectrum disorder-associated gene SYNGAP1 in cortical neurogenesis. Nat Neurosci 26, 2090–2103 (2023). 10.1038/s41593-023-01477-3

32 Ellingford, R. A. et al. Cell-type-specific synaptic imbalance and disrupted homeostatic plasticity in cortical circuits of ASD-associated Chd8 haploinsufficient mice. Mol Psychiatry 26, 3614–3624 (2021). 10.1038/s41380-021-01070-9

33 Shi, X. et al. Heterozygous deletion of the autism-associated gene CHD8 impairs synaptic function through widespread changes in gene expression and chromatin compaction. Am J Hum Genet 110, 1750–1768 (2023). 10.1016/j.ajhg.2023.09.004

34 Pak, C. et al. Cross-platform validation of neurotransmitter release impairments in schizophrenia patient-derived NRXN1-mutant neurons. Proc Natl Acad Sci U S A 118 (2021). 10.1073/pnas.2025598118

35 Yi, F. et al. Autism-associated SHANK3 haploinsufficiency causes Ih channelopathy in human neurons. Science 352, aaf2669 (2016). 10.1126/science.aaf2669

36 Vermaercke, B. et al. SYNGAP1 deficiency disrupts synaptic neoteny in xenotransplanted human cortical neurons in vivo. Neuron (2024). 10.1016/j.neuron.2024.07.007

37 Jung, E. M. et al. Arid1b haploinsufficiency disrupts cortical interneuron development and mouse behavior. Nat Neurosci 20, 1694–1707 (2017). 10.1038/s41593-017-0013-0

38 Fernando, M. B. et al. Phenotypic complexities of rare heterozygous neurexin-1 deletions. bioRxiv (2024). 10.1101/2023.10.28.564543

39 Chen, Q. et al. Dysfunction of cortical GABAergic neurons leads to sensory hyper-reactivity in a Shank3 mouse model of ASD. Nat Neurosci 23, 520–532 (2020). 10.1038/s41593-020-0598-6

40 Geschwind, D. H. Autism: many genes, common pathways? Cell 135, 391–395 (2008). 10.1016/j.cell.2008.10.016

41 Willsey, H. R., Willsey, A. J., Wang, B. & State, M. W. Genomics, convergent neuroscience and progress in understanding autism spectrum disorder. Nat Rev Neurosci 23, 323–341 (2022). 10.1038/s41583-022-00576-7

42 Bicks, L. K. & Geschwind, D. H. Functional neurogenomics in autism spectrum disorders: A decade of progress. Curr Opin Neurobiol 86, 102858 (2024). 10.1016/j.conb.2024.102858

43 Quesnel-Vallieres, M., Weatheritt, R. J., Cordes, S. P. & Blencowe, B. J. Autism spectrum disorder: insights into convergent mechanisms from transcriptomics. Nat Rev Genet 20, 51–63 (2019). 10.1038/s41576-018-0066-2

44 Parikshak, N. N. et al. Integrative functional genomic analyses implicate specific molecular pathways and circuits in autism. Cell 155, 1008–1021 (2013). 10.1016/j.cell.2013.10.031

45 Voineagu, I. et al. Transcriptomic analysis of autistic brain reveals convergent molecular pathology. Nature 474, 380–384 (2011). nature10110 [pii] 10.1038/nature10110

46 Liao, C. et al. Convergent coexpression of autism-associated genes suggests some novel risk genes may not be detectable in large-scale genetic studies. Cell Genom 3, 100277 (2023). 10.1016/j.xgen.2023.100277

47 Pintacuda, G. et al. Protein interaction studies in human induced neurons indicate convergent biology underlying autism spectrum disorders. Cell Genom 3, 100250 (2023). 10.1016/j.xgen.2022.100250

48 Wang, B. et al. A foundational atlas of autism protein interactions reveals molecular convergence. bioRxiv (2023). 10.1101/2023.12.03.569805

49 Murtaza, N. et al. Neuron-specific protein network mapping of autism risk genes identifies shared biological mechanisms and disease-relevant pathologies. Cell reports 41, 111678 (2022). 10.1016/j.celrep.2022.111678

50 Gao, Y. et al. Proximity analysis of native proteomes reveals phenotypic modifiers in a mouse model of autism and related neurodevelopmental conditions. Nat Commun 15, 6801 (2024). 10.1038/s41467-024-51037-x

51 Rubenstein, J. L. & Merzenich, M. M. Model of autism: increased ratio of excitation/inhibition in key neural systems. Genes, brain, and behavior 2, 255–267 (2003). 10.1034/j.1601-183x.2003.00037.x

52 Antoine, M. W., Langberg, T., Schnepel, P. & Feldman, D. E. Increased Excitation-Inhibition Ratio Stabilizes Synapse and Circuit Excitability in Four Autism Mouse Models. Neuron 101, 648–661 e644 (2019). 10.1016/j.neuron.2018.12.026

53 Nelson, S. B. & Valakh, V. Excitatory/Inhibitory Balance and Circuit Homeostasis in Autism Spectrum Disorders. Neuron 87, 684–698 (2015). 10.1016/j.neuron.2015.07.033

54 Cederquist, G. Y. et al. A Multiplex Human Pluripotent Stem Cell Platform Defines Molecular and Functional Subclasses of Autism-Related Genes. Cell stem cell 27, 35–49 e36 (2020). 10.1016/j.stem.2020.06.004

55 Lalli, M. A., Avey, D., Dougherty, J. D., Milbrandt, J. & Mitra, R. D. High-throughput single-cell functional elucidation of neurodevelopmental disease-associated genes reveals convergent mechanisms altering neuronal differentiation. Genome Res 30, 1317–1331 (2020). 10.1101/gr.262295.120

56 Meng, X. et al. Assembloid CRISPR screens reveal impact of disease genes in human neurodevelopment. Nature 622, 359–366 (2023). 10.1038/s41586-023-06564-w

57 Li, C. et al. Single-cell brain organoid screening identifies developmental defects in autism. Nature 621, 373–380 (2023). 10.1038/s41586-023-06473-y

58 Jin, X. et al. In vivo Perturb-Seq reveals neuronal and glial abnormalities associated with autism risk genes. Science 370 (2020). 10.1126/science.aaz6063

59 Willsey, H. R. et al. Parallel in vivo analysis of large-effect autism genes implicates cortical neurogenesis and estrogen in risk and resilience. Neuron 109, 1409 (2021). 10.1016/j.neuron.2021.03.030

60 Weinschutz Mendes, H., et al. High-throughput functional analysis of autism genes in zebrafish identifies convergence in dopaminergic and neuroimmune pathways. Cell reports 42, 112243 (2023). 10.1016/j.celrep.2023.112243

61 Fu, J. M. et al. Rare coding variation provides insight into the genetic architecture and phenotypic context of autism. Nat Genet 54, 1320–1331 (2022). 10.1038/s41588-022-01104-0

62 Marshall, C. R. et al. Contribution of copy number variants to schizophrenia from a genome-wide study of 41,321 subjects. Nat Genet 49, 27–35 (2017). 10.1038/ng.3725

63 Johannesen, K. M. et al. Defining the phenotypic spectrum of SLC6A1 mutations. Epilepsia 59, 389–402 (2018). 10.1111/epi.13986

64 Heyne, H. O. et al. De novo variants in neurodevelopmental disorders with epilepsy. Nat Genet 50, 1048–1053 (2018). 10.1038/s41588-018-0143-7

65 de Leeuw, C. A., Mooij, J. M., Heskes, T. & Posthuma, D. MAGMA: generalized gene-set analysis of GWAS data. PLoS Comput Biol 11, e1004219 (2015). 10.1371/journal.pcbi.1004219

66 Sullivan, P. F. & Geschwind, D. H. Defining the Genetic, Genomic, Cellular, and Diagnostic Architectures of Psychiatric Disorders. Cell 177, 162–183 (2019). 10.1016/j.cell.2019.01.015

67 Schrode, N. et al. Synergistic effects of common schizophrenia risk variants. Nat Genet 51, 1475–1485 (2019). 10.1038/s41588-019-0497-5

68 Wells, M. F. et al. Natural variation in gene expression and viral susceptibility revealed by neural progenitor cell villages. Cell stem cell 30, 312–332 e313 (2023). 10.1016/j.stem.2023.01.010

69 Zhang, Y. et al. Rapid single-step induction of functional neurons from human pluripotent stem cells. Neuron 78, 785–798 (2013). 10.1016/j.neuron.2013.05.029

70 Yang, N. et al. Generation of pure GABAergic neurons by transcription factor programming. Nat Methods (2017). 10.1038/nmeth.4291

71 Mimitou, E. P. et al. Multiplexed detection of proteins, transcriptomes, clonotypes and CRISPR perturbations in single cells. Nat Methods 16, 409–412 (2019). 10.1038/s41592-019-0392-0

72 Hao, Y. et al. Integrated analysis of multimodal single-cell data. Cell 184, 3573–3587 e3529 (2021). 10.1016/j.cell.2021.04.048

73 Deans, P. J. M. et al. Convergent impact of schizophrenia risk genes. bioRxiv, 2022.2003.2029.486286 (2025). 10.1101/2022.03.29.486286

74 Deans, P. M. et al. Non-additive effects of schizophrenia risk genes reflect convergent downstream function. medRxiv, 2023.2003.2020.23287497 (2023). 10.1101/2023.03.20.23287497

75 Breiman, L. Random Forests. Machine Learning 45, 5–32 (2001). 10.1023/A:1010933404324

76 Emani, P. S. et al. Single-cell genomics and regulatory networks for 388 human brains. Science 384, eadi5199 (2024). 10.1126/science.adi5199

77 Gonzalez, F. et al. An iCRISPR platform for rapid, multiplexable, and inducible genome editing in human pluripotent stem cells. Cell stem cell 15, 215–226 (2014). 10.1016/j.stem.2014.05.018

78 Subramanian, A. et al. A Next Generation Connectivity Map: L1000 Platform and the First 1,000,000 Profiles. Cell 171, 1437–1452 e1417 (2017). 10.1016/j.cell.2017.10.049

79 Manji, H. et al. Impaired mitochondrial function in psychiatric disorders. Nat Rev Neurosci 13, 293–307 (2012). 10.1038/nrn3229

80 Rossignol, D. A. & Frye, R. E. Mitochondrial dysfunction in autism spectrum disorders: a systematic review and meta-analysis. Mol Psychiatry 17, 290–314 (2012). 10.1038/mp.2010.136

81 Wang, Y., Picard, M. & Gu, Z. Genetic Evidence for Elevated Pathogenicity of Mitochondrial DNA Heteroplasmy in Autism Spectrum Disorder. PLoS genetics 12, e1006391 (2016). 10.1371/journal.pgen.1006391

82 Varga, N. A. et al. Mitochondrial dysfunction and autism: comprehensive genetic analyses of children with autism and mtDNA deletion. Behav Brain Funct 14, 4 (2018). 10.1186/s12993-018-0135-x

83 Chalkia, D. et al. Association Between Mitochondrial DNA Haplogroup Variation and Autism Spectrum Disorders. JAMA Psychiatry 74, 1161–1168 (2017). 10.1001/jamapsychiatry.2017.2604

84 Wang, Y. et al. Association of mitochondrial DNA content, heteroplasmies and inter-generational transmission with autism. Nat Commun 13, 3790 (2022). 10.1038/s41467-022-30805-7

85 Yardeni, T. et al. An mtDNA mutant mouse demonstrates that mitochondrial deficiency can result in autism endophenotypes. Proc Natl Acad Sci U S A 118 (2021). 10.1073/pnas.2021429118

86 Shen, M. et al. Reduced mitochondrial fusion and Huntingtin levels contribute to impaired dendritic maturation and behavioral deficits in Fmr1-mutant mice. Nat Neurosci 22, 386–400 (2019). 10.1038/s41593-019-0338-y

87 Fernandez, A. et al. Mitochondrial Dysfunction Leads to Cortical Under-Connectivity and Cognitive Impairment. Neuron 102, 1127–1142 e1123 (2019). 10.1016/j.neuron.2019.04.013

88 Gandal, M. J. et al. Shared molecular neuropathology across major psychiatric disorders parallels polygenic overlap. Science 359, 693–697 (2018). 10.1126/science.aad6469

89 Kanellopoulos, A. K. et al. Aralar Sequesters GABA into Hyperactive Mitochondria, Causing Social Behavior Deficits. Cell 180, 1178–1197 e1120 (2020). 10.1016/j.cell.2020.02.044

90 Li, J. et al. Mitochondrial deficits in human iPSC-derived neurons from patients with 22q11.2 deletion syndrome and schizophrenia. Translational psychiatry 9, 302 (2019). 10.1038/s41398-019-0643-y

91 Li, J. et al. Association of Mitochondrial Biogenesis With Variable Penetrance of Schizophrenia. JAMA Psychiatry 78, 911–921 (2021). 10.1001/jamapsychiatry.2021.0762

92 Schafer, S. T. et al. Pathological priming causes developmental gene network heterochronicity in autistic subject-derived neurons. Nat Neurosci 22, 243–255 (2019). 10.1038/s41593-018-0295-x

93 Rajarajan, P. et al. Neuron-specific signatures in the chromosomal connectome associated with schizophrenia risk. Science 362 (2018). 10.1126/science.aat4311

94 Wen, C. et al. Cross-ancestry atlas of gene, isoform, and splicing regulation in the developing human brain. Science 384, eadh0829 (2024). 10.1126/science.adh0829

95 Wong, C. C. Y. et al. Genome-wide DNA methylation profiling identifies convergent molecular signatures associated with idiopathic and syndromic autism in post-mortem human brain tissue. Hum Mol Genet 28, 2201–2211 (2019). 10.1093/hmg/ddz052

96 Ramaswami, G. et al. Integrative genomics identifies a convergent molecular subtype that links epigenomic with transcriptomic differences in autism. Nat Commun 11, 4873 (2020). 10.1038/s41467-020-18526-1

97 Bell, J. Stratified medicines: towards better treatment for disease. Lancet 383 Suppl 1, S3–5 (2014). 10.1016/S0140-6736(14)60115-X

98 Tsimberidou, A. M. et al. Molecular tumour boards - current and future considerations for precision oncology. Nature reviews. Clinical oncology 20, 843–863 (2023). 10.1038/s41571-023-00824-4

99 Zhang, H., Colclough, K., Gloyn, A. L. & Pollin, T. I. Monogenic diabetes: a gateway to precision medicine in diabetes. J Clin Invest 131 (2021). 10.1172/JCI142244

100 Gaugler, T. et al. Most genetic risk for autism resides with common variation. Nat Genet 46, 881–885 (2014). 10.1038/ng.3039

101 Schaaf, C. P. et al. A framework for an evidence-based gene list relevant to autism spectrum disorder. Nat Rev Genet 21, 367–376 (2020). 10.1038/s41576-020-0231-2

102 Shi, Y. et al. Multi-polygenic scores in psychiatry: From disorder specific to transdiagnostic perspectives. Am J Med Genet B Neuropsychiatr Genet 195, e32951 (2024). 10.1002/ajmg.b.32951

103 Andreassen, O. A., Hindley, G. F. L., Frei, O. & Smeland, O. B. New insights from the last decade of research in psychiatric genetics: discoveries, challenges and clinical implications. World Psychiatry 22, 4–24 (2023). 10.1002/wps.21034

104 Weiner, D. J. et al. Statistical and functional convergence of common and rare genetic influences on autism at chromosome 16p. Nat Genet 54, 1630–1639 (2022). 10.1038/s41588-022-01203-y

105 Weiner, D. J. et al. Polygenic transmission disequilibrium confirms that common and rare variation act additively to create risk for autism spectrum disorders. Nat Genet 49, 978–985 (2017). 10.1038/ng.3863

106 Klei, L. et al. How rare and common risk variation jointly affect liability for autism spectrum disorder. Mol Autism 12, 66 (2021). 10.1186/s13229-021-00466-2

107 Bergen, S. E. et al. Joint Contributions of Rare Copy Number Variants and Common SNPs to Risk for Schizophrenia. Am J Psychiatry 176, 29–35 (2019). 10.1176/appi.ajp.2018.17040467

108 Akingbuwa, W. A., Hammerschlag, A. R., Bartels, M., Nivard, M. G. & Middeldorp, C. M. Ultra-rare and common genetic variant analysis converge to implicate negative selection and neuronal processes in the aetiology of schizophrenia. Mol Psychiatry 27, 3699–3707 (2022). 10.1038/s41380-022-01621-8

109 Oliver, K. L. et al. Common risk variants for epilepsy are enriched in families previously targeted for rare monogenic variant discovery. EBioMedicine 81, 104079 (2022). 10.1016/j.ebiom.2022.104079

110 Genetic Modifiers of Huntington’s Disease, C. Identification of Genetic Factors that Modify Clinical Onset of Huntington’s Disease. Cell 162, 516–526 (2015). 10.1016/j.cell.2015.07.003

111 Kingdom, R., Beaumont, R. N., Wood, A. R., Weedon, M. N. & Wright, C. F. Genetic modifiers of rare variants in monogenic developmental disorder loci. Nat Genet 56, 861–868 (2024). 10.1038/s41588-024-01710-0

112 Dobrindt, K. et al. Publicly Available hiPSC Lines with Extreme Polygenic Risk Scores for Modeling Schizophrenia. Complex Psychiatry 6, 68–82 (2021). 10.1159/000512716

113 Bjork, M. H. et al. Association of Prenatal Exposure to Antiseizure Medication With Risk of Autism and Intellectual Disability. JAMA Neurol 79, 672–681 (2022). 10.1001/jamaneurol.2022.1269

114 Anton-Bolanos, N. et al. Brain Chimeroids reveal individual susceptibility to neurotoxic triggers. Nature (2024). 10.1038/s41586-024-07578-8

115 Gandal, M. J. et al. Broad transcriptomic dysregulation occurs across the cerebral cortex in ASD. Nature 611, 532–539 (2022). 10.1038/s41586-022-05377-7

116 Wamsley, B. et al. Molecular cascades and cell type-specific signatures in ASD revealed by single-cell genomics. Science 384, eadh2602 (2024). 10.1126/science.adh2602

117 Yap, C. X. et al. Brain cell-type shifts in Alzheimer’s disease, autism, and schizophrenia interrogated using methylomics and genetics. Sci Adv 10, eadn7655 (2024). 10.1126/sciadv.adn7655

118 Zhang, P. et al. Neuron-specific transcriptomic signatures indicate neuroinflammation and altered neuronal activity in ASD temporal cortex. Proc Natl Acad Sci U S A 120, e2206758120 (2023). 10.1073/pnas.2206758120

119 Bhattacharya, A. et al. Isoform-level transcriptome-wide association uncovers genetic risk mechanisms for neuropsychiatric disorders in the human brain. Nature genetics 55, 2117–2128 (2023). 10.1038/s41588-023-01560-2

120 Gandal, M. J. et al. Transcriptome-wide isoform-level dysregulation in ASD, schizophrenia, and bipolar disorder. Science 362 (2018). 10.1126/science.aat8127

121 Han, V. X., Patel, S., Jones, H. F. & Dale, R. C. Maternal immune activation and neuroinflammation in human neurodevelopmental disorders. Nat Rev Neurol 17, 564–579 (2021). 10.1038/s41582-021-00530-8

122 Seah, C. et al. Modeling gene x environment interactions in PTSD using human neurons reveals diagnosis-specific glucocorticoid-induced gene expression. Nat Neurosci 25, 1434–1445 (2022). 10.1038/s41593-022-01161-y

123 Seah, C. et al. Common genetic variation impacts stress response in the brain. bioRxiv, 2023.2012.2027.573459 (2023). 10.1101/2023.12.27.573459

124 Retallick-Townsley, K. G. et al. Dynamic stress- and inflammatory-based regulation of psychiatric risk loci in human neurons. bioRxiv, 2024.2007.2009.602755 (2024). 10.1101/2024.07.09.602755

125 Cruceanu, C. et al. Cell-Type-Specific Impact of Glucocorticoid Receptor Activation on the Developing Brain: A Cerebral Organoid Study. Am J Psychiatry, appiajp202121010095 (2021). 10.1176/appi.ajp.2021.21010095

126 Teter, O. M. et al. CRISPRi-based screen of Autism Spectrum Disorder risk genes in microglia uncovers roles of ADNP in microglia endocytosis and uptake of synaptic material. bioRxiv, 2024.2006.2001.596962 (2024). 10.1101/2024.06.01.596962

127 Ma, Y. et al. Activity-Dependent Transcriptional Program in NGN2+ Neurons Enriched for Genetic Risk for Brain-Related Disorders. Biol Psychiatry 95, 187–198 (2024). 10.1016/j.biopsych.2023.07.003

128 Roussos, P., Guennewig, B., Kaczorowski, D. C., Barry, G. & Brennand, K. J. Activity-Dependent Changes in Gene Expression in Schizophrenia Human-Induced Pluripotent Stem Cell Neurons. JAMA Psychiatry 73, 1180–1188 (2016). 10.1001/jamapsychiatry.2016.2575

129 Sanchez-Priego, C. et al. Mapping cis-regulatory elements in human neurons links psychiatric disease heritability and activity-regulated transcriptional programs. Cell reports 39, 110877 (2022). 10.1016/j.celrep.2022.110877

130 Boulting, G. L. et al. Activity-dependent regulome of human GABAergic neurons reveals new patterns of gene regulation and neurological disease heritability. Nat Neurosci 24, 437–448 (2021). 10.1038/s41593-020-00786-1

131 Ahn, K. et al. High rate of disease-related copy number variations in childhood onset schizophrenia. Mol Psychiatry 19, 568–572 (2014). 10.1038/mp.2013.59

132 Ahn, K., An, S. S., Shugart, Y. Y. & Rapoport, J. L. Common polygenic variation and risk for childhood-onset schizophrenia. Mol Psychiatry (2014). 10.1038/mp.2014.158

133 Hoffman, G. E. et al. Transcriptional signatures of schizophrenia in hiPSC-derived NPCs and neurons are concordant with post-mortem adult brains. Nat Commun 8, 2225 (2017). 10.1038/s41467-017-02330-5

134 Guss, E. J. et al. Protocol for neurogenin-2-mediated induction of human stem cell-derived neural progenitor cells. Star Protoc 5, 102878 (2024). 10.1016/j.xpro.2024.102878

135 Wang, M. et al. Transformative Network Modeling of Multi-omics Data Reveals Detailed Circuits, Key Regulators, and Potential Therapeutics for Alzheimer’s Disease. Neuron 109, 257–272 e214 (2021). 10.1016/j.neuron.2020.11.002

136 Ho, S. M. et al. Rapid Ngn2-induction of excitatory neurons from hiPSC-derived neural progenitor cells. Methods 101, 113–124 (2016). 10.1016/j.ymeth.2015.11.019

137 Marro, S. G. et al. Neuroligin-4 Regulates Excitatory Synaptic Transmission in Human Neurons. Neuron 103, 617–626 e616 (2019). 10.1016/j.neuron.2019.05.043

138 Zhang, Z. et al. The fragile X mutation impairs homeostatic plasticity in human neurons by blocking synaptic retinoic acid signaling. Science translational medicine 10 (2018). 10.1126/scitranslmed.aar4338

139 Meijer, M. et al. A Single-Cell Model for Synaptic Transmission and Plasticity in Human iPSC-Derived Neurons. Cell reports 27, 2199–2211 e2196 (2019). 10.1016/j.celrep.2019.04.058

140 Zhang, S. et al. Allele-specific open chromatin in human iPSC neurons elucidates functional disease variants. Science 369, 561–565 (2020). 10.1126/science.aay3983

141 Sun, Y. et al. A deleterious Nav1.1 mutation selectively impairs telencephalic inhibitory neurons derived from Dravet Syndrome patients. Elife 5 (2016). 10.7554/eLife.13073

142 Barretto, N. et al. ASCL1- and DLX2-induced GABAergic neurons from hiPSC-derived NPCs. J Neurosci Methods 334, 108548 (2020). 10.1016/j.jneumeth.2019.108548

143 Powell, S. K. et al. Induction of dopaminergic neurons for neuronal subtype-specific modeling of psychiatric disease risk. Mol Psychiatry 28, 1970–1982 (2023). 10.1038/s41380-021-01273-0

144 Miller, J. A. et al. Transcriptional landscape of the prenatal human brain. Nature 508, 199–206 (2014). 10.1038/nature13185

145 Shi, Z. D. et al. Genome Editing in hPSCs Reveals GATA6 Haploinsufficiency and a Genetic Interaction with GATA4 in Human Pancreatic Development. Cell stem cell 20, 675–688 e676 (2017). 10.1016/j.stem.2017.01.001

146 Dobin, A. et al. STAR: ultrafast universal RNA-seq aligner. Bioinformatics 29, 15–21 (2013). 10.1093/bioinformatics/bts635

147 Danecek, P. et al. Twelve years of SAMtools and BCFtools. GigaScience 10 (2021). 10.1093/gigascience/giab008

148 Lindsay, H. et al. CrispRVariants charts the mutation spectrum of genome engineering experiments. Nat Biotechnol 34, 701–702 (2016). 10.1038/nbt.3628

149 Yuan, S. H. et al. Cell-surface marker signatures for the isolation of neural stem cells, glia and neurons derived from human pluripotent stem cells. PLoS One 6, e17540 (2011). 10.1371/journal.pone.0017540

150 Li, W. et al. MAGeCK enables robust identification of essential genes from genome-scale CRISPR/Cas9 knockout screens. Genome biology 15, 554 (2014). 10.1186/s13059-014-0554-4

151 Birey, F. et al. Assembly of functionally integrated human forebrain spheroids. Nature 545, 54–59 (2017). 10.1038/nature22330

152 Butler, A., Hoffman, P., Smibert, P., Papalexi, E. & Satija, R. Integrating single-cell transcriptomic data across different conditions, technologies, and species. Nat Biotechnol 36, 411–420 (2018). 10.1038/nbt.4096

153 Papalexi, E. et al. Characterizing the molecular regulation of inhibitory immune checkpoints with multimodal single-cell screens. Nat Genet 53, 322–331 (2021). 10.1038/s41588-021-00778-2

154 Tirosh, I. et al. Dissecting the multicellular ecosystem of metastatic melanoma by single-cell RNA-seq. Science 352, 189–196 (2016). 10.1126/science.aad0501

155 Tian, R. et al. CRISPR Interference-Based Platform for Multimodal Genetic Screens in Human iPSC-Derived Neurons. Neuron (2019). 10.1016/j.neuron.2019.07.014

156 Willer, C. J., Li, Y. & Abecasis, G. R. METAL: fast and efficient meta-analysis of genomewide association scans. Bioinformatics 26, 2190–2191 (2010). 10.1093/bioinformatics/btq340

157 Saha, A. et al. Co-expression networks reveal the tissue-specific regulation of transcription and splicing. Genome Res 27, 1843–1858 (2017). 10.1101/gr.216721.116

158 Gao, C., McDowell, I. C., Zhao, S., Brown, C. D. & Engelhardt, B. E. Context Specific and Differential Gene Co-expression Networks via Bayesian Biclustering. PLoS Comput Biol 12, e1004791 (2016). 10.1371/journal.pcbi.1004791

159 Gao C, B. C., Engelhardt BE. A latent factor model with a mixture of sparse and dense factors to model gene expression data with confounding effects. arXiv (2013).

160 Yu, G. Gene Ontology Semantic Similarity Analysis Using GOSemSim. Methods in molecular biology 2117, 207–215 (2020). 10.1007/978-1-0716-0301-7_11

161 Demontis, D. et al. Discovery of the first genome-wide significant risk loci for attention deficit/hyperactivity disorder. Nat Genet 51, 63–75 (2019). 10.1038/s41588-018-0269-7

162 Duncan, L. et al. Significant Locus and Metabolic Genetic Correlations Revealed in Genome-Wide Association Study of Anorexia Nervosa. Am J Psychiatry 174, 850–858 (2017). 10.1176/appi.ajp.2017.16121402

163 Walters, R. K. et al. Transancestral GWAS of alcohol dependence reveals common genetic underpinnings with psychiatric disorders. Nat Neurosci 21, 1656–1669 (2018). 10.1038/s41593-018-0275-1

164 Mullins, N. et al. Genome-wide association study of more than 40,000 bipolar disorder cases provides new insights into the underlying biology. Nat Genet 53, 817–829 (2021). 10.1038/s41588-021-00857-4

165 Johnson, E. C. et al. A large-scale genome-wide association study meta-analysis of cannabis use disorder. Lancet Psychiatry 7, 1032–1045 (2020). 10.1016/S2215-0366(20)30339-4

166 Howard, D. M. et al. Genome-wide meta-analysis of depression identifies 102 independent variants and highlights the importance of the prefrontal brain regions. Nat Neurosci 22, 343–352 (2019). 10.1038/s41593-018-0326-7

167 International Obsessive Compulsive Disorder Foundation Genetics, C. & Studies, O. C. D. C. G. A. Revealing the complex genetic architecture of obsessive-compulsive disorder using meta-analysis. Mol Psychiatry 23, 1181–1188 (2018). 10.1038/mp.2017.154

168 Nievergelt, C. M. et al. International meta-analysis of PTSD genome-wide association studies identifies sex- and ancestry-specific genetic risk loci. Nat Commun 10, 4558 (2019). 10.1038/s41467-019-12576-w

169 Trubetskoy, V. et al. Mapping genomic loci implicates genes and synaptic biology in schizophrenia. Nature 604, 502–508 (2022). 10.1038/s41586-022-04434-5

170 Cross-Disorder Group of the Psychiatric Genomics Consortium. Genomic Relationships, Novel Loci, and Pleiotropic Mechanisms across Eight Psychiatric Disorders. Cell 179, 1469–1482 e1411 (2019). 10.1016/j.cell.2019.11.020

171 Marioni, R. E. et al. GWAS on family history of Alzheimer’s disease. Translational psychiatry 8, 99 (2018). 10.1038/s41398-018-0150-6

172 Nalls, M. A. et al. Identification of novel risk loci, causal insights, and heritable risk for Parkinson’s disease: a meta-analysis of genome-wide association studies. The Lancet. Neurology 18, 1091–1102 (2019). 10.1016/S1474-4422(19)30320-5

173 Nicolas, A. et al. Genome-wide Analyses Identify KIF5A as a Novel ALS Gene. Neuron 97, 1268–1283 e1266 (2018). 10.1016/j.neuron.2018.02.027

174 Yu, D. et al. Interrogating the Genetic Determinants of Tourette’s Syndrome and Other Tic Disorders Through Genome-Wide Association Studies. Am J Psychiatry 176, 217–227 (2019). 10.1176/appi.ajp.2018.18070857

175 Hautakangas, H. et al. Genome-wide analysis of 102,084 migraine cases identifies 123 risk loci and subtype-specific risk alleles. Nat Genet 54, 152–160 (2022). 10.1038/s41588-021-00990-0

176 Johnston, K. J. A. et al. Genome-wide association study of multisite chronic pain in UK Biobank. PLoS genetics 15, e1008164 (2019). 10.1371/journal.pgen.1008164

177 Lo, M. T. et al. Genome-wide analyses for personality traits identify six genomic loci and show correlations with psychiatric disorders. Nat Genet 49, 152–156 (2017). 10.1038/ng.3736

178 Yu, G., Wang, L. G., Han, Y. & He, Q. Y. clusterProfiler: an R package for comparing biological themes among gene clusters. OMICS 16, 284–287 (2012). 10.1089/omi.2011.0118

179 Watanabe, K., Taskesen, E., van Bochoven, A. & Posthuma, D. Functional mapping and annotation of genetic associations with FUMA. Nat Commun 8, 1826 (2017). 10.1038/s41467-017-01261-5

180 Wang, J. & Liao, Y. WebGestaltR: Gene Set Analysis Toolkit WebGestaltR. R package version 0.4.3., <https://CRAN.R-project.org/package=WebGestaltR> (2020).

181 Darmanis, S. et al. A survey of human brain transcriptome diversity at the single cell level. Proc Natl Acad Sci U S A 112, 7285–7290 (2015). 10.1073/pnas.1507125112

182 Lake, B. B. et al. Neuronal subtypes and diversity revealed by single-nucleus RNA sequencing of the human brain. Science 352, 1586–1590 (2016). 10.1126/science.aaf1204

183 Schilder, B. M. & Skene, N. G. orthogene: an R package for easy mapping of orthologous genes across hundreds of species. Bioconductor (2022). doi:10.18129/B9.bioc.orthogene.

184 Capps, M. E. S. et al. Disrupted diencephalon development and neuropeptidergic pathways in zebrafish with autism-risk mutations. Proc Natl Acad Sci U S A 122, e2402557122 (2025). 10.1073/pnas.2402557122

185 Calhoun, C. C. S. et al. Removal of developmentally regulated microexons has a minimal impact on larval zebrafish brain morphology and function. bioRxiv, 2024.2008.2019.608697 (2024). 10.1101/2024.08.19.608697

186 Kroll, F. et al. A simple and effective F0 knockout method for rapid screening of behaviour and other complex phenotypes. Elife 10 (2021). 10.7554/eLife.59683

187 Chaudhry, A., Shi, R. & Luciani, D. S. A pipeline for multidimensional confocal analysis of mitochondrial morphology, function, and dynamics in pancreatic β-cells. Am J Physiol Endocrinol Metab 318, E87–E101 (2020).

