## Supplemental Figures & Tables for "Dynamic convergence of neurodevelopmental disorder risk genes across neurodevelopment"

**Inventory of Supplemental Material**

**Supplemental Tables**

**SI Table 1.** gRNA sequences for each NDD target gene.

**SI Table 2.** Cell maturity and subtype markers used to correct for cellular heterogenity and remove subtype clusters in ECCITEseq.

**SI Table 3.** Survival scores for ASD target Genes for Tian et al. 2020 compared with the number of successfully perturbed cells and average abundance of gRNAs by MiSeq.

**SI Table 4.** Generation of stable mutant zebrafish lines.

**SI Table 5.** Design and generation of F0 mutant zebrafish lines.

**Supplemental Figures**

**SI Figure 1**. Characterization of 29 NDD rare variant target genes.
**SI Figure 2**. Characterization of gene expression of 29 NDD rare variant target genes in hiPSC neurons and the developing brain.

**SI Figure 3**. NDD pooled scCRISPR-KO screen validation.

**SI Figure 4**. NDD scCRISPR-KO screen in hiPSC derived iNPCs, iGLUTs, and iGABAs.

**SI Figure 5**. Validation of CRISPR-induced genome editing on selected genes with bulk RNAseq.
**SI Figure 6**. Characterization of perturbed cells and concordance between single-cell and pseudobulk differential gene expression analysis.

**SI Figure 7.** Degree of down-regulation of KO expression does not significantly correlate with downstream transcriptomic impacts.

**SI Figure 8**. Stronger correlations between individual KOs in mature neurons are replicated after random sub-setting of controls.

**SI Figure 9.** GWAS enrichments for individual KO effects across cell-types.

**SI Figure 10.** NDDs show little transcriptomic convergence across cell-types and maturity.
**SI Figure 11**. Gene-level convergence across 9 NDDs is unique across cell-types and increases with maturity in iGLUT neurons.

**SI Figure 12**. Gene-level convergence across 9 NDDs is enriched for pathways involved in neural development, mitochondrial function, and translational regulation.

**SI Figure 13**. Gene-level and network-level convergence across 21 NDDs in iGLUT and iGABA neurons.

**SI Figure. 14.** Schema of random forest training model and external validation.

**SI Figure 15.** Random forest training data characteristics and node frequency across trees.
**SI Figure. 16.** NDD risk genes with strong ASD associations resolve greater convergence across cell-types.

**SI Figure. 17.** Analysis of LNCTP convergent genes.

**SI Figure. 18**. Impact of NDD KO on proliferation and neurogenesis in NPCs and neurons.
**SI Figure. 19.** Pooled CRISPR screen to uncover the NDD KO effects on cortical organoid development.
**SI Figure 20.** High resolution microscopy captures diversity of mitochondrial morphology in iGLUTs and is sensitive to mitochondrial pharmacological insults.

**SI Figure. 21.** NDD KOs converge on mitochondrial function differently in iNPCs and mature iGLUTs.

**SI Figure 22.** Phenotypic clustering of 15 NDD genes in mutant zebrafish.

**SI Figure 23.** Transcriptomic clustering of 15 NDD genes in mature neurons with paired behavioral data in mutant zebrafish.

**SI Figure 24.** Drug Reversal of Behavioral Phenotypes in mutant zebrafish.

**Supplemental Data**

**SI Data 1.** Cell-type-specific convergent gene lists based on NDD pooled scCRISPR-KO.

**SI Data 2.** Convergent sets and drug predictions based on mutant zebrafish behavioral clusters.

**SI Data 3.** Gene x drug mutant zebrafish behavioral effects.

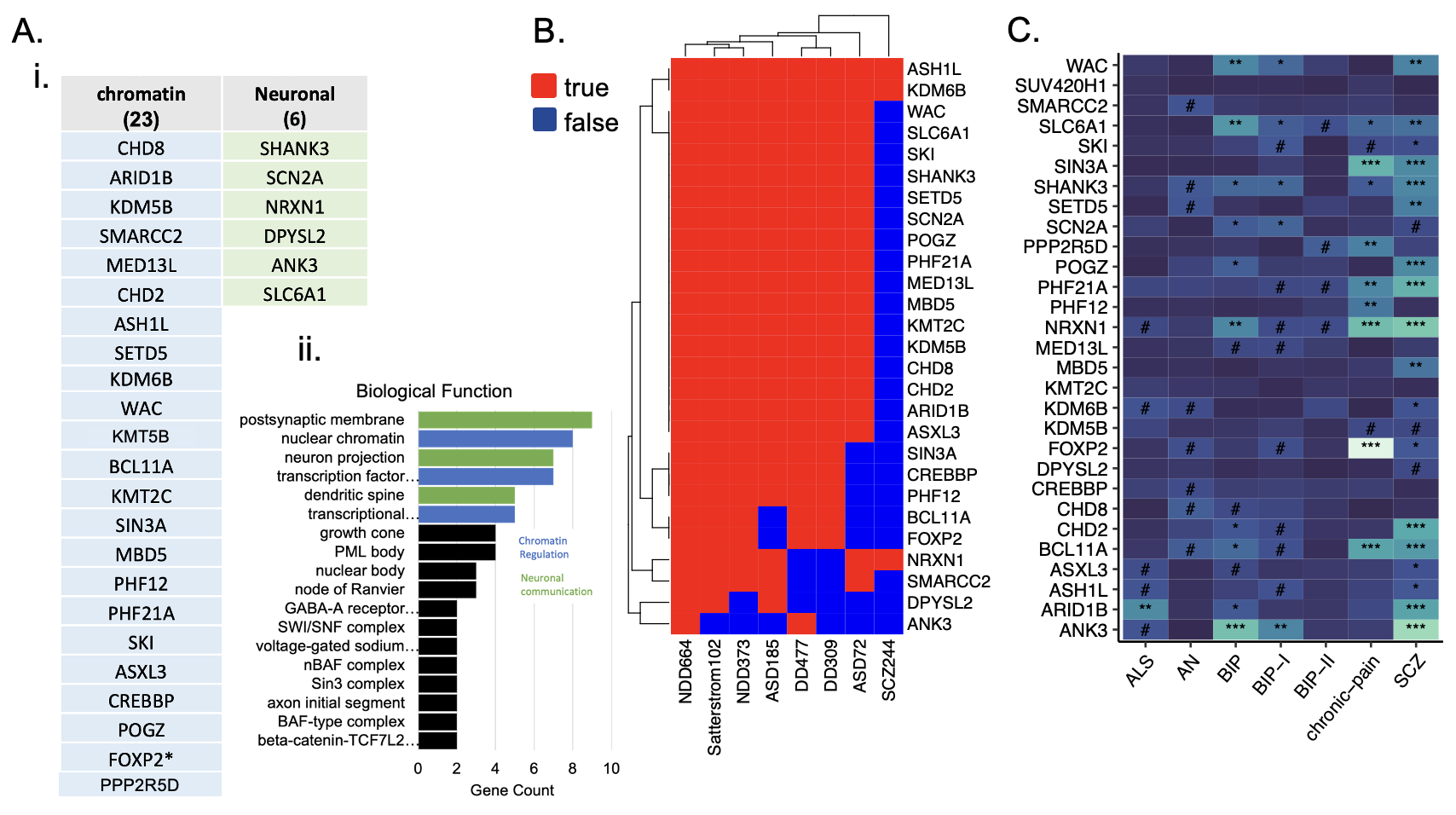
 **SI Figure 1. Characterization of 29 NDD rare variant genes. (A) (i)** Broad functional classification of genes selected for NDD scCRISPR-KO pool screen based on **(ii)** over-representation analysis of gene-ontology biological function gene sets. **(B)** Heatmap of gene-disorder associations from exome-sequencing studies across neurodevelopmental disorders corresponding with **SI Table 1** (Developmental Delay (DD), Neurodevelopmental disorders (NDD), autism spectrum disorder (ASD), and schizophrenia (SCZ). **(C)** Corresponding individual gene-GWAS associations from MAGMA. **^#^**nominal p-value<0.05, *****FDR<0.05, ******FDR<0.01, *****FDR<0.001.

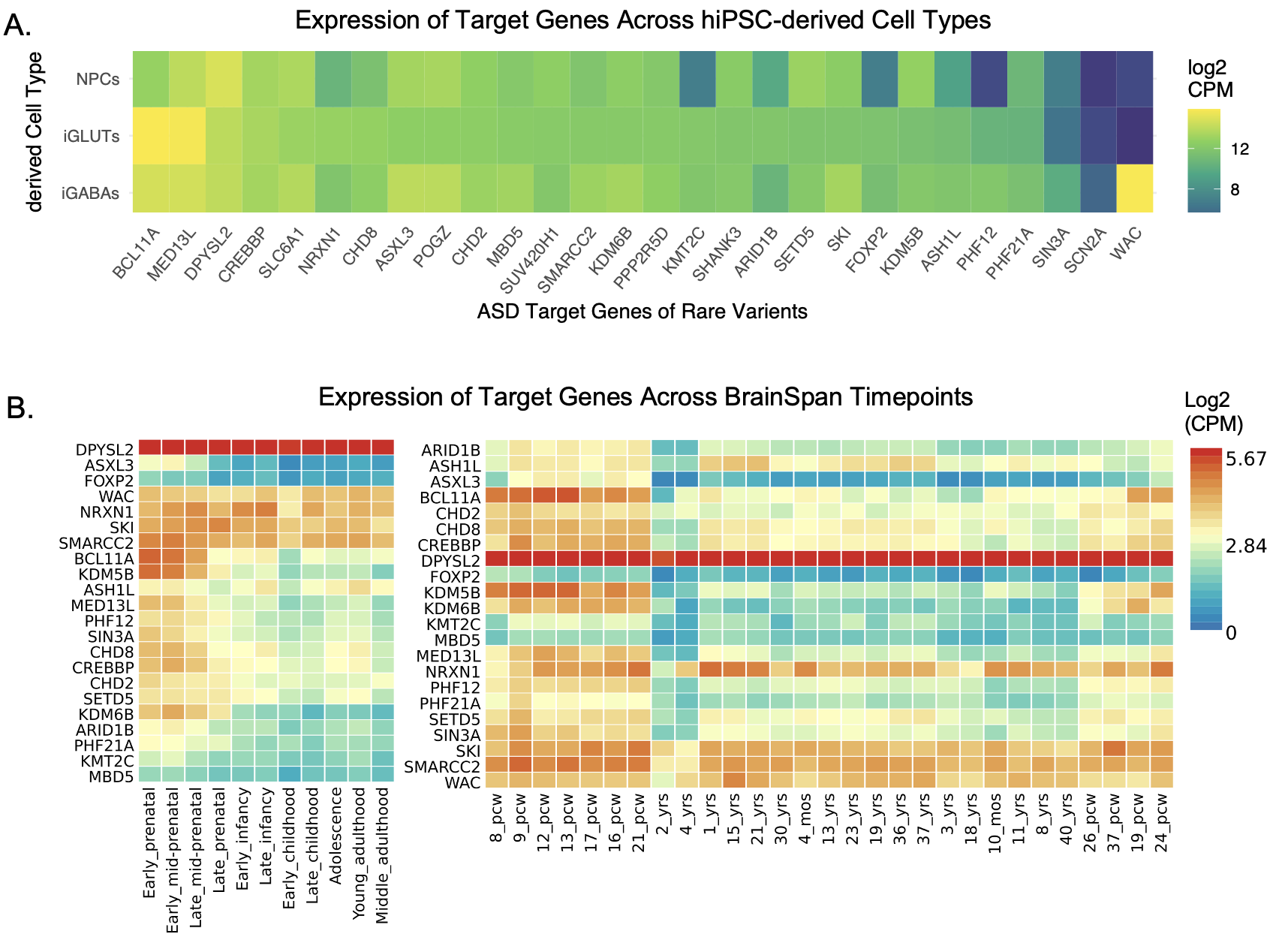
 **SI Figure 2. Characterization of gene expression of 29 NDD rare variant target genes in hiPSC neurons and the developing brain. (A)** Normalized log2(CPM) of 29 NDD risk genes across NPCs, iGLUTs, and iGABAs (ordered by average expression in iGLUTs) and **(B)** BrainSpan developmental stages pulled using FUMA**.**

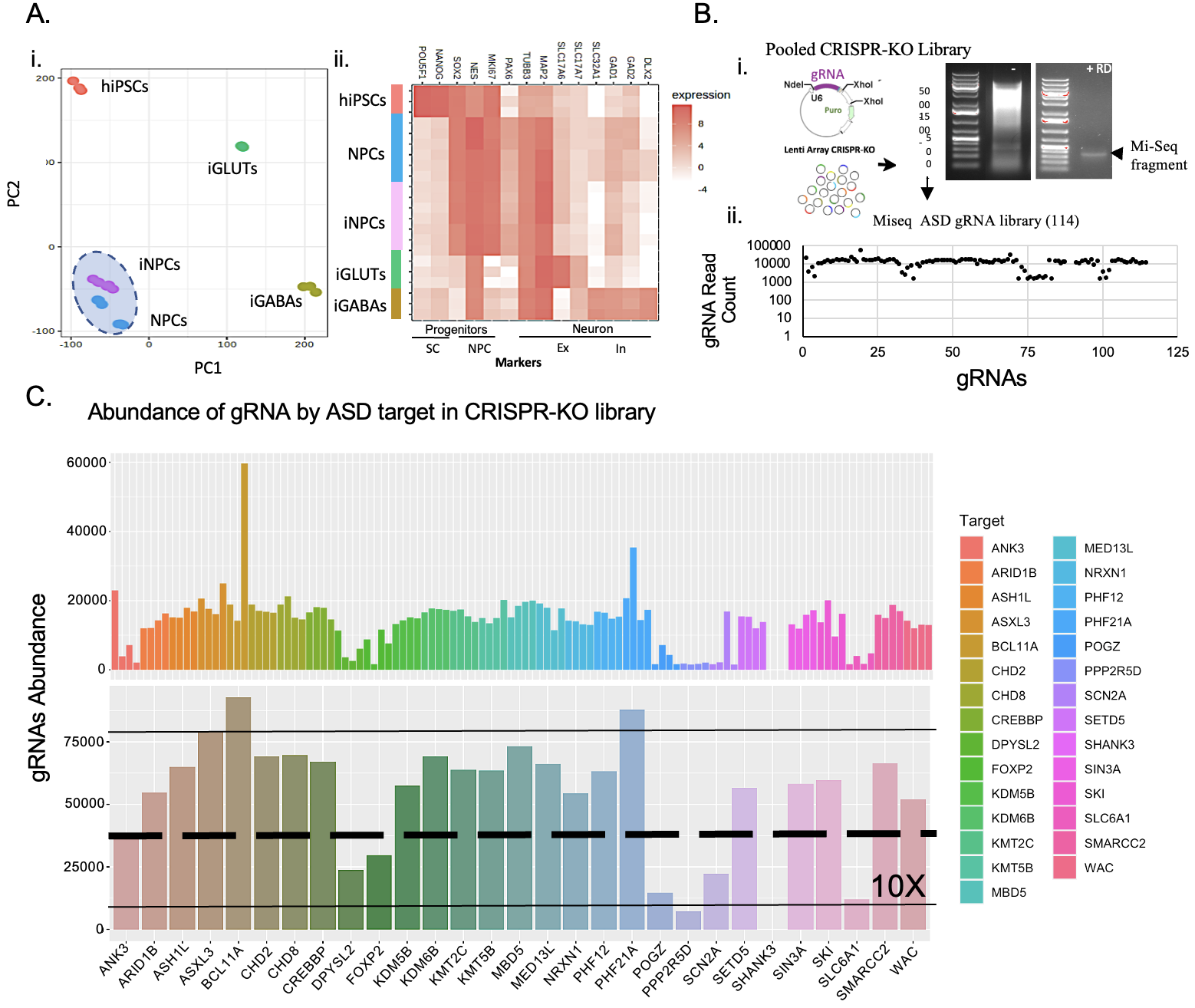
 **SI Figure 3. NDD pooled scCRISPR-KO screen validation. (A)** Characterization of gene expression from clonalized lines of hiPSCs, NPCs/iNPCs, iGLUTs, and iGABAs by **(i)** PCA and **(ii)** average expression of cell-type specific markers. **(B)** Diagram of the gRNA plasmid insertion site and gel electrophoresis of cloned pooled library restriction digested (RD) for Mi-seq validation. **(C)** Frequency of ASD pooled gRNA (counts) in library by Mi-seq.

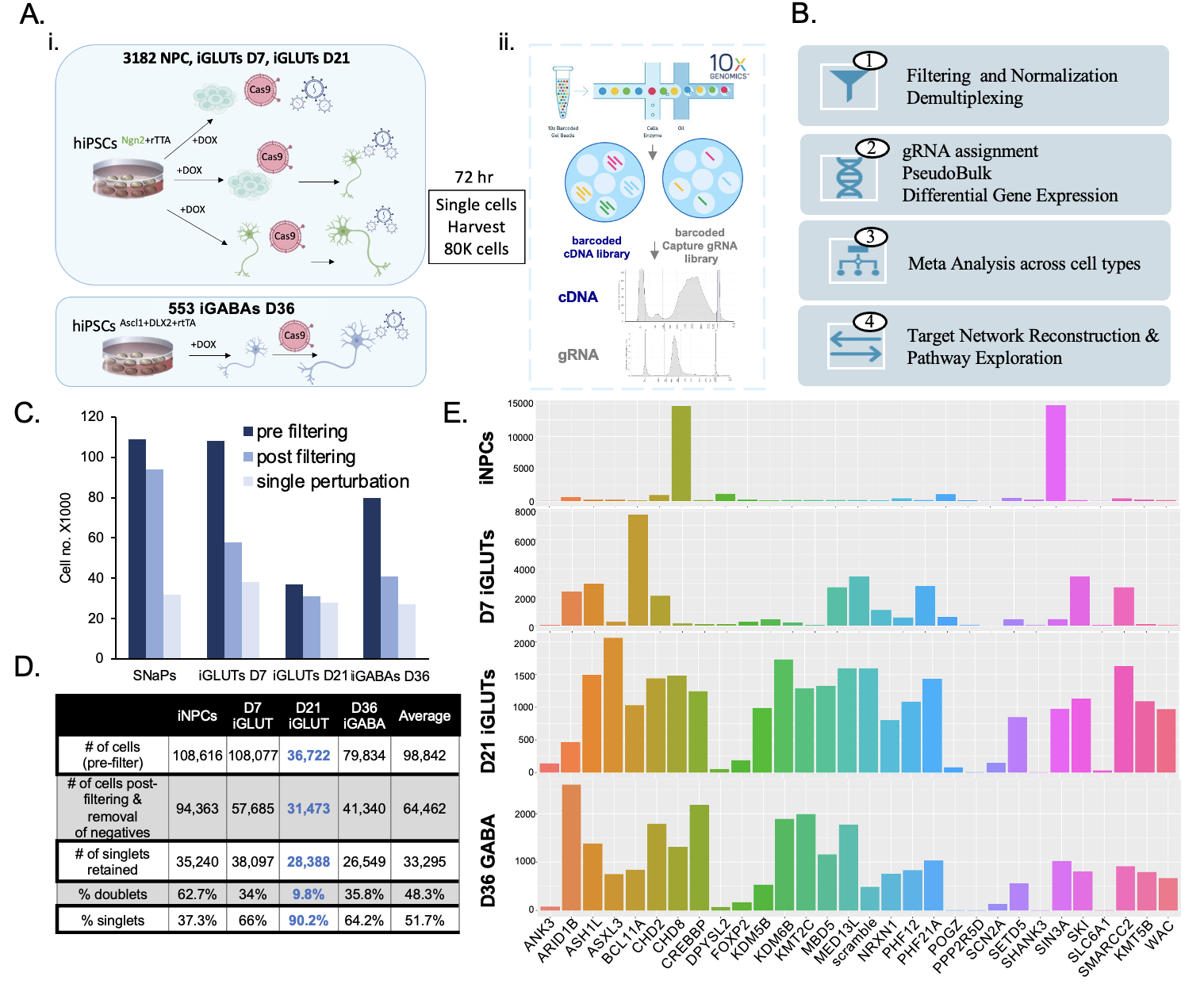
 **SI Figure 4. NDD scCRISPR-KO screen in hiPSC-derived iNPCs, iGLUTs, and iGABAs. (A)** Illustration scCRISPR-KO screen experimental set-up. Note iNPCs and iGLUTs in control 1 (3182-3-clone5) and iGABAs in control 2 (553-3-clone34). **(B)** Analytical process following 10x sequencing. **(C)** Number of cells sequences (pre-filtering), post-filtering, and post selection for single gRNA demultiplexing across cell-types. **(D)** Table of number of cells – including percent singlets and doublets by experiment. **(E)** Frequency of cell gRNA identity by experiment.

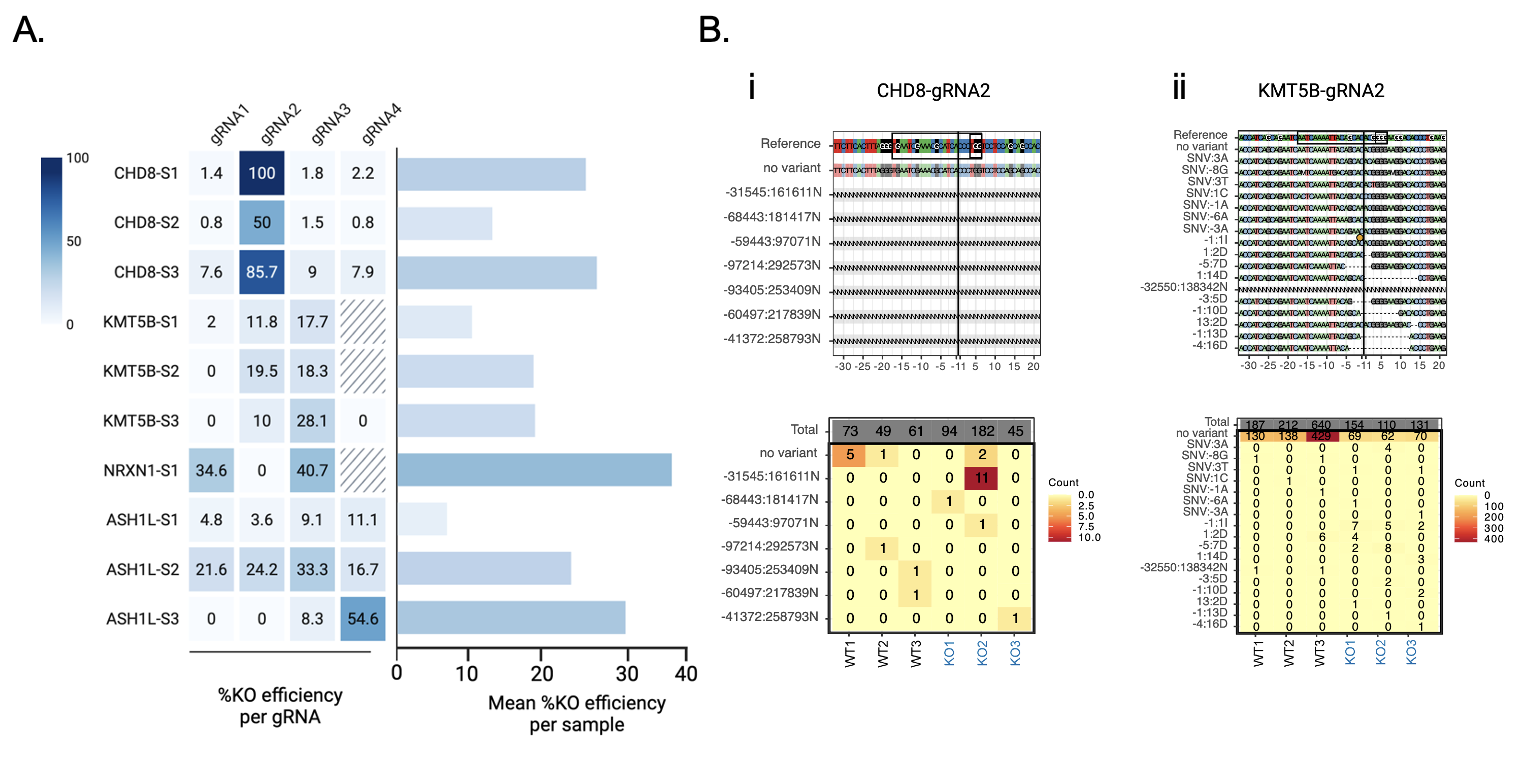
 **SI Figure 5. Validation of CRISPR-induced genome editing on selected genes with bulk RNAseq**. **(A)** CRISPR-editing efficiency of gRNAs across 4 NDD genes using bulk RNAseq. Mean efficiency was calculated by average over all non-zero gRNAs of corresponding sample. Max efficiency was estimated assuming all editing events occur independently. **(B)** Representative variant plots of highest editing efficiency gRNAs for each (i) *CHD8* and (ii) *KMT5B*. Top panel, variants of gRNA-targeting loci, color-coded by nucleotide. Variants were identified based on CIGAR strings (including gaps) and are annotated on the left. The gRNA targeting site and PAM sequence are highlighted with brackets. The x-axis represents the relative position to the predicted Cas9 cut site. Bottom panels show the counts of each variant site in Cas9-induced (KO) and non-induced (WT) samples

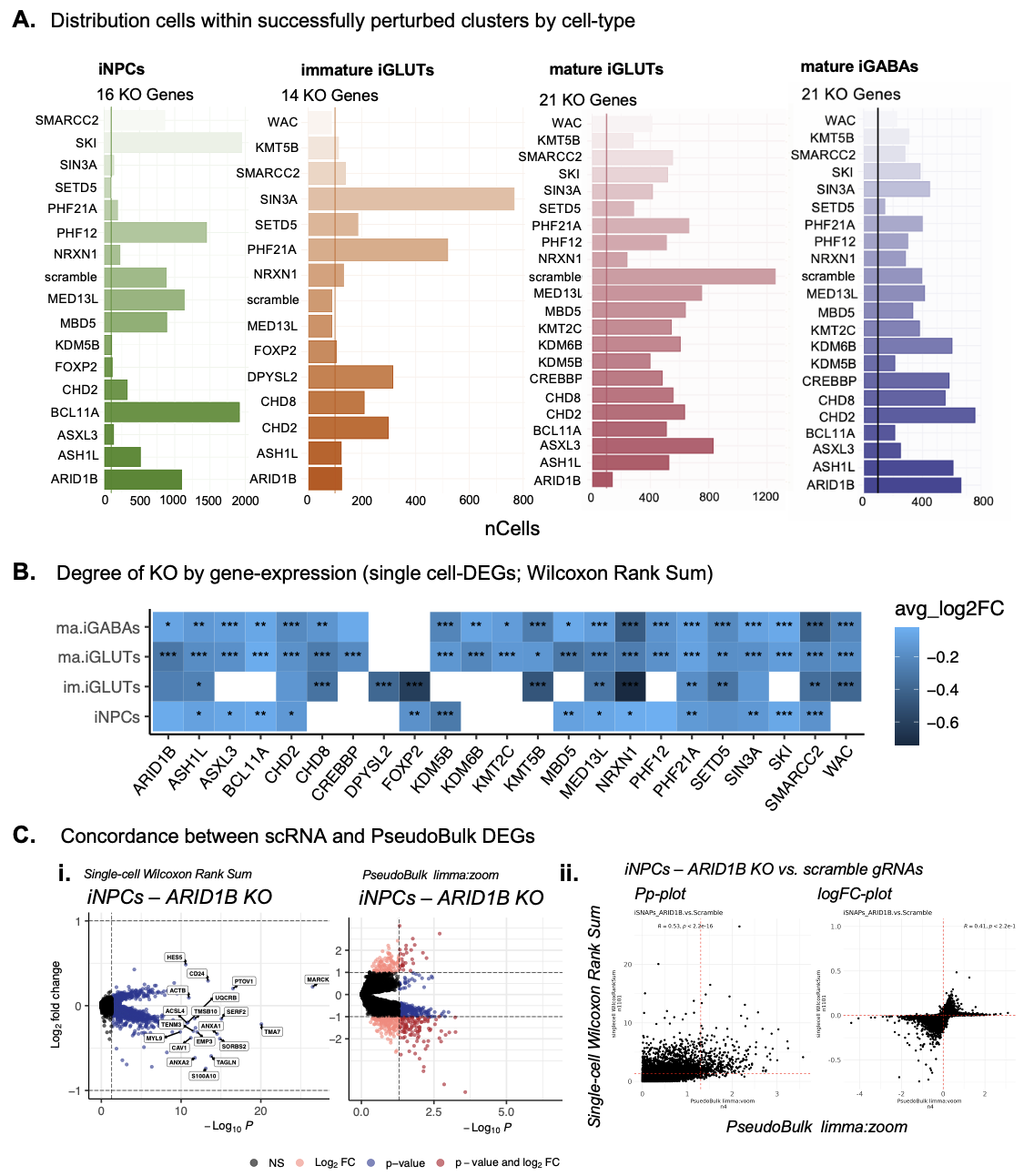
 **SI Figure 6. Characterization of perturbed cells and concordance between single-cell and pseudobulk differential gene expression analysis. (A)** Number of singlet cells assigned as perturbed by gRNA per cell-type after performing WNN and filtering clusters distinct from non-targeting scramble control clusters. **(B)** Degree of logFC downregulation of target genes across cell-types from Wilcoxon and auROC DEG analysis in single-cell data shows that perturbed cells have lower expression of the targeted NDD gene. **(C)** Examples of the concordance of differential gene expression analysis using Wilcoxon Rank Sum test in single-cell data and limma:voom in pseudo-bulked data. **(i)** Volcano plots of differential gene expression in *ARID1B* KO versus non-targeting samples in NPCs. **(ii)** Plots comparing p-values and logFC between differential gene expression analysis results at the single-cell level and pseudo-bulk level for *ARID1B*, *NRXN1*, and *CHD2* (Pearson’s correlation, Holm’s adjusted P-values).

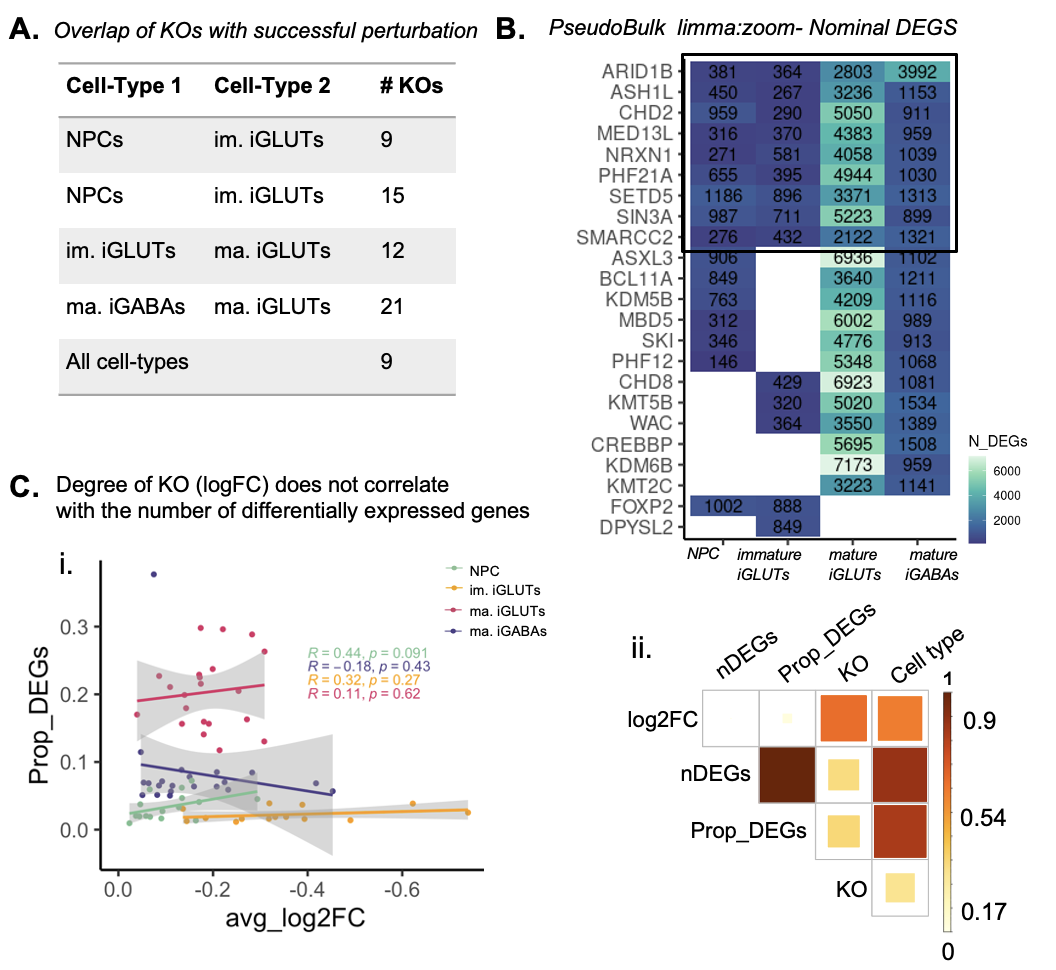
 **SI Figure 7. Degree of down-regulation of KO expression does not significantly correlate with downstream transcriptomic impacts. (A)** The number of KOs successfully resolved shared between cell-types - and thus allowing for direct comparison. **(B)** Transcriptomic impact of ASD/NDD gene KO across cell-type specific screens represented as the number of nominally significant (p<0.01) differentially expressed genes (DEGs). proportion of nominally significant DEGs in any cell-type (Pearson’s correlation, Holm’s adjusted P-values)**. (C)** There was no significant correlation between the average logFC of the target KO gene and the number of significantly differentially expressed genes. (i) Scatter plot of the average log2FC and proportion of differentially expressed genes (relative to the number of overall expressed gene) for each NDD KO colored by cell-type annotated with linear correlation coefficient and p-values. (ii) Heatmap of correlation coefficients across Cell type, KO gene, proportion of DEGS (prop_DEGs), absolute number ofDEGs (nDEGs), and the log2FC of the corresponding KO gene.

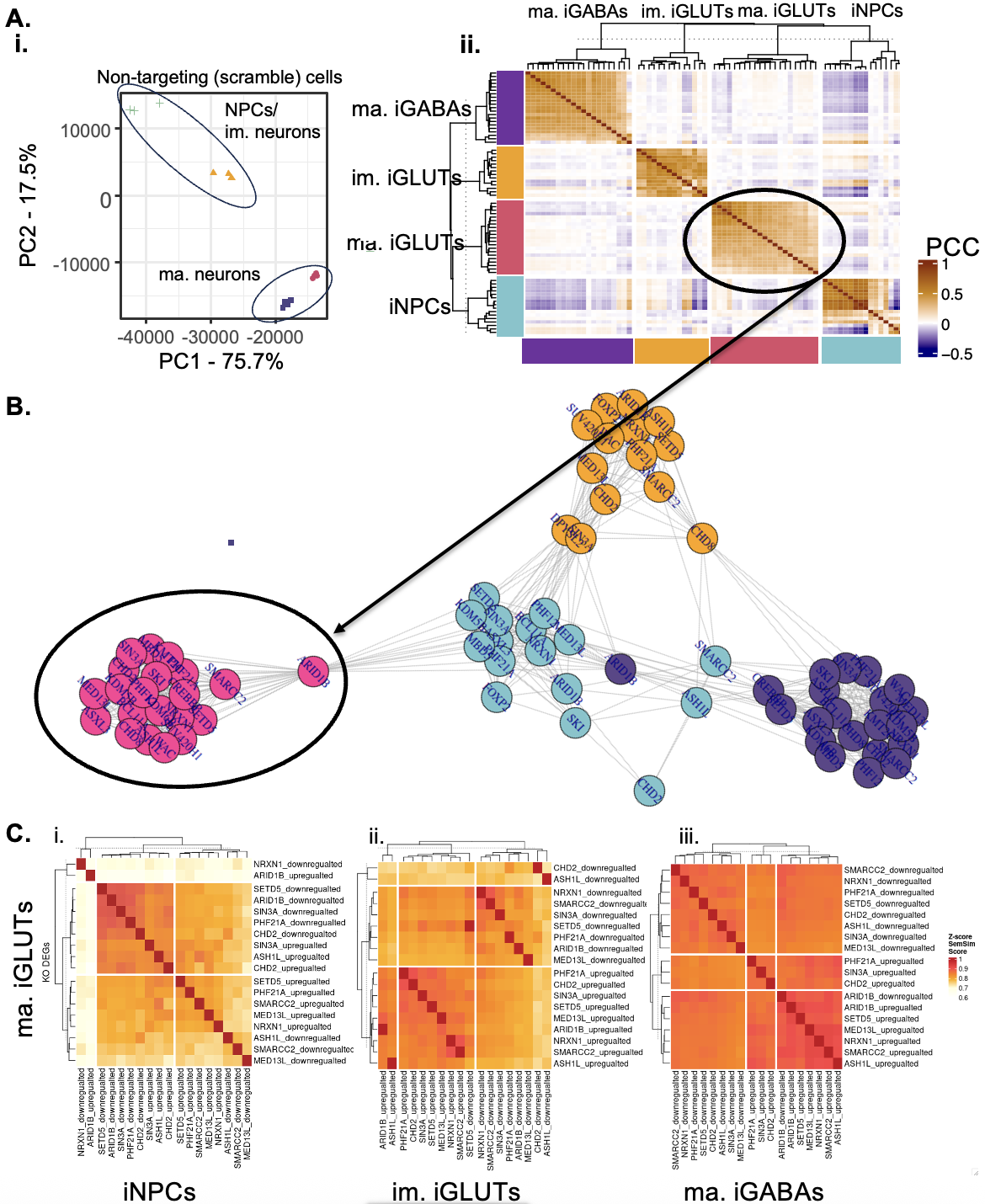
 **SI Figure 8. Stronger correlations between individual KOs in mature neurons are replicated after random sub-setting of controls. (A)** PCC correlation heatmap individual perturbation signatures**:** To validate if the high correlation within cell type was due to exactly the same cells being used as control populations for each KO, the difference in number of controls cell between cells types, or the greater difference between controls in immature cell-types as indicated by PCA **(i) d**ifferential expression was performed between KOs and randomly selected scramble controls. Briefly, for each gene, 50% (if number of pseudobulked sample cells > 50) or 80% (if number of pseudobulked sample cells < 50) of scramble cells were randomly selected differential expression analysis re-run. The process was repeated three times to avoid random selection bias and median of each gene logFC was used as the final logFC. Average overlap of random scramble cells across different genes is approximately 50%. (**ii)** Pearson’s correlation matrix of log2FC across all KOs was then calculated and plotted as heatmap. Greatest similarity remained within same cell type as when using all scramble controls. (**B)** PCC correlation network of individual perturbation signatures: Cross cell-type correlation network diagram across KO perturbations. Mature iGLUTs remained most isolated from the other three cell types, similar to what was observed in DEGs using all scramble controls. **(C)** Heatmap of z-score values of semantic similarity based on biological pathway membership of the significantly up and down-regulated DEGs for each KO compared. (iii) Mature iGABA and iGLUT KO target DEGs are the most functionally similar while (i) iNPC and mature iGLUTs KO DEGs are the least functionally similar.

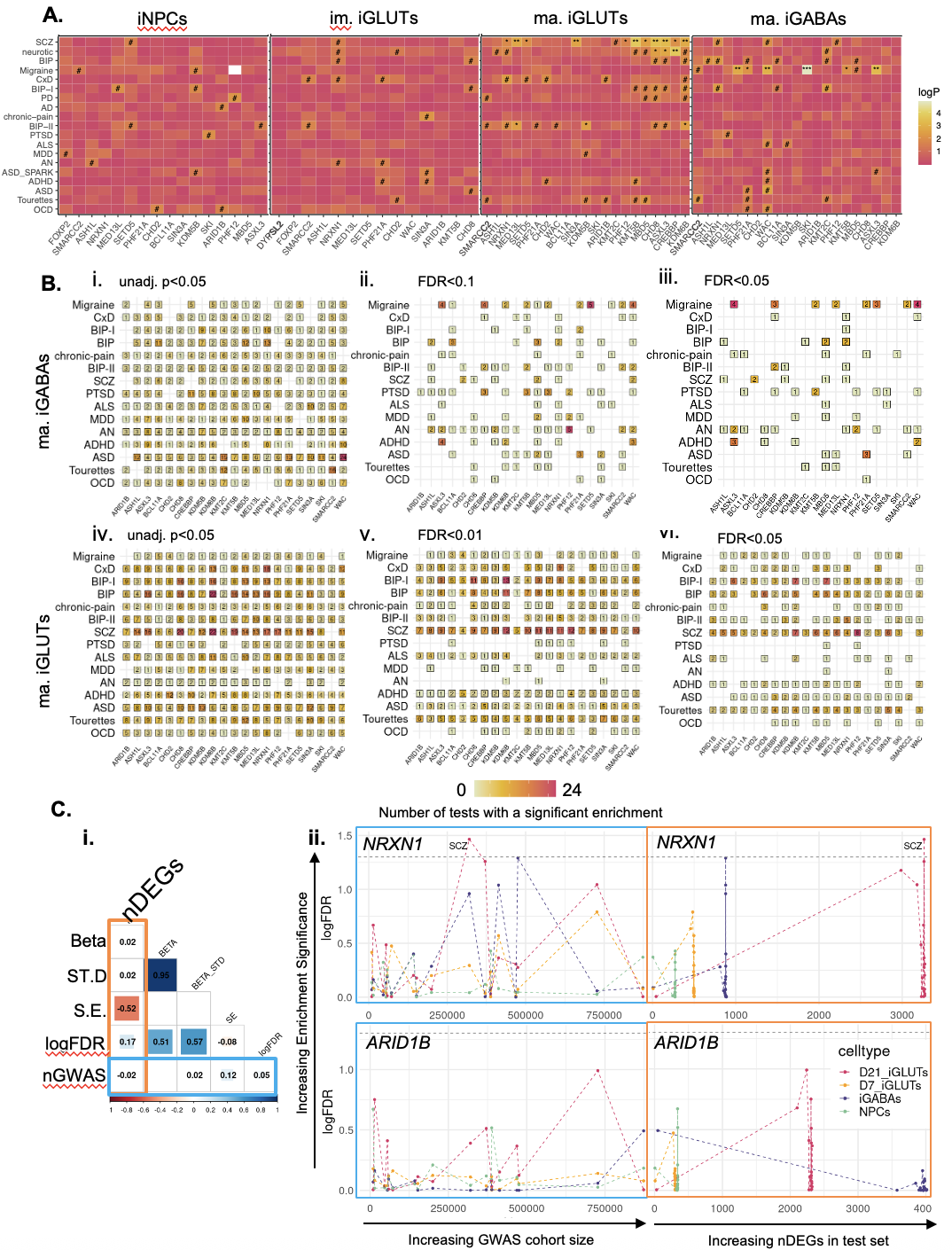
 **SI Figure 9. GWAS enrichments for individual KO effects across cell-types. (A**) Heatmap of MAGMA GWAS enrichments results for individual KO DEGs across cell-types reveals convergence on SCZ common variant risk genes in mature iGLUTs. **(B**) To test if this effect was due to SCZ risk genes being generally enriched in neurons or due to the size of the genesets for each GWAS – GWAS sets were filtered for genes expressed in each cell-type prior to enrichment testing and enrichment tests were performed after randomly down-sampling GWAS GeneSets to 100, 250, 500, 750, and 1000 genes. This process was then performed 10 times within each set size (i.e 50 tests for each GWAS). Strong associations with individual KO effects and SCZ and Bipolar genes remained across enrichments tests in mature iGLUTs. While strong association with migraine remained in iGABAs. Frequency tables showing the number of randomly down-sampled enrichment tests that were significant between KO DEGs and GWAS genesets across three significance thresholds in mature iGABAs and IGLUTs: **(i,iv)** unadjusted p-value <0.05, **(ii,v)** FDR <0.01, and **(iii,vi)** FDR<0.05. GWAS labels on the y-axis are order by the overall sample size from the GWAS studies. **(C**) **(i)** Since it is possible that GWAS sample size (nGWAS) and the number of DEGs (nDEGs) in a set may affect enrichment testing, we tested correlation between enrichment results (beta, standard deviation (ST.D), standard error (S.E) and found no significant correlations between enrichment significance and GWAS or gene set size. **(ii)** Scatter line plots visualizing relationships between GWAS cohort size and number of DEGs in enrichment sets for *NRXN1* and *ARID1B*.

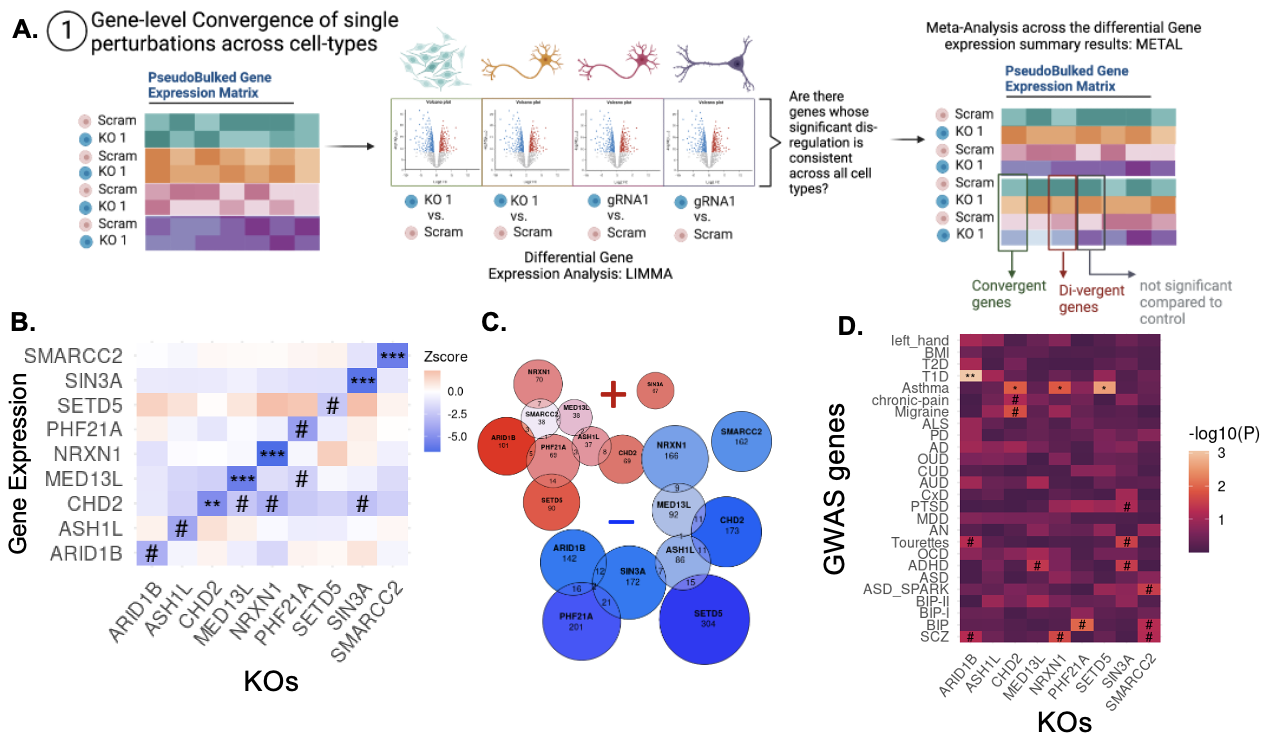
**SI Figure 10. NDDs show little transcriptomic convergence across cell-types and maturity. (A)** Schematic explaining KO-specific cross cell-type convergence at the individual gene level using differential gene expression meta-analysis. **(B)** Heatmap of meta-analyzed z-score effects on gene expression of targeted KO eGenes across perturbation phenotypes (#, unadjusted p-value =<0.05, *FDR<=0.05, **FDR<0.01, ***FDR<0.001). **(C)** Euler diagrams of overlapping convergent genes by eGene KO phenotype (nominal p-value <=0.05; Cochran’s Heterogeneity p-value > 0.05; positive (red) or (negative) direction of effect across all cell types). **(D)** Over representation analysis of KO-specific cross-cell type convergence and GWAS-risk associated genes.

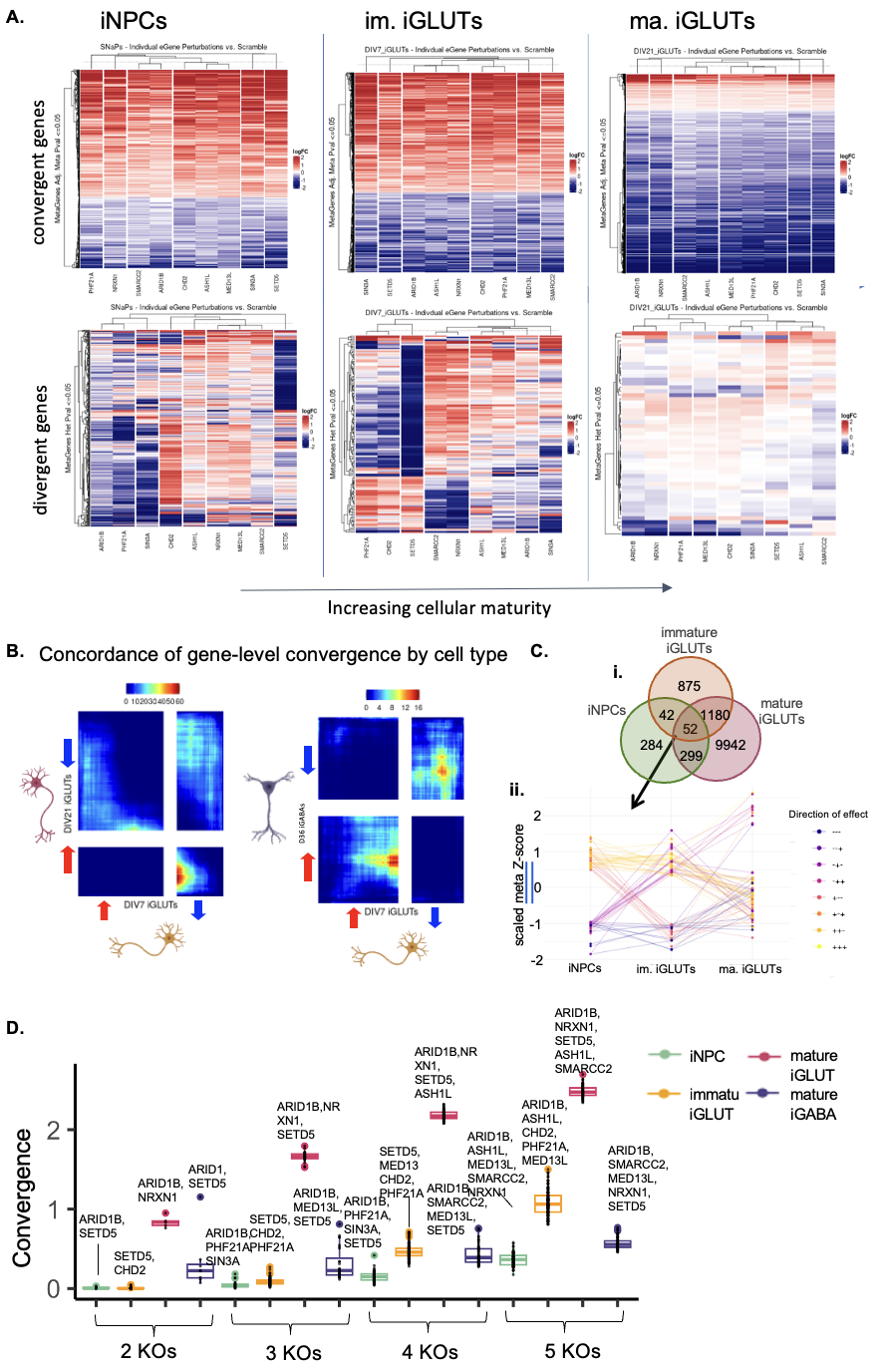

**SI Figure 11. Gene-level convergence across 9 NDDs is unique across cell-types and increases with maturity in iGLUT neurons. (A)** Heatmaps of convergent and divergent genes across 9 individual KOs in NPCs, immature iGLUTs, and mature iGLUTs). **(B)** Comparison over transcriptomic wide meta pFDR and z-scores using RRHO correlation across immature neurons (iGLUTs) and mature neurons (iGLUTs and iGABAs). **(C)** (i) Overlap of convergent genes across NPCs, immature iGLUT, and mature iGLUTs and (ii) their direction of effect in each cell-type. Scaled Z-scores of the 52 overlapping convergent genes across cell-type show that they have unique directions of effect.  **(D)** The top-most convergent KO combinations were unique to each cell-type. Boxplot of convergence magnitude for every tested combination of NDD genes ranging for sets of 2KOs to 5 KOs with the top convergent sets annotated.

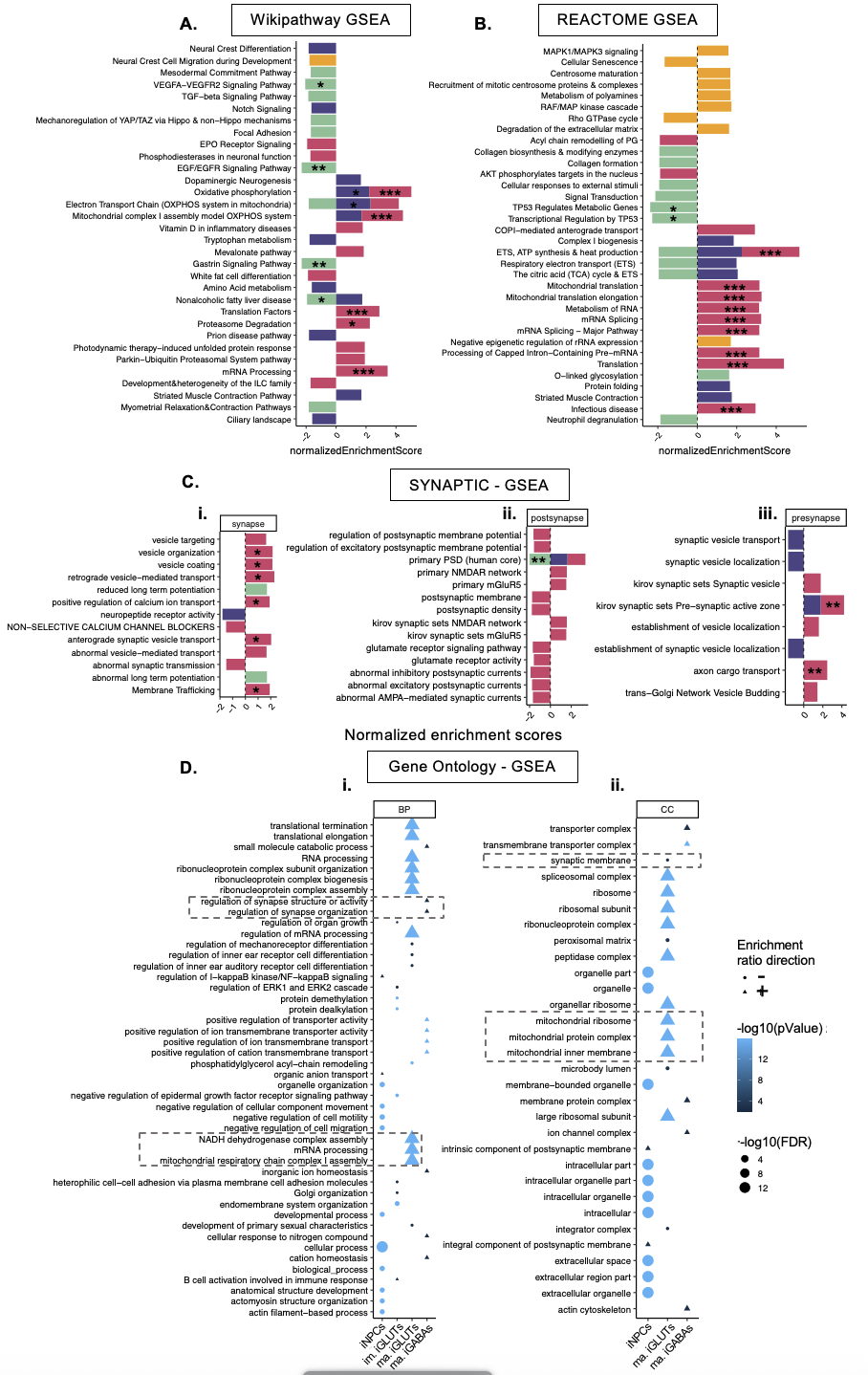

**SI Figure 12. Gene-level convergence across 9 NDDs is enriched for pathways involved in neural development, mitochondrial function, and translational regulation**. Gene set enrichment analysis of gene-level convergence across 9 NDD KOs (*ARID1B, ASH1L, CHD2, MED13L, NRXN1, PHF21A, SETD5, SIN3A, SMARCC2*) for each cell-type was performed using the **(A)** WikiPathway gensets, **(B)** Reactome genesets, **(C)** curated synaptic genesets, and **(D)** gene ontology pathways. Enrichments were filtered for nominal significance (unadjusted p-value <=0.05); FDR significance is indicated through annotations [FDR<=0.05*. FDR<=0.01**, FDR<=0.001***] (A-C) or size of the points.

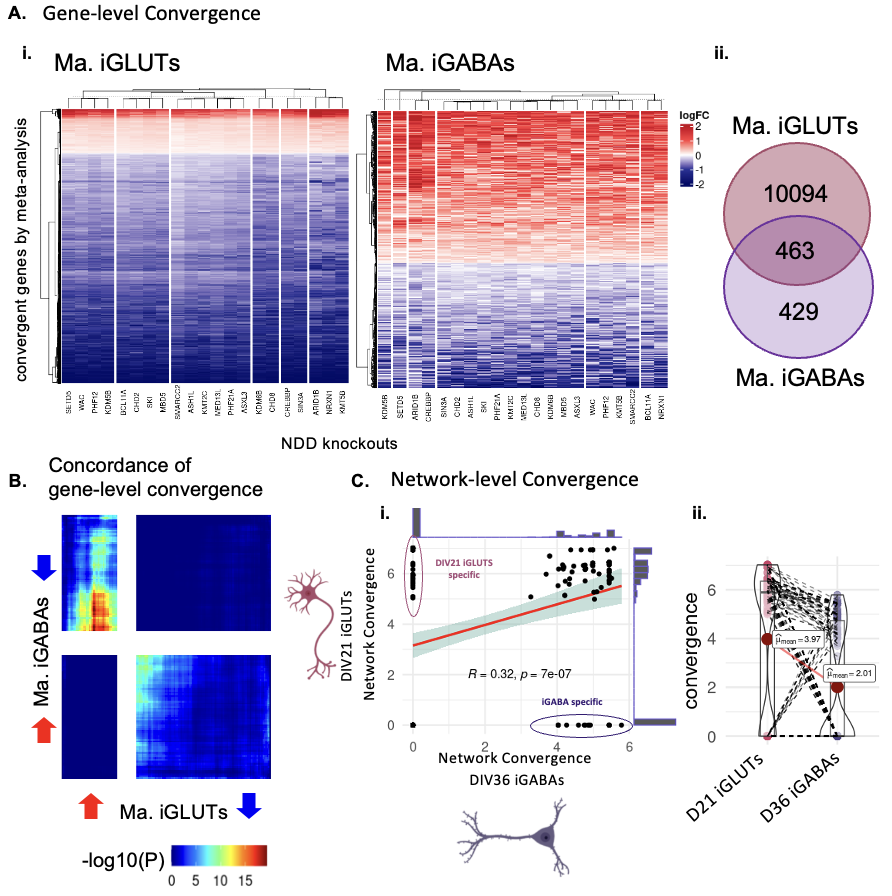
**SI Figure 13. Gene-level and network-level convergence across 21 NDDs in iGLUT and iGABA neurons. (A) (i)** Heatmaps of convergent and divergent genes across 21 NDD KOs between mature iGLUTs and iGABAs **(ii)** Overlap of convergent genes across mature iGLUTs and iGABAs. **(B)** Comparison over transcriptomic wide meta pFDR and z-scores using RRHO correlation between mature iGLUTs and iGABAs. **(C)** Comparison of the strength of network level convergence across different KO combinations – **(i)** While network convergence was generally strongly correlated between neuronal types, some cell-type specific networks exist and **(ii)** mature iGLUTs had significantly stronger mean convergence across all networks.

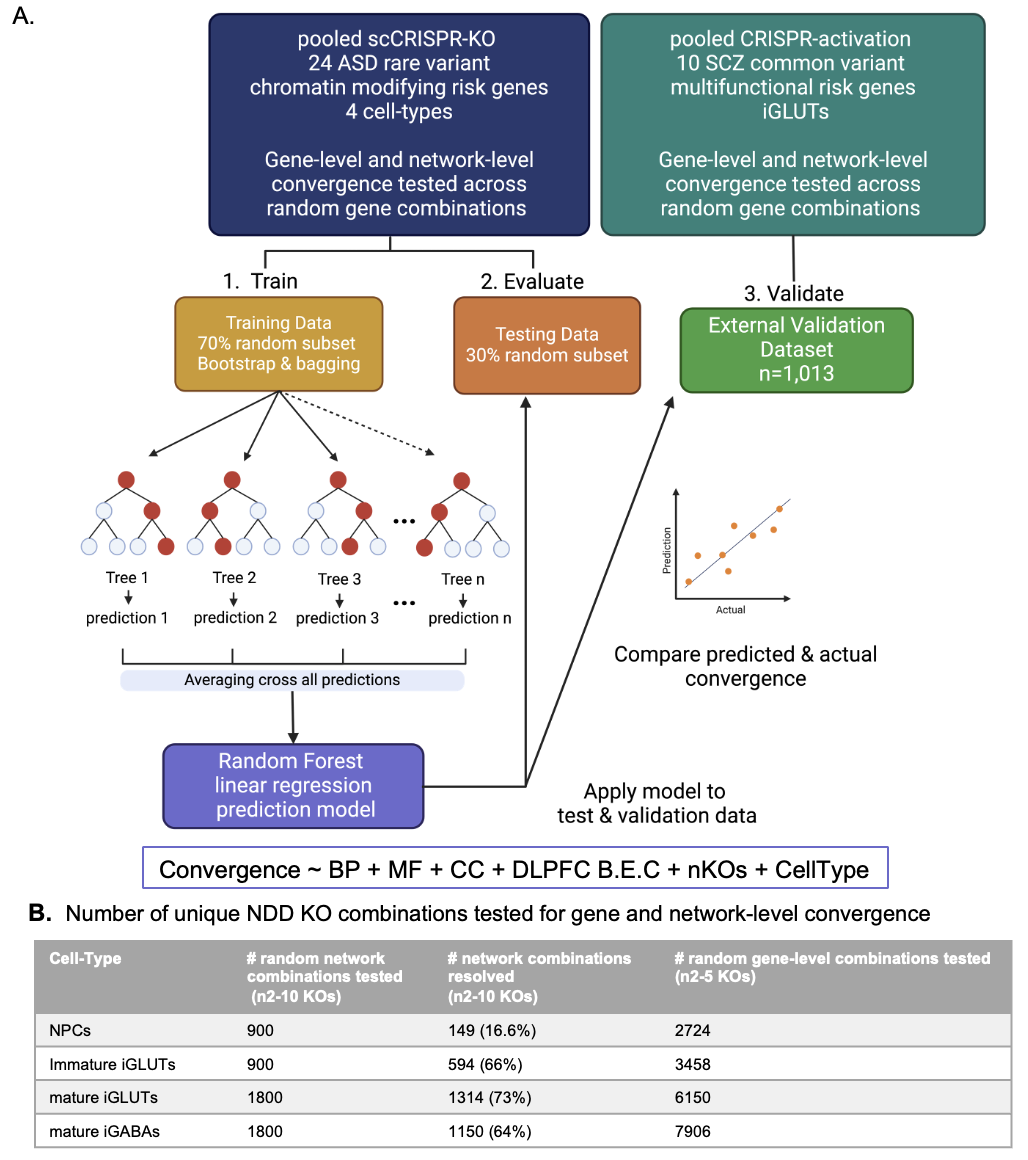
**SI Figure 14. Schema of random forest training model and external validation. (A)** Training random forest models for gene and network-level convergence. **(B)** Number of unique ASD KO combinations tested for gene and network-level convergence and used for random forest predictions. (***B.P score*** = semantic similarity of gene ontology (GO): Biological Process membership between KO genes; ***C.C. score*** = semantic similarity of GO: Cellular Component membership between KO genes; ***M.F score*** = semantic similarity of GO: Molecular functions membership between KO genes; ***B.E.C*** = dorsolateral prefrontal cortex expression correlations between KO genes; ***nKOs*** = number of KO genes tested for convergence). Corresponds with the analysis in **Figure 5**.

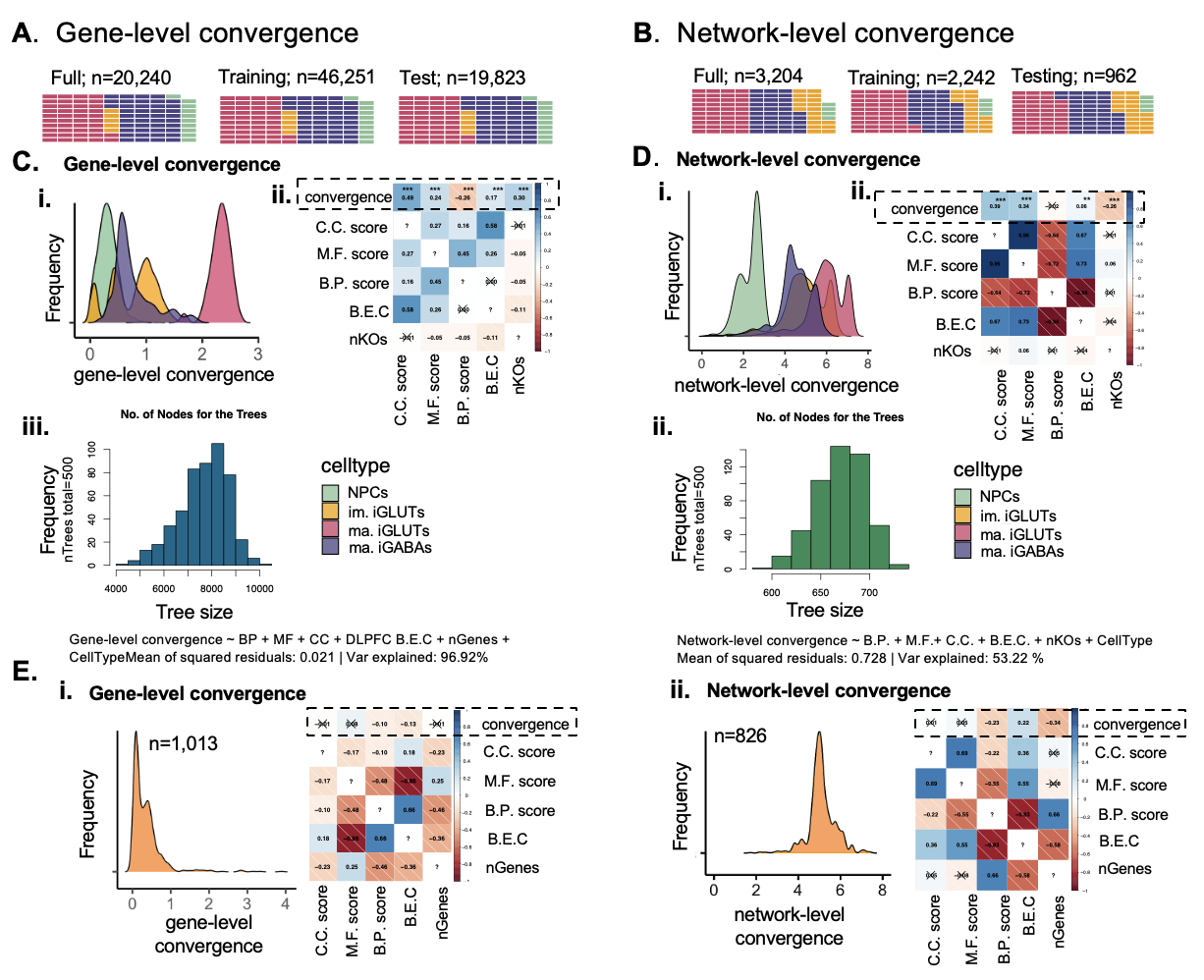
**SI Figure 15. Random forest training data characteristics and node frequency across trees. (A-B)** Functional similarity, brain co-expression, cell-type, and the number of KOs assayed strongly predicted gene-level convergence (98% variance explained; mean of squared residuals=0.02) and moderately predicted network-level convergence (53% variance explained; mean of squared residuals=0.73). Correlation of predictor variables and gene-level **(A-B)** The proportion of cell-types is balanced across the training sets and the testing sets and is representative the full data. **(C, D)** Distribution of gene-level **(C,i)** and network-level **(D,i)** convergence scores for each cell-type. Correlation of predictor variables and gene-level **(C,ii)** and network-level **(D,ii)** convergence. Number of nodes per tree in the random forest models (nTrees total=500) for **(C,iii)** gene-level convergence and for **(D,iii)** network-level convergence. **(E)** Validation of predictor models in an independent scCRISPRa screen of SCZ target genes. Distribution of convergence scores and their correlation with predictor variables at the gene **(E,i)** and network-level **(E,ii).** (***B.P score*** = semantic similarity of gene ontology (GO): Biological Process membership between KO genes; ***C.C. score*** = semantic similarity of GO: Cellular Component membership between KO genes; ***M.F score*** = semantic similarity of GO: Molecular functions membership between KO genes; ***B.E.C*** = dorsolateral prefrontal cortex expression correlations between KO genes; ***nKOs*** = number of KO genes tested for convergence). Data corresponds with the analysis in **Figure 5**.

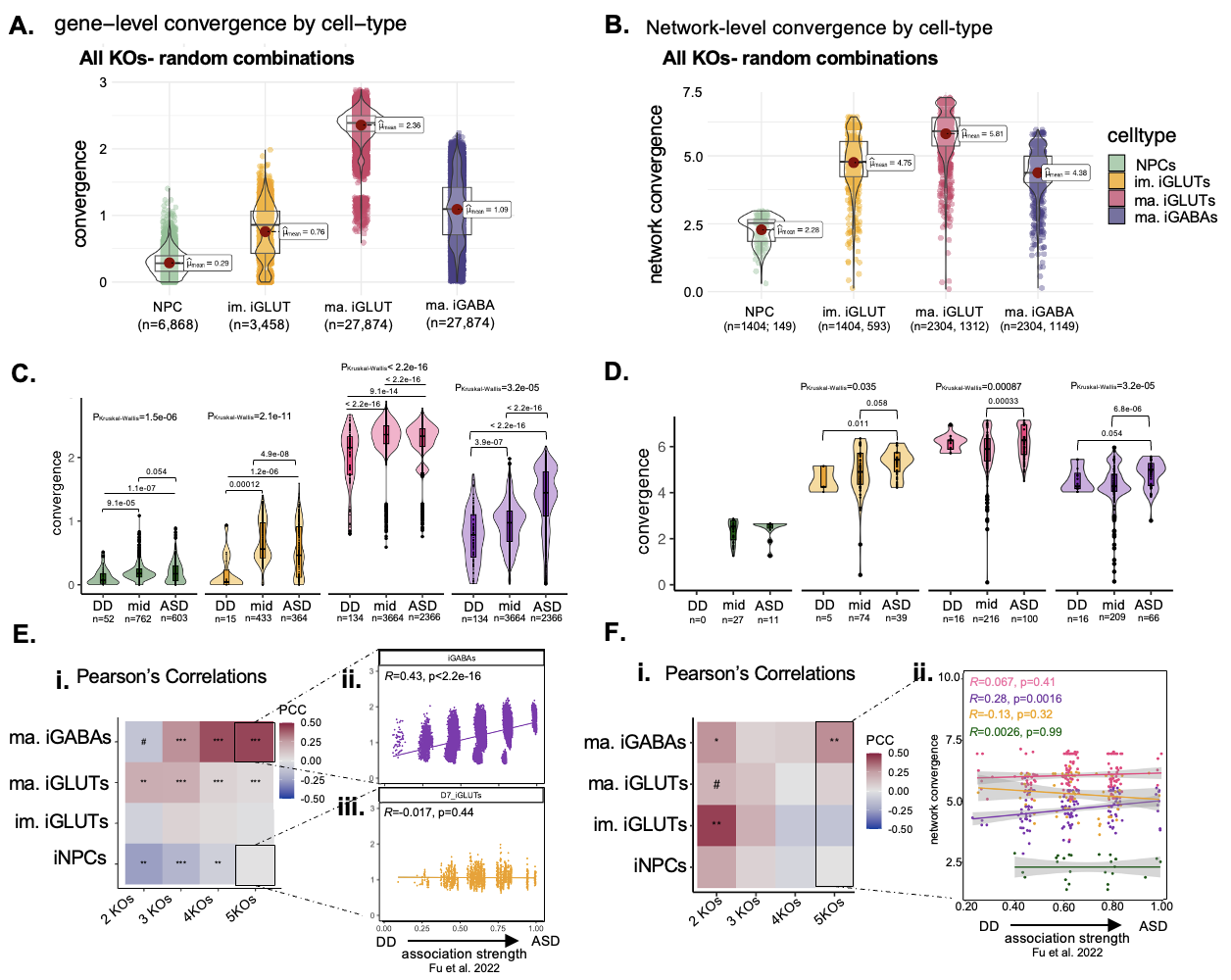
**SI Figure 16. NDD risk genes with strong ASD associations resolve greater convergence across cell-types. (A)** Gene-level and **(B)** network-level convergence remain highest in mature iGLUTs when looking across all NDD gene perturbations. **(C,D)** Transcriptomic convergence is greater between KOs with stronger associations with ASD compared to DD across all cell-types. Associations between the ASD risk specificity and convergence was performed by calculating the mean ASD posterior probability (p.p.) from Fu et al. 2022 across KOs in a set; KO sets were stratified into three groups based on these scores (DD = mean ASD p.p. <=0.1; ASD = mean ASD p.p. >=0.9; mid= mean ASD p.p. <=0.55 & >=0.45) and differences in gene-level **(C)** and network-level **(D)** convergence tested using non-parametric Kruskal-Wallis test, with pairwise comparisons performed with Wilcoxon’s test. For network-level convergence in iNPCs, comparisons between DD and ASD sets were not possible as none of the DD KO sets resolved convergent networks –an observation further suggesting that DD sets are less convergent. **(E,F)** Associations between ASD risk specificity and convergence by cell-type and the number of KOs in a set as assessed through linear correlation (Pearsons Correlation Coefficient (PCC) or “R”). **(E)** ASD p.p was significantly positively associated with gene-level convergence in mature neurons, especially iGABAs, but this was influenced by the number of KOs in a set. For example, gene-level convergence across 5 KOs was significantly positively correlated with ASD specificity in **(E, ii)** iGABAs (R=0.43, p<2.2e-16) but not in **(E, iii)** immature iGLUTs (R=0.017, p=0.44). Likewise, network-level convergence in sets of 5 KOs was positively correlated with ASD specificity in **(F, i-ii)** iGABAs (R=0.28, p=0.0016) but not immature iGLUTs (R=-0.13, p=0.32). The lack of clear correlation patterns suggest that while mean convergence is higher across genes with stronger ASD associations **(C,D),** this relationship is not linear and likely influenced by other shared factors between risk genes.

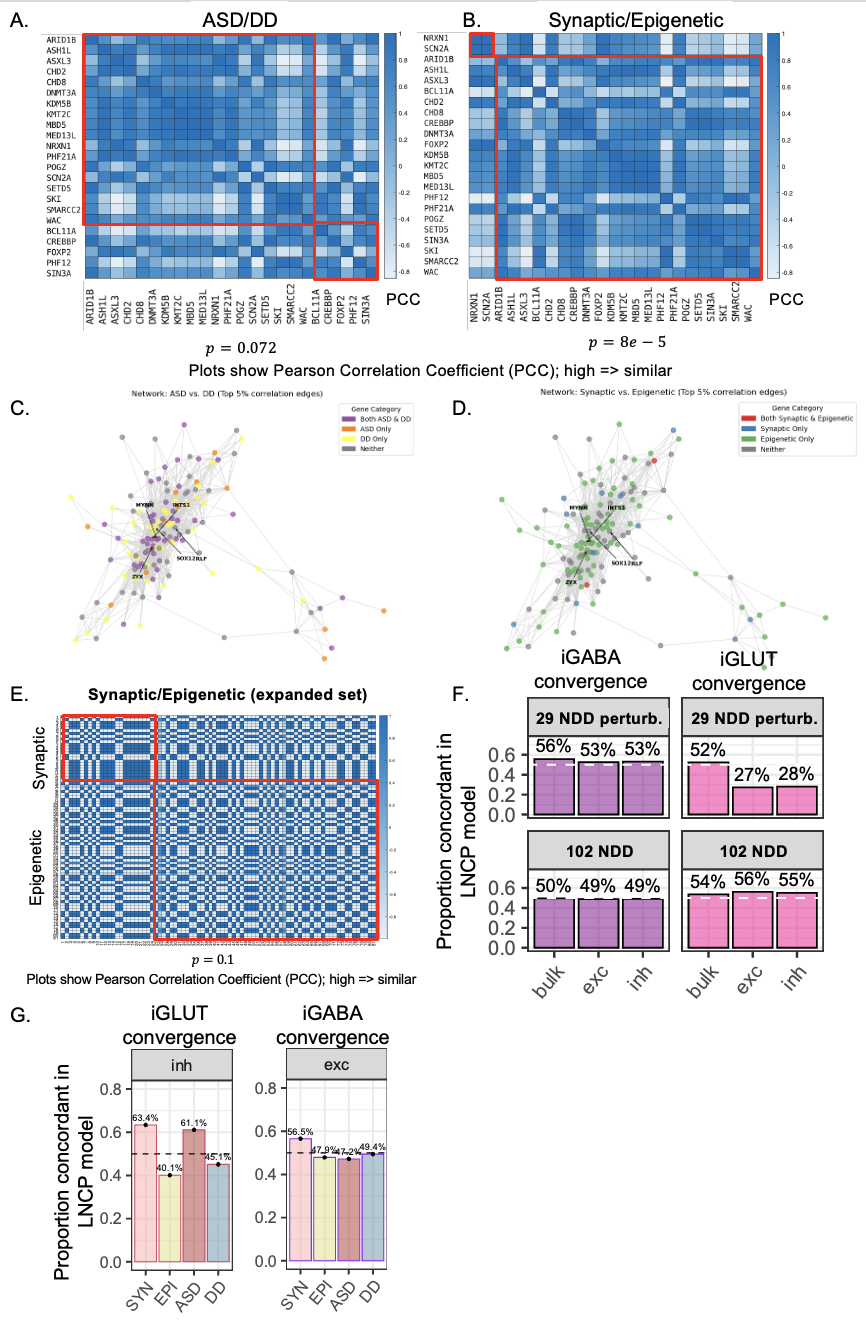

**SI Figure 17. Further analysis of LNCTP convergent genes.** Predicted *in silico* perturbation effects were significantly more correlated within categories defined by **(A)** disorder association (ASD – DD; p=0.072) and **(B)** function (synaptic – epigenetic; p=8x10-5). Heatmaps show the correlations of the LNCTP predicted *in silico* log fold-changes across PsychENCODE individuals for ASD/DD and Synaptic/Epigenetic CRISPR perturbations respectively (p-values, 1-tailed t-test for decreased correlation between perturbation categories). **(C,D)** Network visualization of the proximity of functional classes of *in silico* convergent genes, where adjacency represents co-expression of bulk expression across PsychENCODE subjects. Colored by disorder association **(C)** or functional association **(D)**. **(E)** Repeat of analysis in (A,B) with the expanding list of 102 NDD perturbations (p=0.1). **(F)** Repeat of the analysis in Fig. 5B, using all 102 NDD perturbations. **(G)** Repeat of the analysis in Fig. 5C, using all 102 NDD perturbations showing concordance between *in silico* convergence in inhibitory neurons with convergence in iGLUTs and *in silico* convergence in excitatory neurons with convergence in iGABAs.

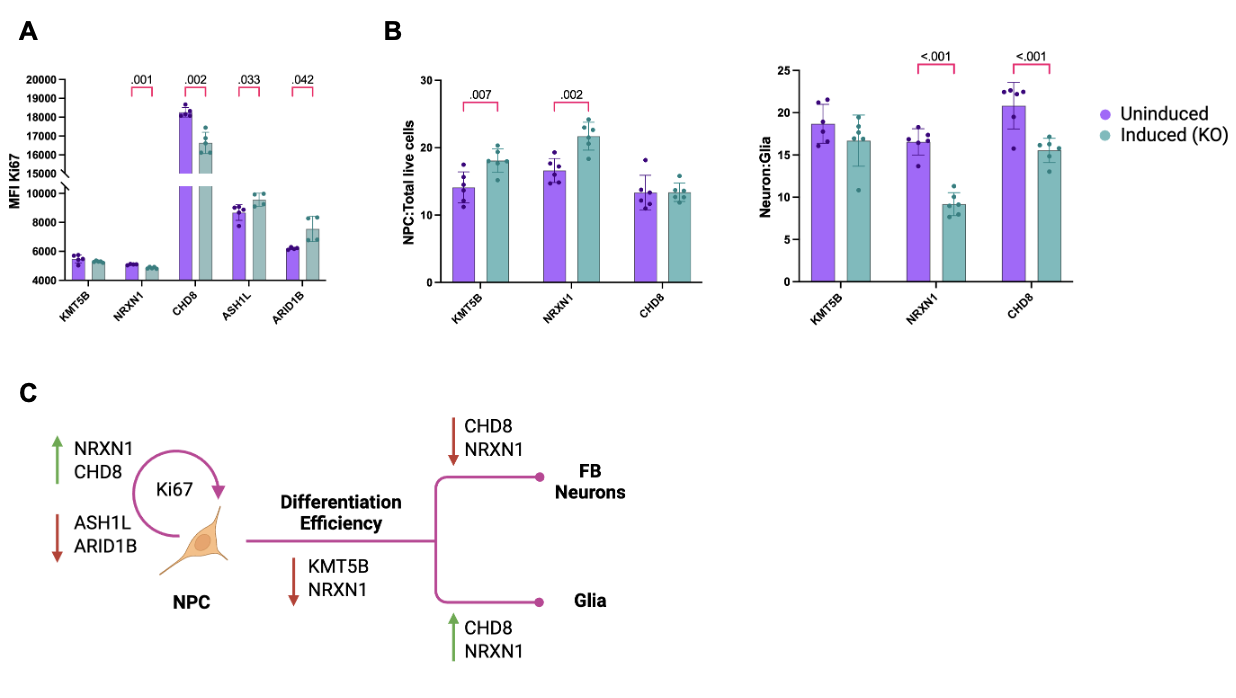
**SI Figure 18. Impact of NDD KO on proliferation and neurogenesis in NPCs and neurons. (A**) Proliferation assessment of NPCs using Ki-67 staining. NPCs were transduced with a pooled gRNA library containing four gRNAs per target gene targeting selected NDD genes (*KMT5B, NRXN1, CHD8, ASH1L, ARID1B*) and analyzed 7 days after Cas9 activation with doxycycline (dox). Flow cytometry analysis revealed significant changes in the median fluorescence intensity (MFI) of Ki-67-FITC. (**B**) Neurogenesis and gliogenesis assessment in differentiated forebrain cultures. NPCs were differentiated into neurons and glia, and the ratio of NPCs to total live cells and neurons to glia was quantified by flow cytometry. Error bars represent standard deviation (SD). Each dot represents an independent replicate. Unpaired t-test with Welch correction; p-values corrected for multiple comparisons using FDR. (**C**) Summary schematic of the findings. CHD8 and NRXN1 KO promote gliogenesis and NPC proliferation, while ASH1L and ARID1B KO reduce NPC proliferation. KMT5B KO reduces differentiation efficiency but does not significantly alter neuron-to-glia ratios.

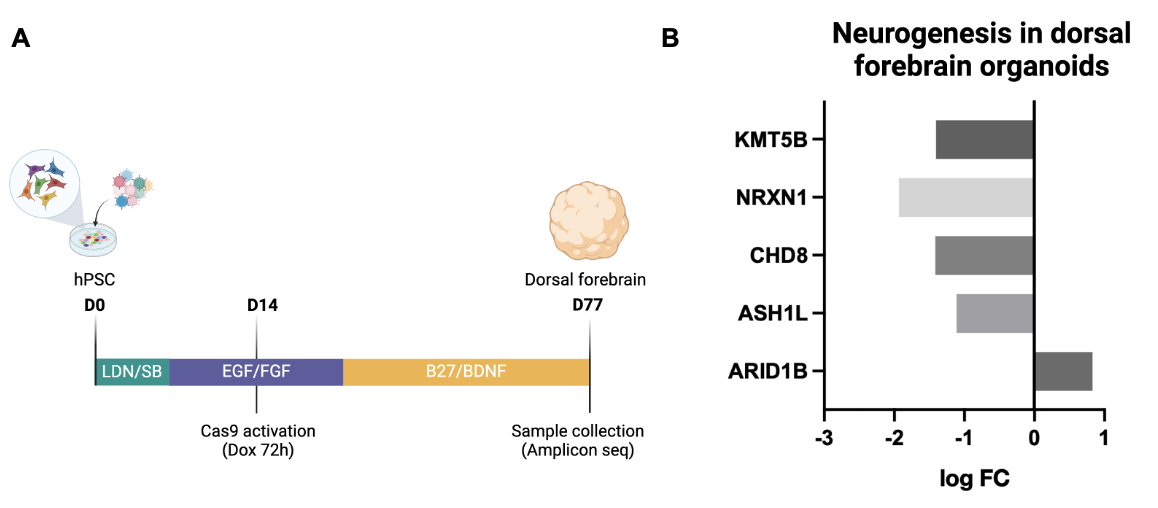
**SI Figure 19. Pooled CRISPR screen to uncover the NDD KO effects on cortical organoid development. (A**) Schematic representation of the experimental workflow. H1-hESC-iCas9 cells were transduced with a pooled gRNA library targeting five NDD genes (*KMT5B, NRXN1, CHD8, ASH1L, ARID1B*) and 20% non-targeting controls. Cas9 was activated on Day 14 via doxycycline (2 µg/mL) for 72 hours and maintained in culture for ~80 days. Organoids were subsequently processed for gRNA enrichment analysis using PCR amplicon sequencing to assess the relative abundance of gRNAs targeting each gene. **(B**) Bar plot showing the log fold change (logFC) of gRNA abundance relative to non-targeting controls. Negative logFC values indicate depletion of gRNAs targeting genes associated with reduced neurogenesis or cell survival (e.g., CHD8 and KMT5B), while positive logFC values (e.g., ARID1B) indicate enhanced proliferation or survival. These results demonstrate distinct roles for NDD genes in regulating organoid developmental dynamics, with ARID1B associated with increased proliferation and KMT5B and CHD8 contributing to neuronal depletion.

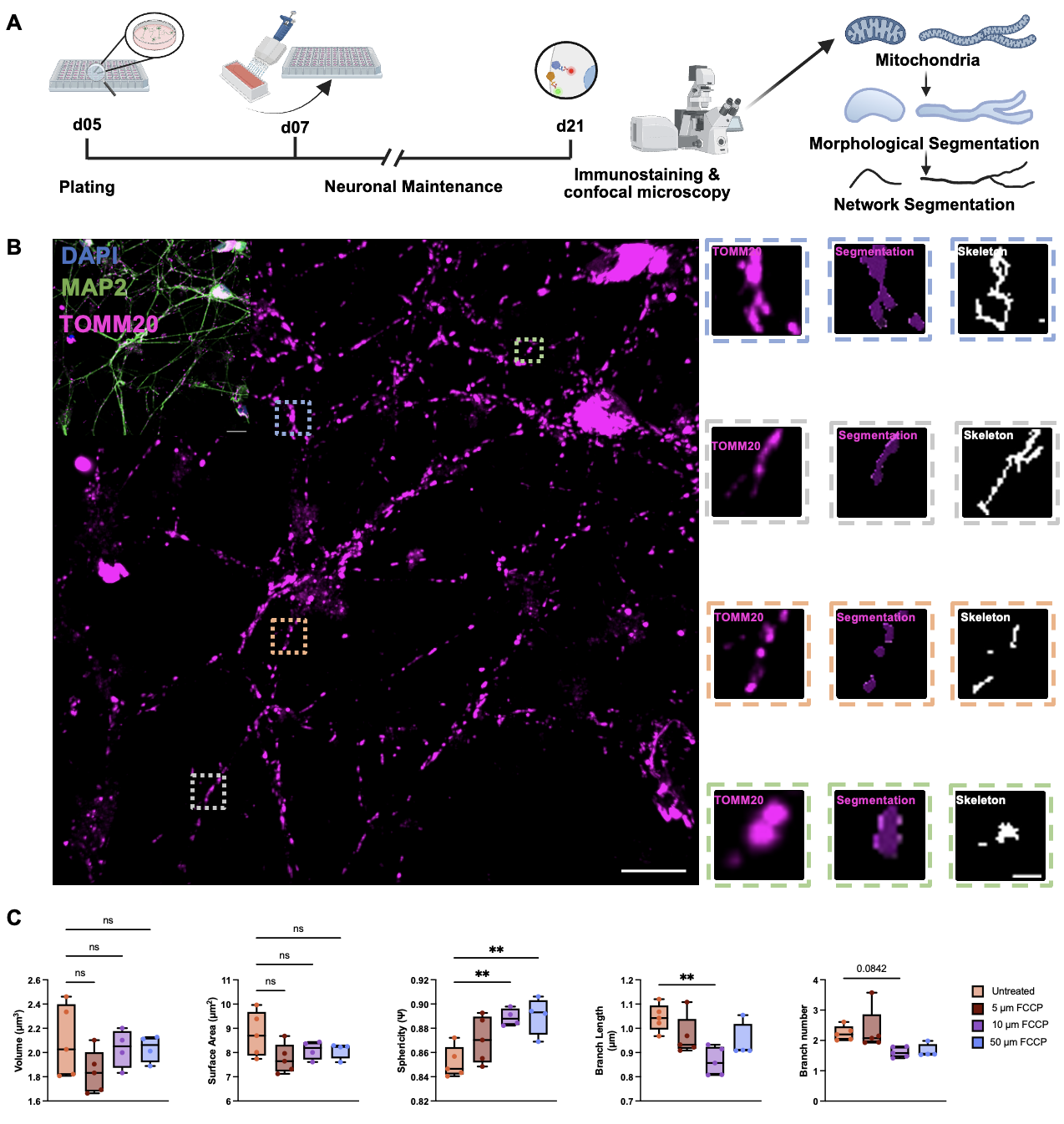
**SI Figure 20. High resolution microscopy captures diversity of mitochondrial morphology in iGLUTs and is sensitive to mitochondrial pharmacological insults. A)** Timeline of culture and microscopy experiments **B)** Representative image of iGLUTs labelled with mitochondria (TOMM20, magenta). Inset: merged representative image of iGLUTs with MAP2 (green) and DAPI (blue). Representative images of networked (blue), fused(grey), fragmented (yellow) and swollen mitochondria (green). Mitochondria are volumetrically segmented and skeletonized for morphological and network analysis. Scale 10 μm, 1 μm**. C)** Microscopy pipeline is sensitive, capturing dose-dependent mitochondrial fragmentation following FCCP treatment. FCCP treatment causes dose-dependent mitochondrial fragmentation (decreases in mitochondrial sphericity and increases in branch length) independent of changes in mitochondrial volume and surface area.

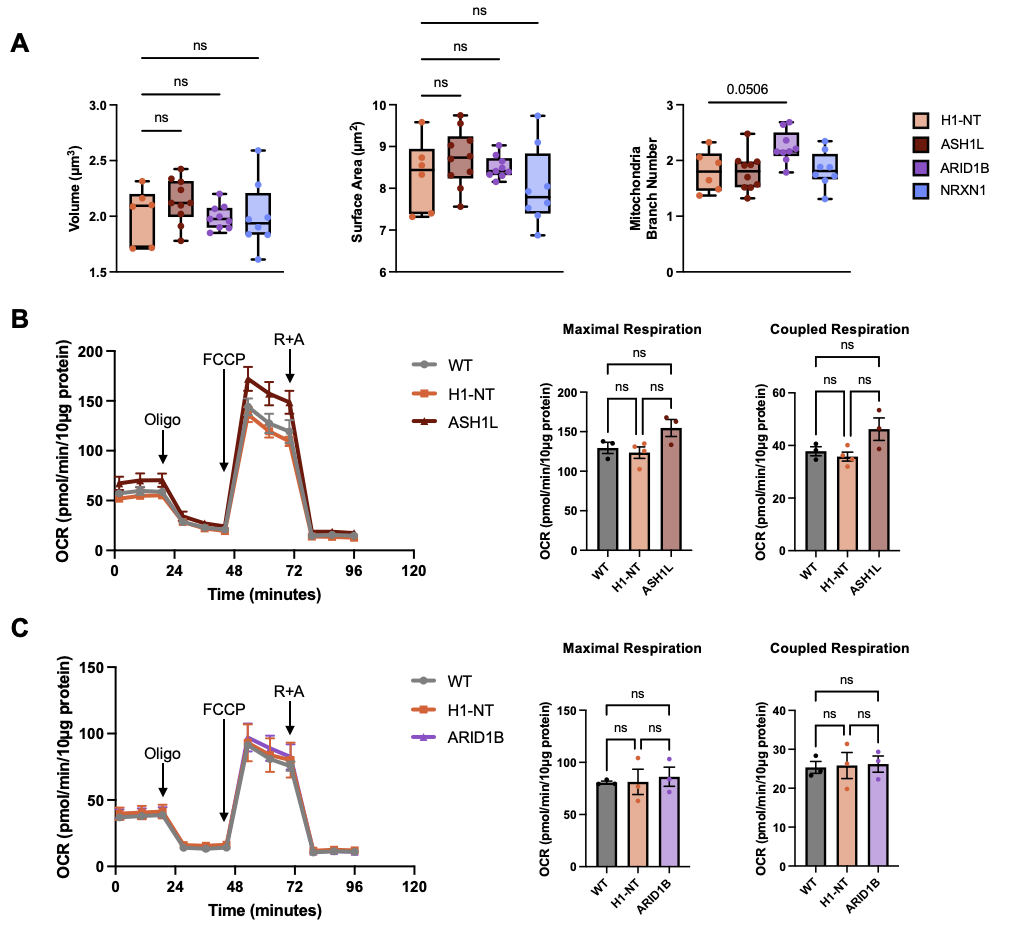
**SI Figure 21. NDD KOs converge on mitochondrial function differently in mature iGLUTs. (A)** High resolution, high-throughput microscopy of mitochondrial morphology of segmented MAP2-positive dendrites labelled for TOMM20-positive mitochondria. Boxplots indicate minimum, maximum and mean values. Each datapoint indicates one well of a 96-well, representing hundreds of μm^2^ of neuronal area and tens of thousands of individual mitochondria (*adjusted p<0.05, ** adjusted p<0.01). **(B–C)** Seahorse assay results for maximal and coupled respiration in ASH1L (B) and ARID1B (C) KO iGLUTs. Oligo: oligomycin; FCCP: carbonyl cyanide 4-(trifluoromethoxy) phenylhydrazone; R+A: rotenone and antimycin A. Data are presented as mean ± SEM. Statistical analysis was performed using one-way ANOVA. Each datapoint represents one well of a 24-well Seahorse assay plate. The experiment was independently replicated twice.

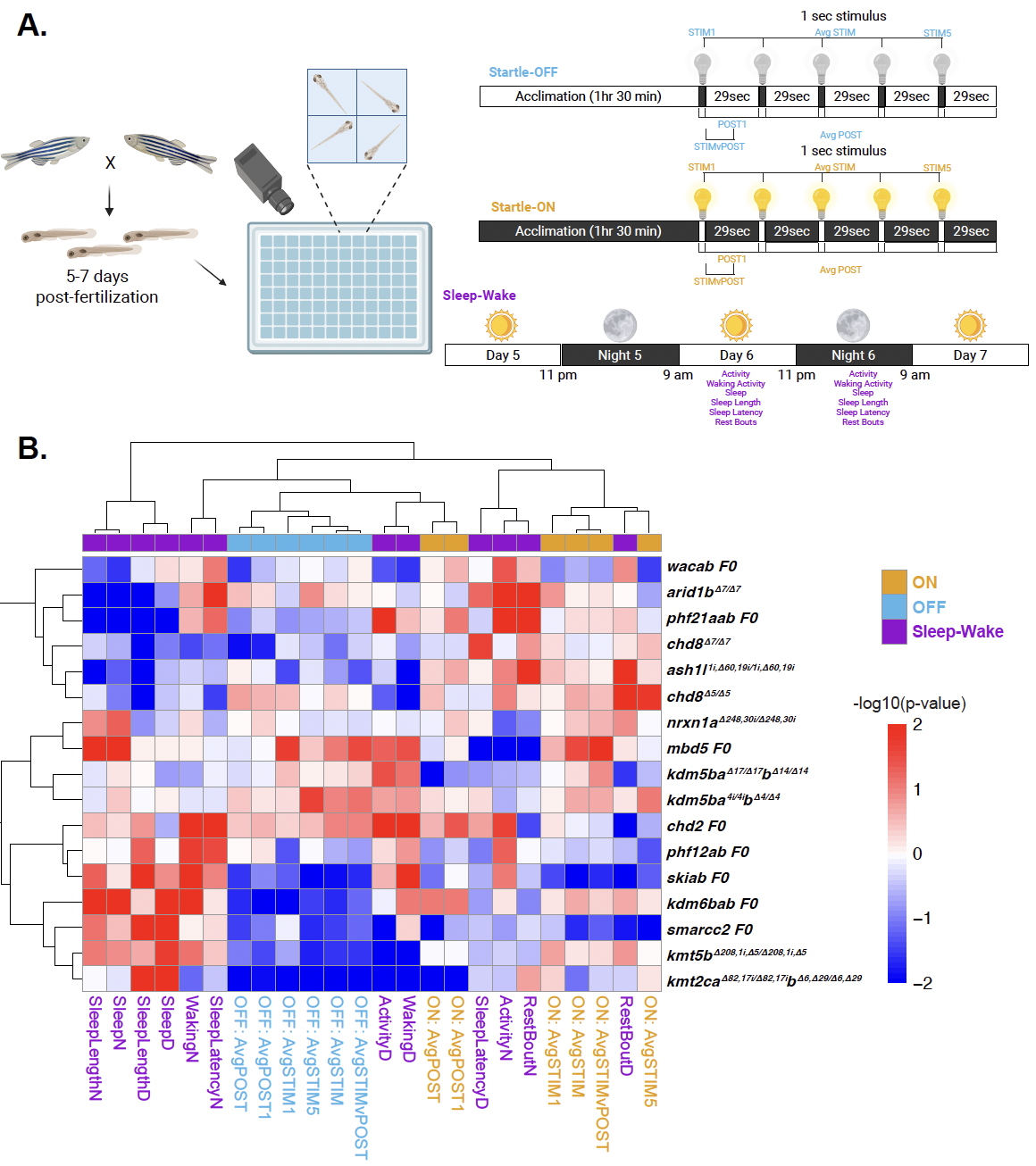
**SI Figure 22. Phenotypic clustering of 15 NDD genes in mutant zebrafish. (A)** Experimental setup and behavioral paradigms for visual-startle and sleep-wake assays. Larval zebrafish mutants are assessed for basic arousal and sensorimotor behaviors between 5-7 days post-fertilization (dpf). Individual larvae are placed in a 96-well plate and behavior is tracked with a video camera within a Zebrabox. *Visual-startle*: Larvae acclimate to ambient light or darkness and are exposed to five 1-second flashes of lights-OFF or lights-ON stimulus at 29-second intervals at 5 dpf. *Sleep-wake:* Larvae are exposed to 14h:10h light:dark cycle in which movement, sleep, and activity parameters are measured over 5-7 dpf. **(B)** Hierarchical clustering of mutant behavioral fingerprints across startle-OFF (light blue), startle-ON (orange) and sleep-wake (purple) parameters. Each box represents the signed -log_10_-transformed p-values from linear mixed models (LMM) comparing behaviors in stable mutant and background-matched wild-type fish and F0 mutant and scrambled control (red, increased in mutant; blue decreased in mutant).

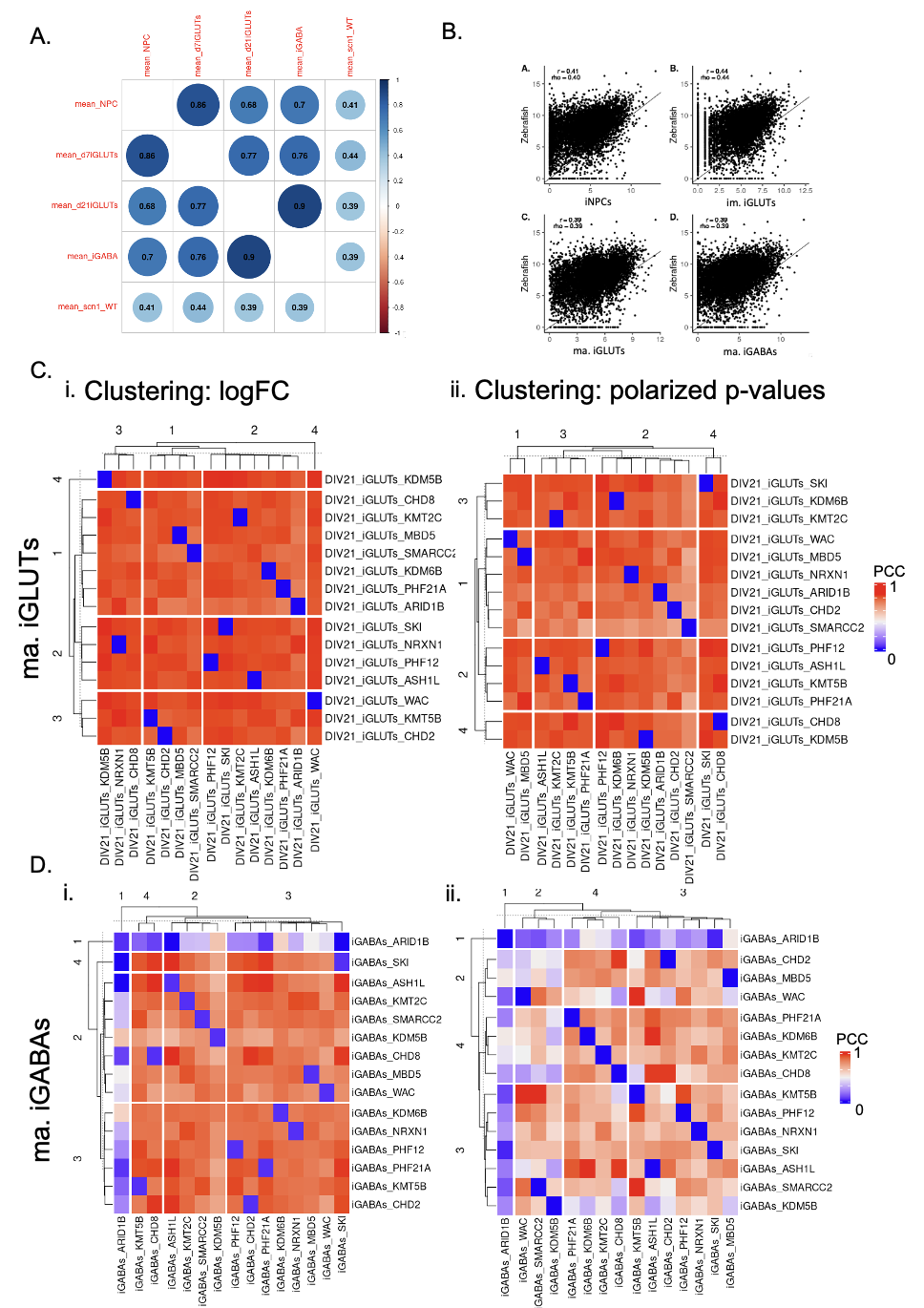
**SI Figure 23. Transcriptomic clustering of 15 NDD genes in mature neurons with paired behavioral data in mutant zebrafish. (A-B)** Expression of non-targeting cells in iNPCs, iGLUTs, and iGABAs significantly correlates with expression of gene homologs and orthologs in the wild-type zebrafish brain. **(C-D**) Heatmap of KO correlations based on logFC **(i)** and polarized –log10 pvalues **(ii)** in mature iGLUTs **(C)** and iGABAs **(D).** Differential gene expression results from individual KOs were filtered for genes with direct zebrafish gene homologs and orthologs using the R package orthogene. Pearson’s correlations across logFC and polarized pvalues (-log10 pvalues multiple by -1 if logFC was below 0) were used to subset KOs into 4 K-means clusters for each cell-type.

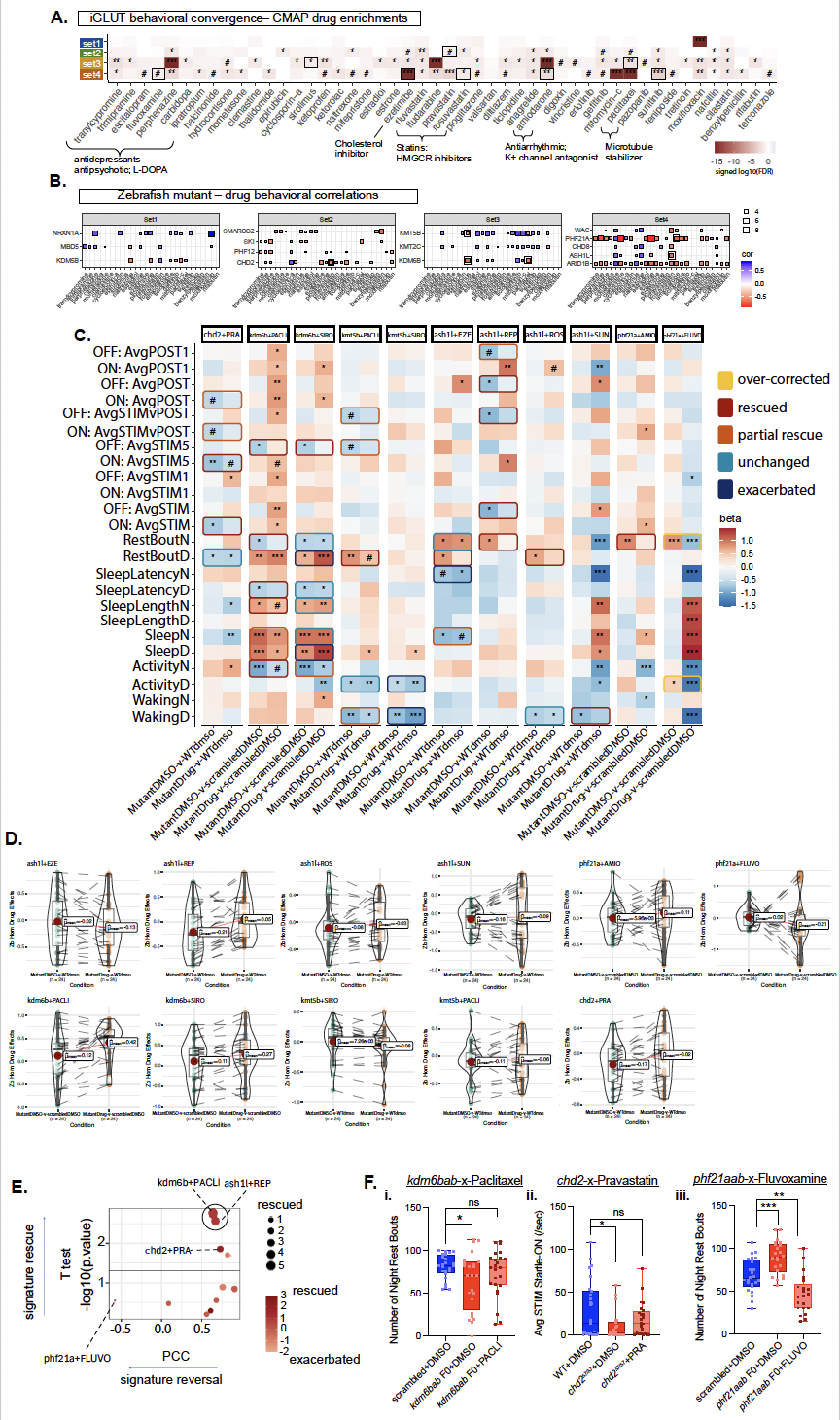

**SI Figure 24.** **Drug reversal of behavioral phenotypes in mutant zebrafish**. **(A)** Drug enrichments using cMAP were calculated for cell-type specific convergence across sets and filtered for drugs included in a screen of 376 compounds for behavioral effects in zebrafish**.** Prediction of drugs targeting these convergent signals in mature neurons identify drug reversers that target shared convergence across the sets (e.g., antipsychotic perphenazine in iGLUTs) and some specific to unique convergence (e.g., valsartan only in Set 3 iGABAs: **SI Data 2**). The heatmap is colored by the polarized (- or + based on direction of enrichment) unadjusted p-values. Significance after multiple testing corrections is indicated by the following: **^#^**nominal p-value<0.05, *****FDR<0.05, ******FDR<0.01, *****FDR<0.001). Notably phenotypic and drug enrichments were only FDR significant in iGLUTs, reflecting that convergent signatures in mature iGLUTs across NDD sets are especially relevant for downstream phenotypes. **(B)** For top predicted drugs from convergence analysis in mature iGLUTs, wild-type zebrafish behavior following exposure to the drug was compared to behavior in individual NDD mutants. **(C)** Heatmap of beta values for each tested behavior between the effect of the mutant alone (homozygous mutant or crispant+DMSO-v-wild-type or scrambled+DMSO) and for the effect of the interaction of the mutant with drug treatment compared to WT vehicle (homozygous mutant or crispant+drug-v-wild-type or scrambled+DMSO). **(D)** Comparison of the magnitude of effect (beta) on behavior between the mutant compared to Mutant+Drug groups for each empirical test. **(E)** Visualization of mutant-x-drug tests, comparing the extent of behavioral signature rescue to signature reversal. **(F)** Representative parameters that were either rescued (i-ii) or over-corrected (iii) in MutantxDrug combinations. (i) Decreased nighttime sleep bouts in *kdm6b* mutants were rescued by paclitaxel. (Unpaired t test with Welch’s correction: crispant+DMSO vs scrambled+DMSO p=0.0289, t=2.291, df=31.12; crispant+PACLI vs scrambled+DMSO p=0.2485) (ii) Decreased response to lights-ON stimulus in *chd2* mutants was rescued by pravastatin. (Unpaired t test with Welch’s correction: *chd2^Δ7/Δ7^*+DMSO vs WT+DMSO p=0.0206, t=2.409, df=40.47; *chd2^Δ7/Δ7^*+PRA vs WT+DMSO p=0.1909) (iii) Nighttime sleep bouts were increased in *phf21a* mutants but significantly decreased beyond wildtype levels with fluvoxamine (Unpaired t test with Welch’s correction crispant+DMSO vs scrambled+DMSO p=0.0004, t=3.808, df=43.86; crispant+FLUVO vs scrambled+DMSO p=0.0016, t=3.392, df=39.59).
